# A minimal model of a forest after neutral theory

**DOI:** 10.64898/2026.10.22.760759

**Authors:** Matthew D. Schwartz, James P. O’Dwyer

## Abstract

Community ecology seeks the mechanisms underlying biodiversity, and simplified models can reveal them. Neutral ecology exemplifies this approach, but it doesn’t distinguish species, limiting its capability for forecasting and prediction. We address this by developing a predictive, minimal model for forest dynamics. We show in a neotropical forest plot that environmental variation couples to the community as a single common effect on all species’ growth rates. Life history acts on that variation, forecasting species’ gains and losses a decade ahead. Finally, immigration, coupled to competition and life history, allows us to predict long-run species abundances. Simple rules thus govern the dynamics of this forest: competition, immigration, and the effect of the environment on growth. Beyond that, species are set apart by their life histories alone.

## Introduction

A central goal of community ecology is to explain the biology of an ecosystem: how species differ and how they interact, and how this underlying biology influences biodiversity outcomes. The simplest theory of a community, neutral biodiversity theory (1, 2), drops species’ individuality altogether. This approximation is predictive: it gives the shape of rarity and abundance well enough to have become a widely-used null model (3–5), and has been tested thoroughly in many forest plots (6–8). What neutral theory cannot do is tell species apart, and so it says nothing about which species will gain or lose individuals, or which will be common in the long run. Forest composition also changes several times faster than neutral drift allows (9–11). Generalizations of neutral models that incorporate environmental stochasticity, a variable factor that multiplies every species’ demographic rates, can restore the pace of change. But these models still cannot say which species will move or in which direction (9–11).

The ingredients that allow us to tell species apart are known. Repeated censuses measure each species’ rates of survival, growth and recruitment directly (12). Built using those rates at the Barro Colorado Island Forest Dynamics plot, the life history model of Jops et al. (13) predicts changes in composition that are faster than drift, but is silent on which species will rise and which will fall at any given moment in time. We derive a minimal model with four ingredients, and test it on the 50-ha Barro Colorado Island (BCI) plot, from seven census intervals spanning 33 years. Each species has a measured life history (14, 15), competes for space, can immigrate from outside the community, and experiences environmental stochasticity. No parameter is tuned per species to improve predictions. With these four ingredients, rates measured in the censuses up to a given date forecast the gains and losses of large trees a decade (two censuses) ahead for each of 84 species better than the species’ own historical trends. When immigrants arrive according to a competition-colonization trade-off (16–18), the same model also ranks the species’ long-run abundances in the order the plot shows today. Because so little is fitted, our results are interpretable. The dynamics of the stand as a whole are governed by competition for space, by immigration and by the good and bad years, and a species’ fate within it follows from its own life history.

**Figure 1.**
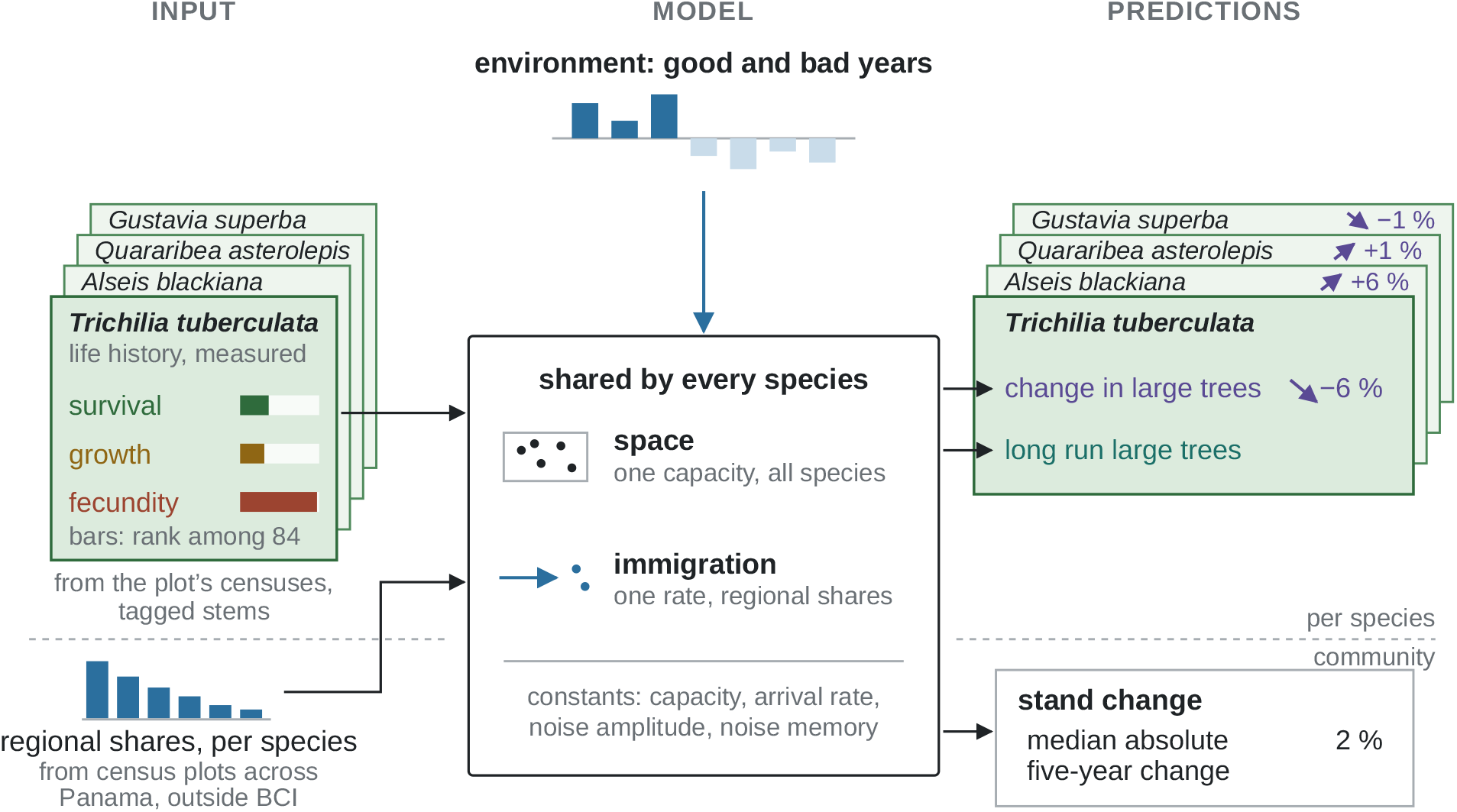
The model as a pipeline. Input, left: each species’ measured life history, in the stage-structured form of its transition matrix, from the plot’s own censuses of tagged stems, summarized by three properties read from that matrix, survival (of its large trees per census interval), growth (the promotion probability out of its sapling classes per interval) and fecundity (new seedlings per large tree per interval), each bar the species’ rank among the 84 species; and each species’ share of the surrounding forest, from census plots across Panama. Model, center: two rules shared by every species, competition for space through one capacity and immigration at one rate in regional proportions, driven from outside by the environment, one factor per census interval that is the same for every species; four constants in all and none per species. Predictions, right: for each species the direction and size of its change in large trees over one census interval, as the model’s forecast from the 1995 census for 1995 to 2000 in percent of its large trees, and, without a number, its long-run abundance of large trees, and for the stand as a whole the median absolute five-year change of its large-tree total, in percent of that total.

## Model

The main census of the 50-ha Barro Colorado Island Forest Dynamics plot records every stem of at least 1 cm diameter (dbh) every five years, and a separate seedling census counts smaller stems in plots of 1 m^2^ (19). Following Jops et al. (13), we sort the censused stems into diameter classes 2 to 8 whose lower edges double from one class to the next, and use class 1 for seedlings below the census diameter threshold. We use ‘censused stems’ to mean every stem of class 2 and above, and we call the stems of classes 5 to 8 (at least 79.4 mm dbh) the large trees. Each species *s* is a stage-structured population (14, 15) with a count of stems in in each of the eight classes, its life stages. Each gets an 8 *×* 8 matrix *A*^*s*^, built by following individual stems from one census to the next, the tagged stems of the main census over its seven intervals from 1982 to 2015 and the seedlings of the seedling census in class 1. Each column of *A*^*s*^ is a life stage, and its entries are the chances that a stem in that stage survives and stays, advances or drops back over one five-year interval, detailed in Supplementary Text S2. The first row characterizes fecundity, holding the seedlings each stage produces, scaled so that the species’ population growth rate is one.

The model shares three processes from neutral theory: a fixed number of places, a lottery that fills vacancies without regard to species, and immigration from a regional pool. We run the dynamics stem by stem, so that in each census interval every stem survives or dies and a survivor stays, advances or drops back, by Poisson draws at its species’ specific rates. We impose competition for space via one carrying capacity, fixed at the observed mean number of censused stems in the plot. When a censused stem dies, a seedling takes its place, drawn at random from that interval’s seedlings ready for promotion (20). Candidate seedlings in excess of the vacancies remain seedlings, and the seedling class is uncapped, with its size an outcome of the model. Seeds arrive from outside the plot at a fixed total rate of *M* per interval, divided among species in fixed proportions *w*^*s*^ that are built from each species’ share *p*^*s*^ of the stems in census plots across Panama. We consider two model variants. In the regional-pool model, the default of neutral theory, seed arrives in proportion to regional abundance, *w*^*s*^ = *p*^*s*^ (21). In the trade-off model, built on a competition–colonization trade-off (16–18), species that do well locally due to their life history strategies colonize poorly, imposed by setting immigration inversely proportional to the large trees produced per arriving seed (22, 23).

Some environmental variation is needed, since without it the model’s forest would be steadier than the census shows (Fig. 2). But there are many ways to add it, since every life stage of every species could respond to the environment in its own way. We ask how few of these responses the predictions need. Nearly every environmental term is statistically detectable in the transition data from one census to the next, and a Bayesian information criterion would keep almost all of them, at the cost of simplicity and interpretability. We instead keep a term only if it is large enough to change one of our predictions: a species’ change over a decade, the long-run composition, or the change in the stand’s large-tree total. Only one effect passes this test, a factor shared by all species. Only when acting on sapling promotion, growth out of classes 2 to 4, does this factor make the recruitment of large trees rise and fall with the stand total as observed, while mortality’s common component is weaker and out of step with growth’s. The noise term we keep is sufficient to generate fluctuations as large as the census shows, and the terms we drop are not: without environmental noise the large-tree total changes over five years 11 times less than in the census, the shared sapling factor alone brings that change and the variability of recruitment close to the census values, and each species’ own mortality variation alone gives only about half the census’s change in the total and a tenth to a sixth of its variability in recruitment. Started from observed initial conditions, the same holds species by species (Supplementary Text S5 and S7).

**Figure 2.**
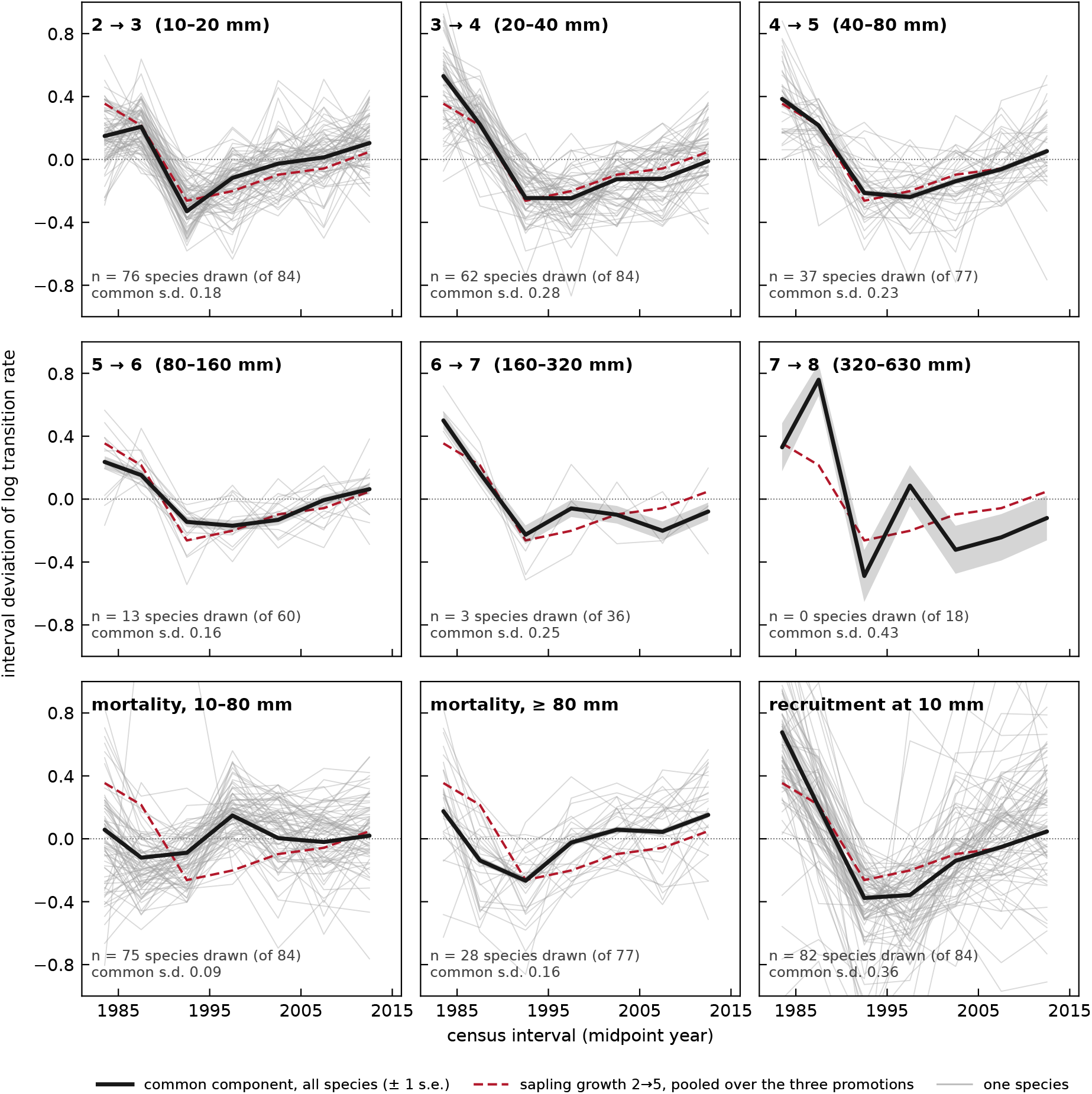
Between-census variation of nine transition rates for the 84 modeled species at Barro Colorado, 1982–2015, from the census data. The top row shows sapling promotions (classes 2 *→* 3, 3*→* 4, 4 *→* 5), the middle row promotions among large trees (5 *→* 6, 6 *→* 7, 7 *→* 8), and the bottom row mortality below and above 79.4 mm dbh together with recruitment across 10 mm per standing stem. Gray lines, single species’ log-rate deviations from their own seven-interval means, for species with an expected count of at least twenty events per interval (*n* in each panel, none for 7 *→* 8). The bold line and band show the component common to all species with its standard error, and the numbers give the standard deviation of that component. The dashed line is the three sapling promotions pooled into one rate (the pooled sapling path, standard deviation 0.22), from which we estimate the noise amplitude *σ*^*g*^, the standard deviation of the factor (Materials and Methods, S1). One vertical scale throughout. Sapling growth varies between intervals in common across species, promotions among large trees follow it, and the common component of mortality is weaker and not synchronized with growth (species-level statistics in Supplementary Text S5).

## Results

In summary, the model relies on each species’ life history via the matrix *A*^*s*^, and immigration is determined by its regional share *p*^*s*^. In addition, the model has four constants, all shared by every species: the carrying capacity, the arrival rate *M*, the amplitude of the environmental variation and its memory, the correlation between environmental conditions over time. We now test the model’s predictions on two timescales, each species’ change in large trees over the next decade and the composition of the forest in the long run.

### Each species’ next decade

We first ask whether the model can forecast which species will gain or lose large trees over the next decade, given only data known up to a given date. We compute each species’ matrix from the censuses of 1982 to the start date of 1995, 2000 or 2005, initialize the model from each species’ observed stage vector, and predict its large-tree count five and ten years ahead. No future data enters the rates. The three forecasts when pooled rank the species’ observed changes with a rank correlation of 0.72 (90% CI 0.61–0.81) five years ahead and 0.70 ten years ahead, and they get the direction of change right for 75% and 72% of the species whose count changed. Had we used the matrices estimated from all the censuses of 1982–2015 (i.e. including the intervals being forecast), the rank correlation on the same species and dates would rise to 0.84 and 0.86 and the direction of change correct at 84% and 83%, for 5 and 10 years respectively (Fig. 3). Since the capacity, the lottery, and immigration act on seedlings which cannot grow into large trees within a decade, these elements of the model do not much affect the forecast. Indeed, turning all three off and applying the forecast’s matrices directly to each species’ stems gives little change: the rank correlation still comes out as 0.72 and 0.70 for the two intervals.

**Figure 3.**
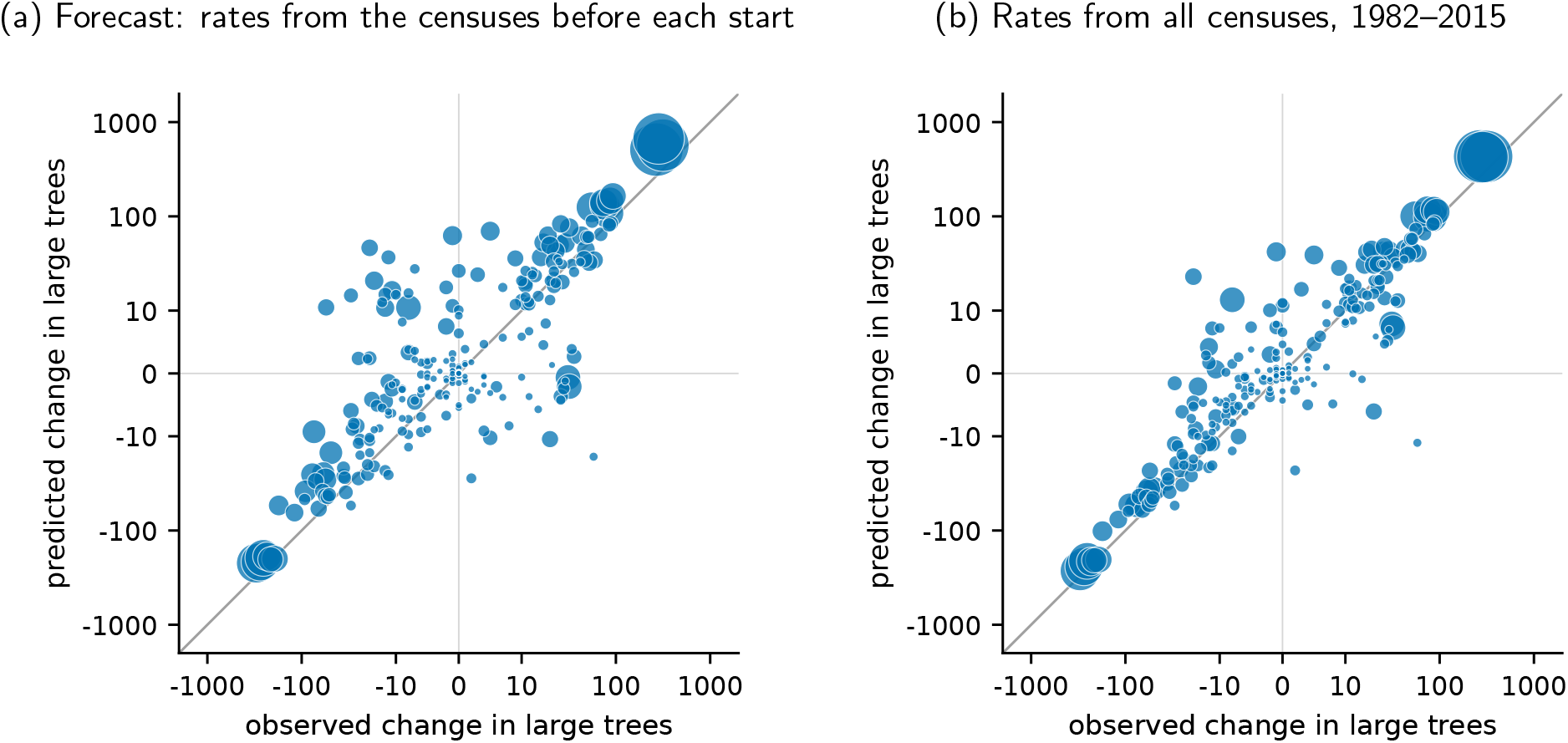
Predicted versus observed change in each species’ large trees (stems *≥* 79.4 mm dbh, classes 5–8) two censuses ahead at Barro Colorado, one point per species-start pair (one species at one starting census), for the 252 pairs that start in 1995, 2000 and 2005. Symbol area scales with the species’ large-tree abundance at the start, the line is the identity, and the axes are logarithmic away from zero. (a) At each start, matrices from all the censuses before that start, a forecast because none of the predicted intervals enters the rate estimates. In absolute numbers these forecasts are biased upward. (b) The same pairs with matrices from all censuses of 1982–2015, which include the predicted intervals. Predictions are each species’ matrix applied twice to its observed stage vector (Materials and Methods, S1), and observations are census counts. Rank correlations are 0.70 (a) and 0.86 (b). Sign, error and bias, the comparison with the long-term trend and one-census panels are in Supplementary Text S3.

A neutral model cannot make such a forecast, since drift carries no direction. Neither can a model with environmental variance but no life histories, which does not single out which species will change (9–11). The simplest alternative with a direction, each species’ previous trend (either across all censuses or just the previous interval), is outperformed by our model, and long-term trends cannot ever anticipate a reversal in direction. Species specificity is also important; our forecast needs each species’ own life history to work, and the mean of all species’ matrices, applied to each species’ stage vector, ranks the changes at only 0.24 and 0.26 out of sample and at 0.02 and 0.01 in sample. None of 1,000 random reassignments of the matrices among species reaches the species-specific prediction performance (Supplementary Text S3).

### Long-run composition

Turning to the long-run forecast of the model, where the environment and immigration play an important role, we begin with exact results in the mean-field approximation. Holding the environmental factor at one and replacing every random draw by its average, the model settles to a stationary state that we can write down in closed form. In the mean-field approximation, competition enters through one number common to all species, the waiting fraction *d*, the fraction of seedlings ready for promotion that find no vacancy in an interval. In our model the stationary number of large trees of species *s* is

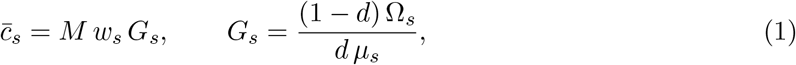

where *µ*^*s*^ is its seedling mortality per interval, Ω^*s*^ the number of large trees per seedling in its stable stage structure, and *G*^*s*^, the local gain, the number of large trees standing per arriving seed. The waiting fraction holds every species slightly below replacement, since cutting its seedling promotions lowers its growth rate below one, so that without immigration every species would slowly decline and immigration sustains it. The two model variants give distinct long-run compositions. In the regional-pool model, *w*^*s*^ = *p*^*s*^, and the large-tree composition is proportional to *p*^*s*^*G*^*s*^, independent of *M* and *d*. In the trade-off model, arrivals are weighted inversely to the gain, *w*^*s*^ ∝ *p*^*s*^/*G*^*s*^, and the long-run composition is the regional composition *p*^*s*^ itself.

A forest need not be near its stationary state. Linearized at the fixed point, our model predicts the pace of approach to steady state, set by the decay of the species held least far below replacement, with timescale *T*^*s*^/(*d q^s^*), its generation time *T*^*s*^ (the mean age of the parents of its seeds in the stable stage structure) divided by the waiting fraction *d* and by *q*^*s*^, the chance that a seedling dies unpromoted. These timescales turn out to be 2,900 years under the regional-pool model and 19,000 under the trade-off model, under which fewer candidates wait and *d* is smaller. Simulations started from unrelated compositions become stationary (counts stop changing and the starts agree species by species) after 10,000 to 12,000 years under the regional-pool model and, under the trade-off model, after up to 16,000 years from above the capacity and up to 42,000 from an empty plot. Thus, the timescale for the model to assemble a steady-state community at Barro Colorado Island is one to four times the timescale of the Holocene, with that factor dependent on the immigration process and initial conditions. We know neither: the regional pool was itself assembling over that interval, so we cannot take immigration as constant through it, and the plot’s composition at any earlier time is unrecorded.

We can’t establish that the forest has arrived at a steady state, but we can use our 33 years of censuses to compare the present composition with the predicted stationary state, simulated with environmental noise. We find that the observed order of species broadly matches the stationary order under the regional-pool model (Spearman correlation 0.675, Fig. 4a), but the regional-pool model gets the levels wrong: in absolute numbers its identity-line *R*^2^ on log abundance is -0.726. Our trade-off model introduces a plausible hypothesis that has the potential to fix this mismatch; with no more fitted parameters than the regional-pool model, our trade-off model does better on every count (Fig. 4b, c): it keeps the rank order (Spearman 0.714), brings absolute abundances close to the census (*R*^2^ = +0.362) and the large-tree total to 0.84 times the observed, and its species abundance distribution is closer to the census than the regional-pool model’s (Kolmogorov distance 0.215 against 0.278) and about as close as a dispersal-limited neutral model (2, 3) fitted to the same 79 species (0.18).

**Figure 4.**
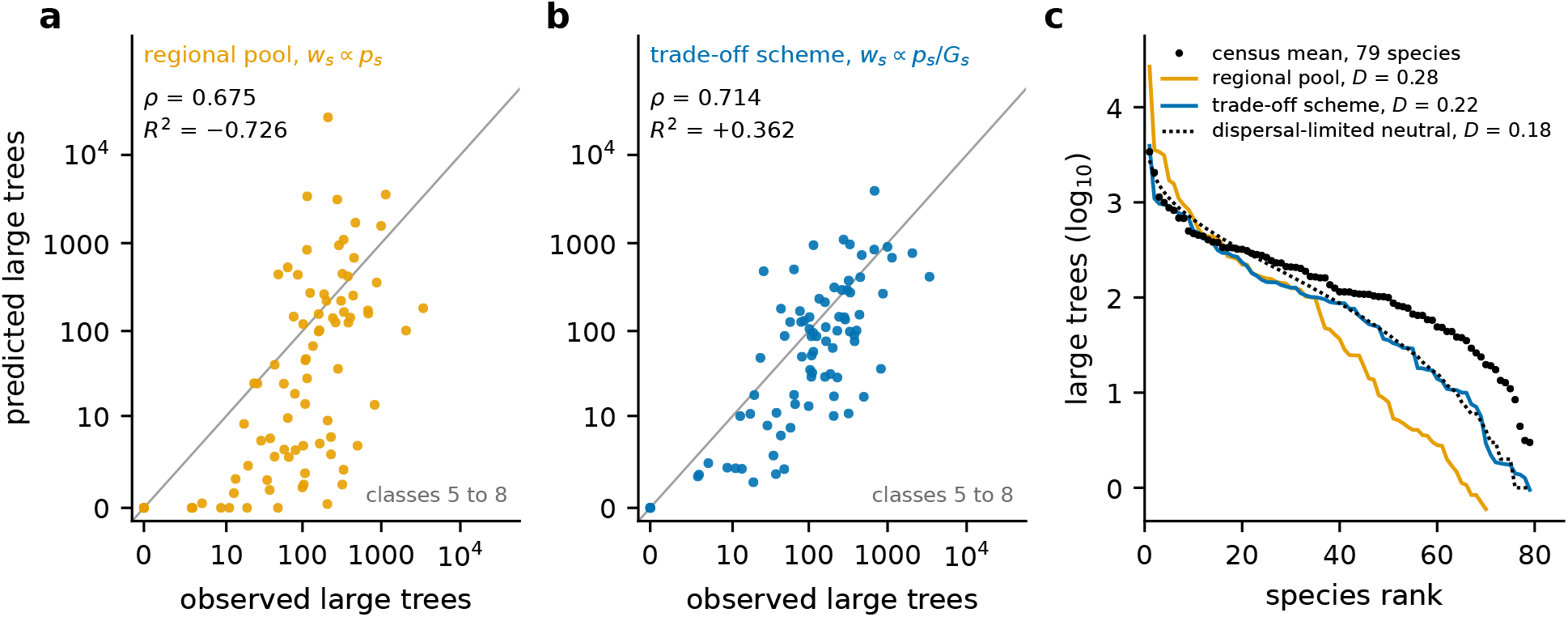
Stationary large-tree abundances (stems of at least 79.4 mm dbh, classes 5–8) under the two model variants, from the simulations with the environmental noise on, run to certified stationarity (72 replicates per variant, 120,000 model years each, species’ means over the last five windows), compared with the 1982–2015 Barro Colorado censuses. (a) Regional-pool model, arrivals weighted by regional share *p*^*s*^: predicted versus observed mean abundance for the 84 species (log axes, identity line). Spearman *ρ* = 0.675, identity-line *R*^2^ = *−*0.726 on log abundance (negative when the observed mean alone fits better). (b) Trade-off model, arrivals weighted by *p*^*s*^/*G*^*s*^, inversely to the local gain *G*^*s*^: *ρ* = 0.714, *R*^2^ = +0.362. (c) Ranked log_10_ abundance of the 79 species reaching the large-tree classes: census (points), both model variants (lines), Hubbell’s dispersal-limited neutral model fitted, with its two parameters free, to the census abundance distribution of the same species (dotted, with fundamental biodiversity number *θ* = 21 and immigration probability *m* = 0.0085). The neutral fit is separate from the immigrant fraction 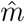behind the arrival rate (Materials and Methods, S1) and uses neither the regional shares nor species’ identities. *D*, Kolmogorov distance from the census. The closed form of Eq. (1), the mean-field reference, gives the same order at a higher level (Supplementary Text S4 and S6).

Aggregating trees across all species, the overall trend of total abundance across the censused classes is captured well, but specific size classes do deviate from observed data. Large-tree totals under the trade-off model are 0.84 times the observed, while we underpredict the saplings, at 0.65 of the census (Supplementary Text S4 and S6). Finally, the seedling density was left unconstrained in the model, and is also a prediction that can be tested. Our model overpredicts its size: the seedling prediction is 1.7 times the seedling-census mean. This excess holds at every arrival rate tried, ranging from half to four times our original value of *M*, and other alternative model formulations of the system capacity can reduce, but do not remove this excess entirely (Supplementary Text S6 and S7). We speculate that these deviations may in part reflect a historical, successional process. About half of the model’s predicted number of seedlings belong to a single species, *Gustavia superba*, a long-lived pioneer that can dominate secondary forest (24–26). If this species is declining on the plot, our model likely overestimates its fecundity, leading to an excess of predicted seedlings, and with that species left out of the count, the model’s seedling pool is 0.82 times the census.

## Discussion

Our minimal model of a forest predicts outcomes on two timescales, incorporating stochastic community dynamics, stage-structured demography, and environmental variability as a shared effect, common across all species. This parsimonious treatment captures most of the variability in the recent history of the BCI forest plot. From early census data we predict each named species’ change in large trees a decade ahead, while in the long run the model recovers the census order and typical magnitude of species’ abundances. The model draws upon several existing ingredients (2, 9, 13), but the result is more than the sum of the parts.

We used observed transition data, rather than a modeling choice, to place the environmental coupling on sapling growth. We do not identify which environmental variable drives this factor, though the data strongly constrain what it *could* be. Canopy opening, which droughts and storms synchronize across a plot (27–29), would act on sapling growth as the factor does, though light-demanding species follow the factor no more strongly than shade-tolerant ones. Of the climatic drivers (light, temperature and rainfall (30–33)), no series we examined tracks the factor beyond chance over seven intervals. Measured life histories and a single common effect on growth acting through them thus account for most of the faster-than-drift change observed in this forest plot, and species-specific couplings add little in return for the cost of generating a much more highly-parametrized model.

Summed across all species, the model predictions approximately match stem counts across size classes, but deviate from observed densities the most in predictions for total numbers of sapling and seedling stems. In the long run, how common the model makes a species depends on how common that species is across the region, and we know that only from 28 census plots elsewhere in Panama. The species whose abundances the model gets most wrong, those that are common on the regional plots and scarce at Barro Colorado, or the reverse, drive these departures. This suggests the need to refine which regional plots inform the immigration process, and the specific species driving those deviations indicate that successional dynamics may be an important next step in the development of this modeling approach.

Our parametrization is particular to Barro Colorado, and any other plot would need at least two censuses of tagged stems for the projection matrices, several intervals for the environmental noise, and a regional sample for long-term abundances. Other forests experiencing site-wide, significant disturbance much more often than about once in a few thousand years (e.g. (34)), would likely not be near the stationary state. Our results do speak though in this forest to an old question: whether a plant community is an integrated whole (35) or a set of species that merely share a place, each answering the environment on its own (36–38). At Barro Colorado we show something coherent shared by all species in their response to the environment, while the idiosyncrasy in how species’ abundances then move is mediated by their life histories.

## Acknowledgments

We thank R. Condit, S. P. Hubbell, R. B. Foster, R. Pérez, S. Aguilar and S. Lao for making the Barro Colorado Island (BCI) and Panama plot censuses public, L. S. Comita and her coauthors for releasing the seedling census, and K. Jops for archiving the demographic matrices. The BCI forest dynamics research project was made possible by National Science Foundation grants to Stephen P. Hubbell: DEB-0640386, DEB-0425651, DEB-0346488, DEB-0129874, DEB-00753102, DEB-9909347, DEB-9615226, DEB-9615226, DEB-9405933, DEB-9221033, DEB-9100058, DEB-8906869, DEB-8605042, DEB-8206992, DEB-7922197, support from the Forest Global Earth Observatory (ForestGEO), the Smithsonian Tropical Research Institute, the John D. and Catherine T. MacArthur Foundation, the Mellon Foundation, the Small World Institute Fund, and numerous private individuals, and through the hard work of over 100 people from 10 countries over the past three decades. The plot project is part of the Forest Global Earth Observatory (ForestGEO), a global network of large-scale demographic tree plots. We acknowledge the support of the Forest Global Earth Observatory of the Smithsonian Tropical Research Institute for the BCI plot.

## Author contributions

M.D.S. brought exact methods from mathematical physics to bear on neutral theory and the dynamics of forest communities; J.P.O’D. supplied the ecological framing, the demographic structure of the model and the published life tables on which it is built. Together they conceived the study, decided its questions and tests, and wrote the paper. Modeling, simulation and data analysis were carried out by M.D.S. with an AI research assistant (Claude Fable 5.1, Anthropic). Claude acquired the data, wrote and ran the code, kept the record of every analysis, and drafted text and figures for the authors to revise. The authors take responsibility for all results and their interpretation.

## AI use

Claude Fable 5.1 (Anthropic) was used for writing or drafting manuscript content: it produced an initial draft of the manuscript, which the authors then almost entirely rewrote. It was also used for editing and formatting the visualizations of research data and results (Figs. 2 to 4 and the Supplementary figures), which its scripts draw from the derived tables. Its use for writing code and data analysis is described in the Supplementary Materials (Materials and Methods, S1).

## Data and materials availability

All data analyzed here are public, and we used them under the licenses stated with each source. The 1982–2015 censuses of the 50-ha BCI plot are the complete-plot release archived on Dryad (39, doi:10.15146/5xcp-0d46, CC0); the plot and its census methods are described by Condit (40) and Hubbell et al. (41). The seedling and small-sapling census of the same plot is the Dryad release of Comita et al. (42, doi:10.5061/dryad.05qfttf85, CC0), described by Comita et al. (19). We took each species’ stage-structured projection matrix from the Zenodo archive of Jops and O’Dwyer (43, doi:10.5281/zenodo.14563651, CC BY 4.0) that accompanies Jops et al. (13), and recounted its seedling column from the seedling census. We built the regional species pool from the Panama plot-network tables of Condit et al. (44, Smithsonian, doi:10.5479/data.stri.2016.0622, CC BY 4.0); the network’s current archive is on Dryad (45, doi:10.15146/mdpr-pm59, CC0). Species’ light responses, functional traits and the climate series, which we use only for exploratory comparisons with the environmental noise, come from further public datasets listed with their DOIs and licenses in Supplementary Text S2. All of our code, the stochastic simulation engine, the closed-form solver, the estimation of the environmental noise and every analysis script behind the numbers, tables and figures reported here and in the Supplementary Materials, is held with its run records in the project repository and will be archived with a README and a DOI on publication, together with the derived tables from which all results can be reproduced. The archive also contains the projection matrices that we estimated for the out-of-sample forecasts from the censuses before each start, and the code that computes those matrices from the census data.

## Supplementary Materials

Materials and Methods (section S1)

Supplementary Text, sections S2 to S7: S2 Barro Colorado data, species and sources; S3 the decade-ahead forecast in detail; S4 the stationary state and long-run composition; S5 environmental noise estimated from the transition data; S6 model variants and sensitivity of results; S7 fluctuation predictions

Figs. S1 to S9

Tables S1 to S27

References

