## Supplemental Materials for "A minimal model of a forest after neutral theory"

Matthew D. Schwartz

Department of Physics, Harvard University

James P. O'Dwyer

Department of Plant Biology, University of Illinois at Urbana-Champaign

### S1 Materials and Methods

The BCI censuses record every free-standing stem of at least 1 cm diameter at breast height (dbh) every five years (Condit et al., 2012, 2017, 2019). We use the censuses of 1982–2015 (seven intervals) and, of the 90 species with published stage-structured matrices (Jops et al., 2025; Jops and O'Dwyer, 2024), follow the 84 that gained or lost at least one tree of 10 cm dbh, the conventional tree threshold, over these censuses (our large-tree floor of 79.4 mm is the class edge nearest that convention), keeping the matrices' eight stages and their normalization to growth rate one. We recount the seedling column of each matrix from the seedling census (Comita et al., 2023a,b) and rescale each fecundity row so that the growth rate stays one—in effect, introducing an assumption of equal fitness, shared across all species. Adding the nine further species of the plot whose rates are as well constrained as those of the 84 leaves the stationary results for the 84 within their 90% ranges (Section S6). Four community-level constants complete the model, none per species: the capacity (200,721 censused stems), the arrival rate $M$ (26,499 immigrant seeds per census interval), the immigrant fraction $\hat{m} = 0.026$ times a nominal seed production of 1.01 million seeds per interval, the order the fecundity rows give, with $\hat{m}$ the value at which Hubbell's neutral model best reproduces the plot's census-to-census turnover of trees of at least 10 cm dbh, and the amplitude and memory of the environmental noise. We take regional shares from the census plots across Panama (Condit et al., 2022, 2016) with BCI and its six neighbors excluded. Simulating the environmental noise at the census dates, we reproduce the spread of the pooled sapling path in Fig. 2 of the main text at an amplitude ($\sigma_g$) of 0.32 (90% CI 0.17–0.84). Since seven serially correlated intervals cannot constrain the memory, we assume an exponentially decaying memory with a correlation time of 18 years, a correlation of 0.76 between consecutive five-year intervals. We forecast each species' large-tree count one or two censuses ahead from its matrix and observed stage vector alone (Fig. 3 of the main text). Full methods are in Sections S2 to S7.

**AI use.** Writing code and data analysis were performed largely by AI, Claude Fable 5.1 (Anthropic): it acquired and processed the public data sets, wrote and edited the code for the simulations, the closed-form solver, the estimation of the environmental noise and every analysis behind the numbers, tables and figures, and ran that code. No data were generated. The authors provided all decisions on which analyses to perform and which methodology to use, guidance on model construction, prioritization and scientific direction, and checked every result against the run records. The same AI suggested text for the manuscript and produced the figures; the final text was written by the authors.

| class $k$ | 2 | 3 | 4 | 5 | 6 | 7 | 8 |
| --- | --- | --- | --- | --- | --- | --- | --- |
| lower edge (mm) | 10 | 20.0 | 39.8 | 79.4 | 158.5 | 316.2 | 631.0 |

| stems | population |
| --- | --- |
| 201,000 | the censused stand: classes 2–8 (stems of at least 10 mm, the 1 cm census threshold), the 84 modeled species, mean over the censuses of 1982–2010 at which stage vectors are built; the model’s capacity, the fixed total of censused stems, is this mean unrounded |
| 21,700 | large trees: classes 5–8 (stems of at least 79.4 mm), the 84 species, mean over the same censuses |
| 21,000–22,200 | large trees, the range of the per-census totals over the same censuses |
| 14,900–35,300 | inferred flow of stems from class 1 into class 2 per census interval over 1982–2010 (mean 22,300) |

Table S1: **Stage classes and stem populations at Barro Colorado.** Top: lower diameter edges of the model’s stage classes. Class 1, the seedling pool, holds stems below 10 mm. Classes 2 through 8, the censused classes, are log-spaced with lower edges $10 \cdot 10^{0.3(k-2)}$ mm, and class 8 is unbounded above. Classes 5 through 8 are the large trees, whose abundances the model’s predictions are compared with. Bottom: the stem populations used in the main text and in this supplement, rounded.

### S2 Barro Colorado data, species and sources

Jops et al. (2025) published projection matrices for 90 tree species of the 50-ha plot on Barro Colorado Island (BCI), in the Panama Canal (Jops and O’Dwyer, 2024). In the BCI censuses of 1982–2015 (Condit et al., 2019), 255 species have a live tree of at least 10 cm dbh at one census or more. We model the 84 species on both lists (their species codes are listed at the end of this section). The six species with a published matrix but no tree of that size are *Anaxagorea panamensis*, *Herrania purpurea*, *Miconia nervosa*, *Ouratea lucens*, *Palicourea guianensis* and *Psychotria marginata*, all understory shrubs and treelets. Each is abundant on the plot, with at least 251 censused stems at every census, but none had a live stem of 10 cm dbh at any census, and their largest stems measure 5 to 9.9 cm. Their matrices, fitted on smaller stems, are not used. Table S1 gives the lower diameter edges of the stage classes and the stem populations used throughout. We call the stems of classes 5 to 8 (at least 79.4 mm dbh) the large trees.

The immigration weights come from the network of census plots across Panama (Condit et al., 2022, 2016), with BCI itself and the six plots on or beside the island excluded. A species’ regional share $p_s$ is its share of the reproductive-size stems summed over the remaining plots. The nine modeled species absent from those plots are given the smallest share of any sampled species. Section S4 gives the stationary composition with them at zero weight instead. Table S2 lists the 28 remaining plots, with their locations, areas and the censuses used.

The rates of the seedling class (class 1) are estimated from the BCI seedling census (Comita et al., 2023a,b), as described below. It covers every free-standing woody stem at least 20 cm tall and under 1 cm dbh in 19,313 plots of 1 m<sup>2</sup>, at fourteen censuses from 2001 to 2018. Scaled to the 50-ha plot, the modeled species averaged 1.12 million such stems (every record of status A, the status the census gives a seedling found alive, whose dbh is not recorded or is under 10 mm, whatever its codes) over those censuses (0.92 to 1.37 million at single censuses), 67 to 75% of the live seedlings of the 413 species found. The seedling column is estimated on the stems of the seedling census, so class 1 of the model holds the population that the census follows: stems tagged once they reach

| plot | locality | latitude | longitude | area (ha) | census years used |
| --- | --- | --- | --- | --- | --- |
| cocoli | Cocoli | 8.98641 | −79.618 | 4 | 1994, 1997, 1998 |
| gigante1 | Gigante Peninsula | 9.09919 | −79.854 | 1 | 1999, 2000, 2003, 2008 |
| gigante2 | Gigante Peninsula | 9.10353 | −79.8536 | 1 | 1999, 2000, 2003, 2008, 2010, 2011 |
| lp1 | Las Pavas | 9.1247 | −79.9048 | 1 | 2001, 2008 |
| lp2 | Las Pavas | 9.12526 | −79.8853 | 1 | 2001, 2008 |
| p01 | Fort Sherman | 9.33253 | −79.9541 | 1 | 1996, 2012 |
| p02 | Fort Sherman | 9.32296 | −79.9621 | 1 | 1996, 2012 |
| p03 | Fort Sherman | 9.26033 | −79.956 | 1 | 1996, 2012 |
| p04 | Fort Sherman | 9.25802 | −79.9529 | 1 | 1996, 2011 |
| p05 | Pipeline Road | 9.15682 | −79.7517 | 1 | 1996, 2010 |
| p06 | Pipeline Road | 9.1565 | −79.7442 | 1 | 1996, 2010, 2011 |
| p07 | Pipeline Road | 9.16093 | −79.7427 | 1 | 1996, 2010 |
| p08 | Pipeline Road | 9.1683 | −79.7464 | 1 | 1996, 2008 |
| p09 | Pipeline Road | 9.16878 | −79.7407 | 1 | 1996, 2008, 2011 |
| p15 | Pipeline Road | 9.16176 | −79.7453 | 1 | 1996, 2010 |
| p16 | Pipeline Road | 9.14674 | −79.7125 | 1 | 1996, 2010, 2011 |
| p17 | Pipeline Road | 9.15181 | −79.7158 | 1 | 1996, 2012 |
| p19 | Pipeline Road | 9.19917 | −79.7741 | 1 | 1996, 2008, 2011 |
| p20 | Pipeline Road | 9.19028 | −79.7643 | 1 | 1997, 2008 |
| p21 | Gamboa | 9.14149 | −79.6935 | 1 | 1997, 2012 |
| p22 | Gamboa | 9.14787 | −79.6931 | 1 | 1997, 2012 |
| p23 | Gamboa | 9.1217 | −79.6731 | 1 | 1997, 2012 |
| p24 | Gamboa | 9.12372 | −79.6767 | 1 | 1997, 2012 |
| p25 | Laguna | 9.07852 | −79.7989 | 1 | 1997, 2008 |
| p26 | Laguna | 9.07646 | −79.7869 | 1 | 1997, 2009, 2012 |
| p27 | Cruces | 9.08048 | −79.6451 | 1 | 1997, 2008 |
| p28 | Cruces | 9.08652 | −79.6424 | 1 | 1997, 2009 |
| sherman | Fort Sherman | 9.28087 | −79.9747 | 5.96 | 1996, 1997, 1999, 2009 |

Table S2: **Census plots along the Panama Canal used for the regional shares $p_s$.** These 28 plots are those of the Panama network (Condit et al., 2022) that remain after BCI and the six plots on or beside the island are removed, and that have at least one census of every stem of at least 1 cm dbh. Plot names are those of the data release. Coordinates (decimal degrees) and plot areas are from the site table of the network (Condit et al., 2016). Localities are the place names under which the plots were described (Pyke et al. 2001 for the plots named p01–p28, Cocoli and Sherman). Gigante Peninsula and Las Pavas are the sites of the remaining four. Each plot’s stem counts are averaged over the census years listed.

20 cm in height and followed from then on, whatever their later height. The comparison with the model’s seedling pool is in Section S4.

All data analyzed are public. The Dryad and figshare releases are under CC0 licenses, and the Zenodo and Smithsonian releases under CC BY 4.0. The tree censuses are the complete-plot release on Dryad (Condit et al., 2019). Condit (1998) and Hubbell et al. (1999) describe the plot and its census methods. The seedling census is the Dryad release of Comita et al. (2023a, doi:10.5061/dryad.05qfttf85). Columns 2 to 8 of the survival-and-growth matrices $U_s$ are those of the Zenodo archive of Jops and O’Dwyer (2024, doi:10.5281/zenodo.14563651). The seedling column is recomputed from the seedling census and each fecundity row is rescaled to growth rate one, as described below. The regional shares come from the Panama plot-network tables of Condit et al. (2016, Smithsonian, doi:10.5479/data.stri.2016.0622), archived on Dryad (Condit et al., 2022, doi:10.15146/mdpr-pm59). The trait and climate comparisons of Section S5 use further data. Species’ demographic rates in high and low light and their maximum heights are from the Dryad tables of Rüger et al. (2022). Sapling growth and mortality indices, wood density, seed mass and stature are from the figshare supplement (doi:10.6084/m9.figshare.3550359.v1) of Wright et al. (2010), with the maximum height listed by Condit et al. (2020, Dryad) as the stature of species missing there. Each species’ dry-season moisture association is the index (Smithsonian, doi:10.5479/data.bci.20130204) published with Condit et al. (2013). The BCI climate series are the Smithsonian Tropical Research Institute dataset of Paton (2024), and the Oceanic Niño Index is the public-domain series of the NOAA Climate Prediction Center. Our code and the derived tables it produces will be archived with a DOI on publication. Among the tables are our projection matrices, in the format of the Zenodo archive, and the survival and growth rates among classes 2 to 8 estimated for the forecasts from the censuses before each start. The code builds our projection matrices from the published matrices and the seedling census, and reproduces the numbers, tables and figures of the main text and of this supplement from the data listed above.

### Inference of the seedling column

The first column of each survival-and-growth matrix $U_s$, the seedling column, gives the probabilities that a seedling stays in class 1, moves to class 2 or class 3, or dies within five years. A seedling is a stem of class 1, whose dbh is under 10 mm or was not recorded. We estimate this column from the public release of the seedling census (Comita et al., 2023a,b) with the rules below, and take columns 2 to 8 from the archive of Jops and O’Dwyer (2024). In those columns a stem counts as lost when the next census no longer records it alive with a dbh of at least 10 mm. The losses therefore include stems that are alive with no recorded diameter, 24% of the losses when the 90 species and the seven census intervals are pooled. The first of those intervals has a median length of 3.4 years, against 4.8 to 5.3 years for the other six, and is pooled with them. We do not re-estimate these columns for the 90 published species. They are reproduced from the public tree census by the following rule, which we use where we estimate such columns ourselves (Section S6). A tree is counted at a census when its status is A, its dbh is at least 10 mm and its plot coordinates are recorded. Status A means alive, and the tree census gives it to a tree with any living stem or sprout (Condit et al., 2019). For each species the sparsest occupied class among classes 3 to 8, if it holds fewer than 20 such records over all the censuses, is merged into the class below. Each entry is the number of trees in one class at a census and in a given class at the next, summed over the seven census intervals, divided by the number of trees at risk in the starting class, with recorded diameters taken as they stand.

The seedling column is counted on eight pairs of seedling censuses five years apart, 2001–06, 2003–08, 2006–11, 2008–13, 2009–14, 2011–16, 2012–17 and 2013–18. These are the pairs on which the published column of Jops et al. (2025) is reproduced (below). The census has a ninth such pair, 2004–09; with it added the published column is no longer reproduced, and we leave it out so that the two columns are counted on the same pairs. Only the records of the 90 published species are used, and among these, of records with the same identifier and census the first in the file is kept. A seedling is matched between the two censuses of a pair by its identifier and species code, so one recorded under another code at the second census has no record there. A record counts as alive if its status is A, unless N, T or D is its only code; these codes mean nothing found, only the tag found, and dead (Comita et al., 2023b), and a record may carry other codes beside them. Status D means dead or presumed dead. Status Z is given to a seedling that has been tagged in the census of trees and lianas of at least 1 cm dbh and is no longer followed in the seedling census, and the code Ca says that the tag is one of the tree census (Comita et al., 2023b). A seedling alive and in class 1 at the first census of a pair is classed at the second census as follows. It stays in class 1 if it is alive and in class 1. It is promoted to class 2 or class 3 if it is alive with a dbh of at least 10 mm that falls in one of those classes (Table S1). It is promoted to class 2 if the seedling census entered it as handed over to the tree census (status Z, with Ca among its codes) after the first census of the pair and up to the second, and it has no record of status A or D at the second census. Otherwise it counts as dead; this includes a seedling with no record at the second census. For each species the counts are pooled over the eight pairs, a seedling entering once for every pair at whose first census it is alive and in class 1, and each entry of the column is the number with that outcome divided by the number at the start. No minimum count is imposed and no species is pooled with another: the smallest pooled number at the start is 69, and for six of the 84 modeled species fewer than five promotions are counted (Section S6).

These rules follow the census protocol. The tree census takes every stem of at least 1 cm dbh, the lower edge of class 2, so a seedling measured at exactly 10.0 mm has entered class 2. A seedling that grows past 10 mm is tagged at the next tree census, entered once in the seedling census as handed over, and as a rule no longer followed there (Comita et al., 2023b). A hand-over record therefore marks a seedling that survived and grew out of class 1, and we place it in class 2, where the tree census begins. Hand-over records are concentrated at the seedling censuses of 2006, 2011 and 2016, which directly follow a tree census and hold 96% of them, so counting these seedlings as dead would remove most promotions from the pairs that end at those censuses. As a check, a species’ seedling stock (its mean count in the seedling census, scaled to the plot) times its promotion rate, summed over the 84 modeled species, is 1.01 times the 20,503 recruits per five years that the tree censuses record for these species (three intervals of 4.97 to 5.15 years, 2000 to 2015). Here the seedling stock is counted from every record of status A, as in the census counts above; counted with the rule of the seedling column, it gives a ratio of 1.00. The two counts are not independent, since a handed-over seedling is one that the tree census tagged.

Our seedling column differs from that of Jops et al. (2025) only in promotion to class 2. Their published column is reproduced exactly, for all 90 of their species, by the rules above with the handed-over seedlings and the stems of exactly 10.0 mm counted as dead. The entries for staying in class 1 and for promotion to class 3 are therefore the same in both, and each added promotion lowers the five-year probability of death by the same amount. The pairs overlap, so a seedling alive in class 1 at the first census of several pairs is counted once in each; the counts below are of these seedling-intervals. Pooled over the 84 modeled species, 6,649 of the 368,905 seedling-intervals on the eight pairs end in a promotion, 1.80% per five years, against 2,777 (0.75%) in the column of Jops et al. (2025). The added promotions are the 3,719 seedling-intervals that end in a hand-over, from 1,956 seedlings, since a hand-over can fall within more than one overlapping pair, and the 153 that end at exactly 10.0 mm. For two of the modeled species no promotion is added, so their columns are the published ones.

Jops et al. (2025) set the level of each fecundity row so that the species' growth rate is one. We keep that normalization by multiplying each fecundity row by one factor per species (median 0.42 over the 84 modeled species, quartiles 0.34 to 0.48). We use these matrices, with the seedling column above and the rescaled fecundity rows, throughout.

The forecast based on life history inferred from the censuses preceding each start (Section S3) re-estimates survival and growth among classes 2 to 8 from the tree censuses before that start. Applied alone to the censused stems, it involves neither the seedling column nor the fecundity rows. In the forecast of the main text, which is run with the full model, these re-estimated entries replace the corresponding entries of our matrices, and the seedling column and the fecundity rows keep the values estimated from all the censuses; the two give the same predictions one census ahead and nearly the same two censuses ahead (Section S3). Projections of the large trees one census ahead depend only on columns 2 to 8, since in one five-year step no seedling reaches the large-tree classes. From two censuses ahead they also depend on the promotion entries of the seedling column and, through the seedling entry of the starting vector, which is set from the fecundity row (Section S3), on the level of that row. The dependence is slight: for 40 of the 84 species a seedling can reach the large-tree classes in two steps, with probability at most 0.0010.

### Species codes

The plot's species codes for the 84 modeled species are acaldi, alchco, alibed, alsebl, annoac, apeime, aspicr, beilpe, brosal, calolo, cappfr, caseac, cassel, chr1ec, chr2ar, coccma, cordbi, cordla, cou2cu, crotbi, cupasy, des2pa, drypst, ery2ma, eugeco, eugega, eugene, eugeoe, faraoc, gar2in, gar2ma, guargu, guatdu, gustsu, hassfl, heisco, hirttr, hybapr, ingago, ingama, ingaqu, ingas1, ingaum, laciag, laetth, licapl, loncla, maquco, micoaf, micoar, mourmy, ocotce, ocotwh, pentma, picrla, poular, poutre, pri2co, protco, protpa, protte, psycho, pterro, quaras, randar, rinosy, simaam, sipapa, soroaf, stylst, swars1, swars2, tab1ro, tab2ar, tachve, talipr, tet2pa, tri2pa, tri2tu, tripcu, tropra, unonpi, virose, xyllma. The codes of the six species with a published matrix that are not modeled are anaxpa, herrpu, micone, ouralu, paligu, psycma, in the order of their names at the start of this section.

### S3 The decade-ahead forecast in detail

The quantity forecast is a species' change in large trees (stems of at least 79.4 mm dbh, classes 5 to 8) one and two censuses after a starting census (Section S1). It is the species' five-year matrix $A_s$ (that of Jops et al. 2025 with our seedling column and rescaled fecundity row, Section S2) applied once or twice to its observed stage vector at the start, summed over the large-tree classes, minus the starting number. A species-start pair is one species at one start. We treat each census interval as five years. The first lasted 3.4 years and the other six between 4.8 and 5.3 years. An interval of $t$ years could instead be projected with our matrices raised to the fractional power $t/5$. Each stem is measured on a known date within a census, and a more careful treatment would use those dates. Comparisons with censuses after 2015, whose intervals depart further from five years, will need to allow for intervals of different lengths. At this horizon only the survival and growth entries of the censused classes matter. Under any of our matrices a seedling contributes at most 0.0010 stems to the large-tree classes two censuses later, and a class-2 stem at most 0.014 per census.

The forecast leaves out four parts of the full model: competition for space (the capacity, with the lottery that fills vacancies in class 2), immigration, environmental stochasticity and demographic stochasticity. Demographic stochasticity, the chance deaths and transitions of individual stems, scatters the observed change about the prediction, which is the expected change, without moving its mean. Competition and immigration act in the full model only on entry into class 2, where the lottery fills the places that deaths open, and stems admitted to class 2 during an interval do not reach the large-tree classes by the next census. A start census rich in reproductive trees raises the number of candidates for those places but not the number admitted, since candidates already outnumber the places (Section S4). One census ahead, competition and immigration therefore leave the prediction unchanged, and two censuses ahead they could change it slightly. To check, we ran the mean dynamics of the full model, with the seedling stage, fecundity, immigration and the capacity all acting, from the same starting vectors. Its one-census predictions are identical to the forecast. Its two-census predictions differ by 0.41 stems per pair in sample and 0.46 out of sample (1.5% and 1.5% of the mean predicted change), and their rank correlations with the forecast are 0.998 and 0.999.

Environmental stochasticity enters the model as a factor with mean one on sapling growth, shared by all species, and the forecast holds it at its mean. Because the factor moves all species together, it shifts mainly the large-tree total and changes the ranking of the species' changes little. In the forecasts from 1990 to 2015 of Section S7, rescaling the sapling promotions of all species to the rates realized in those years removes 92% of the mean bias and leaves the rank correlation nearly unchanged. The forecast does not reproduce the changes of the total (Table S8), whereas the model with the environmental noise reproduces their size and also widens the spread of outcomes about each species' prediction (Section S7). The forecast thus tests only the survival and growth of the censused stems, and the mechanisms of long-run abundance are tested in Section S4.

Table S3 summarizes the test. In sample, we apply each species' matrix at the first five censuses (420 pairs). Two predictors that use no demographic rates are applied to the same pairs. The long-term trend is the species' mean change per interval over the intervals other than those tested, and the short-term trend is its change over the preceding interval, continued. The rank correlation between the matrices' predicted changes and the observed changes is 0.81 one census ahead and 0.86 two ahead. It exceeds the long-term trend's by +0.16 (90% CI over species +0.10 to +0.24) and +0.21 (+0.14 to +0.32). Between species (mean predicted against mean observed change over the five starts) the rank correlation is 0.96 and 0.98. Within species (each start's deviation from the species' mean) it is 0.28 and 0.29, and it is positive for 62 of the 79 species whose numbers vary. A long-term trend does not distinguish one start of a species from another, and the forecast does so through the stage structure at the start. The mean of the 84 matrices, applied to each species' own stage vector, gives only 0.02 and 0.01, so the accuracy of the forecast depends on each species having its own matrix.

Most forest plots have only a few censuses. To test the forecast under that constraint we estimate the rates from earlier censuses only, and we call the result the out-of-sample forecast. At each start, each species' survival and growth frequencies among classes 2 to 8 are estimated from all the censuses before it (1982–1995 for the start of 1995, 1982–2000 for 2000 and 1982–2005 for 2005), applied to the stage vector at that start, and compared with the censuses through 2015 (252 pairs at each horizon; Table S4). No interval at or after a start enters its rates. The frequencies are counted as in the published columns 2 to 8 (Section S2): a stem leaves the classes when the next census does not record it alive with a dbh of at least 10 mm, and the counts are pooled over the intervals, the first interval entering as one step. Unlike the rule of that section, these counts keep the trees with no recorded plot coordinates. Each species keeps the range of classes of its published matrix, and a stem outside that range is counted in the nearest class within it. A class with fewer than five stems at risk takes the frequencies pooled over all species; over the three starts this applies to one stem. Estimated from all seven intervals, these counts agree with the published columns to within 0.002 in every entry of the 447 columns that have at least 20 stems at risk. The figures that the main text quotes for the same pairs with rates from all the censuses use the published survival and growth entries among classes 2 to 8 in the same way, applied to the censused classes, with no seedling class and no fecundity row. The out-of-sample forecast is biased upward. Over 1995–2005

Table S3: **Species’ matrices rank the changes in large trees better than either trend, in and out of sample.** Change in each species’ large trees (stems of at least 79.4 mm dbh) one and two censuses ahead, predicted from its matrix and from two predictors that use no demographic rates. A species-start pair is one species at one starting census. Sign is the percentage of pairs with a nonzero observed change whose direction is predicted correctly. $\rho$ is the Spearman rank correlation over pairs (90% CI over species). Within is the rank correlation of each start’s deviation from its species’ mean, undefined for a long-term trend that is the same at every start. MAE is the mean absolute error in stems. The long-term trend is each species’ mean change per interval outside the predicted intervals (in sample) or from 1982 to the start (out of sample), and the short-term trend is its change over the previous interval. Both are doubled at two censuses. Reversals are observed changes one census ahead opposite in sign to the species’ net change from 1982 to the start, counted in sample at every start from 1985 to 2010 (504 pairs). Predicting no change gives MAE 16.1 and 30.4 stems.

| predictor | one census ahead |  |  |  |  | two censuses ahead |  |  |  | reversals anticipated |
| --- | --- | --- | --- | --- | --- | --- | --- | --- | --- | --- |
| | pairs | sign (%) | $\rho$ (90% CI) | within | MAE | sign (%) | $\rho$ (90% CI) | within | MAE | |
| *In sample: rates from 1982–2015; starts 1982–2000* |  |  |  |  |  |  |  |  |  |  |
| projection matrices | 420 | 84 | 0.81<br>(0.75–0.87) | 0.28 | 8.0 | 86 | 0.86<br>(0.80–0.91) | 0.29 | 12.1 | 78 of 128 (61%, 90% CI 52–69%); trend up to the start, none |
| long-term trend, other intervals | 420 | 74 | 0.66 | – | 10.0 | 71 | 0.65 | – | 18.7 | – |
| short-term trend | 336 | 72 | 0.67 | – | 11.2 | 72 | 0.64 | – | 21.0 | – |
| *Out of sample: at each start, rates from all censuses before it; starts 1995, 2000 and 2005* |  |  |  |  |  |  |  |  |  |  |
| projection matrices | 252 | 75 | 0.72<br>(0.61–0.81) | 0.09 | 9.7 | 72 | 0.70<br>(0.58–0.80) | 0.10 | 18.5 | – |
| long-term trend, 1982 to the start | 252 | 69 | 0.58 | –0.24 | 10.2 | 66 | 0.55 | –0.20 | 19.9 | – |
| short-term trend | 252 | 74 | 0.64 | –0.12 | 8.3 | 71 | 0.62 | –0.23 | 16.4 | – |
| rates frozen at 1982–1995 | 252 | 74 | 0.71<br>(0.60–0.80) | 0.47 | 11.4 | 71 | 0.68<br>(0.55–0.79) | 0.46 | 22.1 | – |

it lies above the observed change for 65 of the 84 species, and the predicted changes sum to +1,167 stems against -578 observed. One census ahead, the forecasts sum to +1,172 large trees against -517 observed.

Both trends rank the later changes less well than the out-of-sample forecast at either horizon (Tables S3 and S4). The forecast’s rank correlation exceeds that of each species’ long-term trend from 1982 to the start, continued, by +0.14 (90% CI +0.07 to +0.23) one census ahead and +0.15 (+0.07 to +0.25) at two. It exceeds that of the short-term trend by +0.08 (+0.01 to +0.14) and +0.08 (+0.01 to +0.15). The forecast’s skill lies in the ordering of species. Between species its rank correlation is 0.79 one census ahead and 0.76 at two, and within species it is 0.09 (90% CI -0.05 to 0.25) and 0.10 (-0.06 to 0.26). The long-term trend’s within-species correlation is -0.24 and -0.20. That of the short-term trend is negative (-0.12), because larger changes tend to be followed by smaller ones, and its overall rank correlation of 0.64 comes from the ordering of species (between species 0.88).

Updating the rates at each start, in place of rates frozen at 1982–1995 for every start, changes the rank correlations little, by +0.01 (90% CI -0.02 to +0.04) one census ahead and +0.02 (-0.02 to +0.06) at two. With the frozen rates they are 0.71 and 0.68 (Table S3, Fig. S1b). The frozen rates order the species less well (between species 0.74 and 0.71) and keep more of the within-species timing (0.47, 90% CI 0.33 to 0.60, and 0.46). The species’ long-term trends over 1982–1995 rank the changes at 0.54 and 0.51. Shrinking each species’ frequencies toward the community’s by the method of moments changes the rank correlation of the frozen rates by +0.01 (90% CI -0.01 to +0.02) at one census, because every species followed has at least 774 stem-intervals in those censuses. Dropping the first interval changes it by -0.03 and -0.02. Table S4 also gives single tested intervals. For every set of rates that ends before 2010, $\rho$ is lowest for the change from 2010 to 2015, which for most species continued the change over 2005 to 2010 (rank correlation 0.756).

Out of sample, too, the forecast depends on each species having its own matrix (Table S5). The arithmetic mean of the 84 matrices estimated before each start, applied to each species’ own stage vector, ranks the changes at 0.24 one census ahead and 0.26 at two (sign right in 54% and 55% of the pairs with a nonzero observed change). The species’ own matrices exceed these values by +0.48 (90% CI +0.28 to +0.68) and +0.44 (+0.22 to +0.65). One matrix from the pooled stems of all species gives 0.24 and 0.26. We also permuted the matrices among species at random 1,000 times, with one permutation per draw used at every start. A permutation may leave a species with its own matrix, which happens to about one species per draw, so almost every species is projected by another species’ matrix. The median rank correlations are 0.16 and 0.17 (5th to 95th percentiles 0.00 to 0.31 and 0.02 to 0.34), and the largest, 0.44 and 0.47, are below the 0.72 and 0.70 of the species’ own matrices. With the rates frozen at 1982–1995, and in sample with our matrices, the mean matrix and the permutations also fall well short of the species’ own matrices (Table S5).

Table S6 repeats the test for every contiguous division of the censuses into those that supply the rates and those tested, forward and backward in time. Each division is compared with the long-term trend over the censuses that supply its rates. In the 18 cases with at least two intervals behind the rates, the rank correlation lies between 0.60 and 0.73 and exceeds the long-term trend’s by +0.05 to +0.22. The lowest values come from rates estimated on a single interval. Leaving out only the intervals a pair is tested on gives 0.75 and 0.77 over the 420 pairs. The between-species order is then intact (0.95 and 0.97) and the within-species correlation is negative (-0.05 and -0.20), as expected when the transitions removed are those of the tested intervals.

A reversal is an observed change one census ahead opposite in sign to the species’ net change from 1982 to the start, and a continuation is one of the same sign. Pairs with a zero on either side are in neither class. A long-term trend, continued, cannot change sign, whereas a projection of the stage structure at the start under fixed rates can. In sample, at every start from 1985 to

| rates estimated from | tested on | pairs | sign (%) | $\rho$ trend | $\rho$ within | MAE |
| --- | --- | --- | --- | --- | --- | --- |
| 1982 to each start | 1995, 2000, 2005 → one ahead | 252 | 75 | 0.72 | 0.58 | 0.09 9.7 |
| 1982 to each start | the same → two on | 252 | 72 | 0.70 | 0.55 | 0.10 18.5 |
| 1982–1995 | 1995, 2000, 2005 → one ahead | 252 | 74 | 0.71 | 0.54 | 0.47 11.4 |
| 1982–1995 | the same → two on | 252 | 71 | 0.68 | 0.51 | 0.46 22.1 |
| 1982–1995 | 1995 → 2005 | 84 | 77 | 0.75 | 0.59 | 22.1 |
| 1982–1995 | 2005 → 2010 | 84 | 69 | 0.63 | 0.45 | 11.4 |
| 1982–2005 | 2005 → 2010 | 84 | 69 | 0.62 | 0.50 | 8.6 |
| 1982–2005 | 2005 → 2015 | 84 | 66 | 0.61 | 0.51 | 16.9 |
| 1982–2005 | 2010 → 2015 | 84 | 70 | 0.56 | 0.51 | 10.1 |
| 1982–2010 | 2010 → 2015 | 84 | 74 | 0.63 | 0.59 | 9.3 |
| 1982–2015 (in sample) | 2005 → 2010 | 84 | 80 | 0.80 | 0.68 | 6.4 |
| 1982–2015 (in sample) | 2010 → 2015 | 84 | 82 | 0.78 | 0.71 | 8.0 |
| published fits (in sample) | 2005 → 2010 | 84 | 80 | 0.80 | 0.68 | 6.4 |
| short-term trend | 1995, 2000, 2005 → one ahead | 252 | 74 | 0.64 | –0.12 | 8.3 |
| pooled matrix, 1982–1995 | the same | 252 | 53 | 0.26 | 0.40 | 21.3 |

Table S4: **Matrices estimated from part of the census series rank the later changes better than the long-term trend over the same censuses.** Each species’ survival and growth frequencies among classes 2 to 8 are estimated from the censuses in the first column and applied to its stage vector at the start, and the predicted change in its large trees is compared with the census in the second column. Published fits: the survival and growth entries among classes 2 to 8 of the matrices of Jops et al. (2025), estimated from all seven intervals and used unchanged. Sign, $\rho$, within and MAE are as in Table S3. Trend $\rho$ is the rank correlation for the species’ long-term trend, its mean change per interval over the censuses in the first column, continued. The pooled matrix is one matrix from the pooled stems of all species. In the rows above the first rule no tested interval enters the rates. The first two rows and the short-term trend row repeat the out-of-sample rows of Table S3, and the next two hold the rates at 1982–1995 for every start.

Table S5: **A mean or permuted matrix ranks the changes far less well than each species’ own matrix.** Rank correlation $\rho$ between predicted and observed change in large trees one and two censuses ahead (columns “one” and “two”), when each species’ stage vector is projected with its own matrix, with the arithmetic mean of the 84 matrices, and with the matrices permuted among species at random (median and largest $\rho$ over the permutations). The first two rows are out of sample and the third is in sample, on the pairs of Table S3.

| rates estimated from | own matrix |  | mean matrix |  | permuted, median |  | permuted, largest |  |
| --- | --- | --- | --- | --- | --- | --- | --- | --- |
|  | one | two | one | two | one | two | one | two |
| 1982 to each start | 0.72 | 0.70 | 0.24 | 0.26 | 0.16 | 0.17 | 0.44 | 0.47 |
| 1982–1995 | 0.71 | 0.68 | 0.26 | 0.27 | 0.17 | 0.18 | 0.46 | 0.49 |
| 1982–2015 (in sample) | 0.81 | 0.86 | 0.02 | 0.01 | 0.03 | 0.05 | 0.33 | 0.30 |

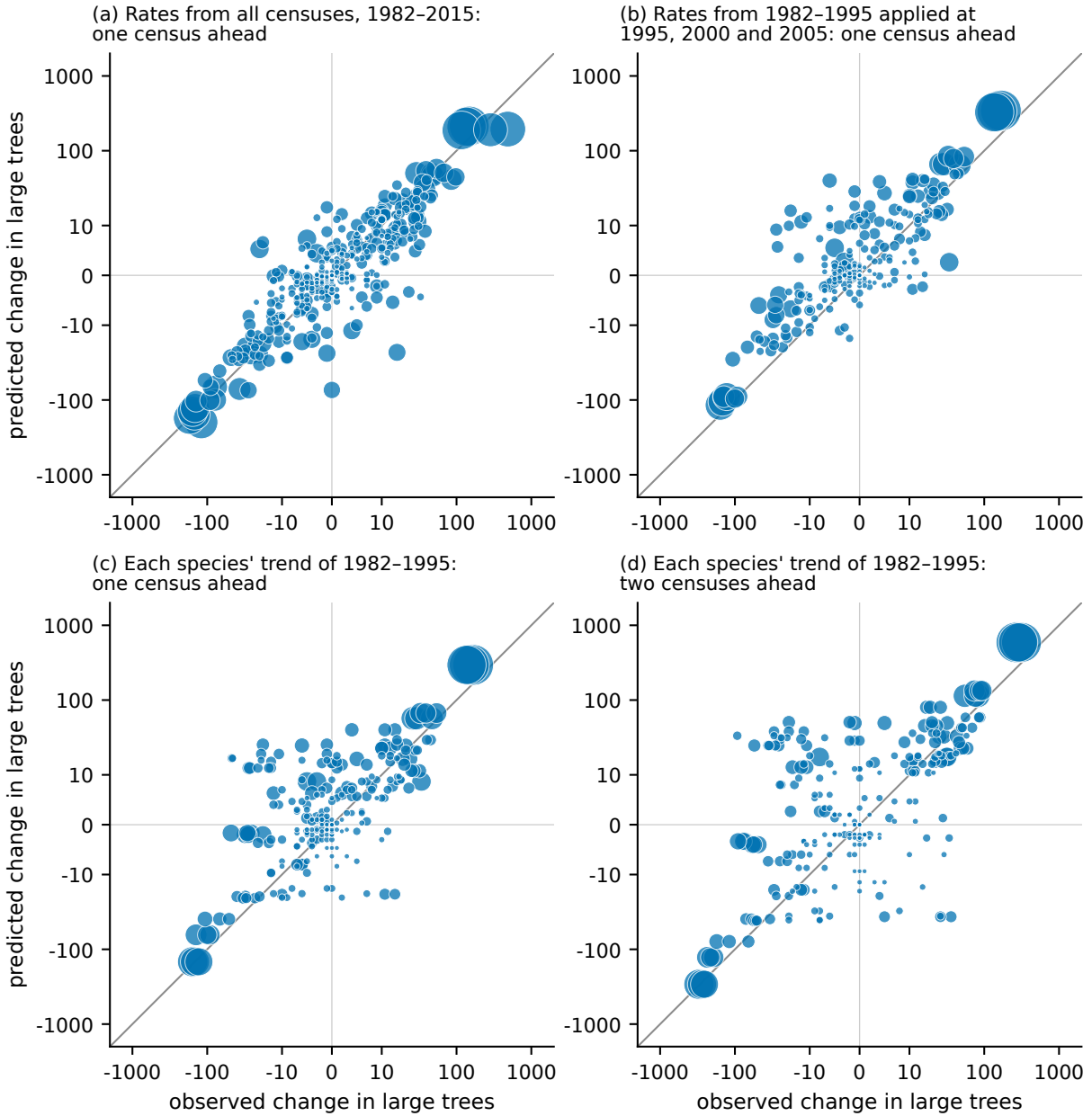

Figure S1: **Each species' matrix predicts the change in its large trees one census ahead, in and out of sample.** Predicted against observed change in each species' large trees one census ahead at Barro Colorado. Each point is a species-start pair, symbol area scales with the species' large trees at the start, the line is the identity, and the axes are logarithmic away from zero. (a) In sample: our matrices applied at the first five censuses (420 pairs); rank correlation 0.81. (b) Out of sample: rates from the 1982–1995 censuses alone, applied at 1995, 2000 and 2005 (252 pairs); 0.71. (c) The long-term trend on the pairs of (b), each species given its mean change per interval over 1982–1995; 0.54. (d) The same long-term trend two censuses ahead; 0.51. Fig. 3 of the main text shows the changes two censuses ahead, with the rates from all censuses before each start in its panel (a) and from all censuses in its panel (b).

Table S6: **Matrices rank the decade-ahead changes better than the long-term trend however the censuses are divided.** The decadal test under every contiguous division of the censuses. The first columns give the censuses whose intervals supply the transition frequencies and the number of intervals. For each horizon the table gives the starting censuses tested, the number of pairs (84 per start), the rank correlation $\rho$ over them, the same for the long-term trend over the same censuses (each species' mean change per interval, times the horizon), and the difference. Forward rows use the first one to six intervals and backward rows the last one to six. The first row is the in-sample row of Table S3. The last row leaves out only the interval a pair is tested on (two intervals, two censuses ahead), from the rates and from the trend alike. <sup>†</sup>Rates from one interval. A dash marks an entry with no census two steps beyond the last start.

|  | rates estimated from |  | one census ahead |  |  |  |  | two censuses ahead |  |  |  |  |
| --- | --- | --- | --- | --- | --- | --- | --- | --- | --- | --- | --- | --- |
| | censuses | int. | starts | pairs | $\rho$ | trend | diff. | starts | pairs | $\rho$ | trend | diff. |
| all intervals | 1982–2015 | seven | 1982–2000 | 420 | 0.81 | – | – | 1982–2000 | 420 | 0.86 | – | – |
| forward<sup>†</sup> | 1982–1985 | 1 | 1985–2010 | 504 | 0.55 | 0.45 | +0.10 | 1985–2005 | 420 | 0.60 | 0.49 | +0.11 |
|  | 1982–1990 | 2 | 1990–2010 | 420 | 0.66 | 0.54 | +0.12 | 1990–2005 | 336 | 0.68 | 0.54 | +0.14 |
|  | 1982–1995 | 3 | 1995–2010 | 336 | 0.66 | 0.51 | +0.15 | 1995–2005 | 252 | 0.68 | 0.51 | +0.18 |
|  | 1982–2000 | 4 | 2000–2010 | 252 | 0.64 | 0.52 | +0.12 | 2000, 2005 | 168 | 0.66 | 0.52 | +0.14 |
|  | 1982–2005 | 5 | 2005, 2010 | 168 | 0.60 | 0.51 | +0.09 | 2005 | 84 | 0.61 | 0.51 | +0.10 |
|  | 1982–2010 | 6 | 2010 | 84 | 0.63 | 0.59 | +0.05 | – | – | – | – | – |
| backward<sup>†</sup> | 2010–2015 | 1 | 1982–2005 | 504 | 0.61 | 0.51 | +0.10 | 1982–2000 | 420 | 0.65 | 0.53 | +0.12 |
|  | 2005–2015 | 2 | 1982–2000 | 420 | 0.61 | 0.46 | +0.16 | 1982–1995 | 336 | 0.65 | 0.49 | +0.16 |
|  | 2000–2015 | 3 | 1982–1995 | 336 | 0.66 | 0.49 | +0.16 | 1982–1990 | 252 | 0.68 | 0.53 | +0.15 |
|  | 1995–2015 | 4 | 1982–1990 | 252 | 0.68 | 0.47 | +0.22 | 1982, 1985 | 168 | 0.68 | 0.50 | +0.19 |
|  | 1990–2015 | 5 | 1982, 1985 | 168 | 0.71 | 0.53 | +0.18 | 1982 | 84 | 0.73 | 0.59 | +0.14 |
|  | 1985–2015 | 6 | 1982 | 84 | 0.67 | 0.53 | +0.14 | – | – | – | – | – |
| all but the tested window | 1982–2015 | six (five) | 1982–2000 | 420 | 0.75 | 0.66 | +0.10 | 1982–2000 | 420 | 0.77 | 0.65 | +0.12 |

| predictor | reversals anticipated |  | continuations anticipated |  |
| --- | --- | --- | --- | --- |
|  | one census ahead (of 128) | two censuses ahead (of 110) | one census ahead (of 306) | two censuses ahead (of 261) |
| published fits (in sample) | 78 (61%) | 70 (64%) | 288 (94%) | 245 (94%) |
| rates without the tested interval (out of sample) | 66 (52%) | 53 (48%) | 278 (91%) | 235 (90%) |
| rates from 1982–1995 | 49 (38%) | 42 (38%) |  |  |
| long-term trend up to the start | 0 | 0 | 306 | 261 |
| short-term trend | 41 (32%) | 31 (28%) | 270 | 234 |

| starting census (one census ahead) | 1985 | 1990 | 1995 | 2000 | 2005 | 2010 |
| --- | --- | --- | --- | --- | --- | --- |
| reversals | 26 | 16 | 17 | 23 | 26 | 20 |
| anticipated by the published fits (in sample) | 17 | 12 | 10 | 16 | 13 | 10 |
| by the rates without the tested interval | 16 | 11 | 9 | 12 | 11 | 7 |
| by the rates from 1982–1995 | 16 | 6 | 5 | 7 | 9 | 6 |

Table S7: **Life history matrices anticipate more reversals in a species’ direction of change than either trend.** All 84 species at every start with an earlier interval, 1985 to 2010 one census ahead (504 pairs) and 1985 to 2005 two ahead (420 pairs). A reversal is a pair whose observed change in large trees has the sign opposite to the species’ net change from 1982 to the start, and a continuation is one of the same sign. Pairs with no change on one side are in neither column. Published fits: the survival and growth entries among classes 2 to 8 of our matrices, estimated from all seven intervals. Rates without the tested interval are the same entries re-estimated from the other intervals, and are out of sample at every start. Rates from 1982 to 1995 are the early matrices at every start. Their continuation entries are omitted because two of the six starts lie inside the years they were estimated from. The long-term trend from 1982 to the start cannot change sign, and the short-term trend repeats the sign of the last interval. The lower table gives the reversals from each start one census ahead and the number each set of rates anticipates.

2010 (504 pairs), our matrices anticipate 78 of the 128 reversals one census ahead, 61% (90% CI over species 52 to 69%), and 70 of 110 two ahead; they give the correct direction for 288 of the 306 continuations (Table S7). Re-estimated without the tested interval, and so out of sample at every start, the same rates anticipate 66 reversals, 52% (43 to 60%). Including the tested interval in the rates thus accounts for 9 points of the reversal fraction, and the rest is due to the stage structure and the rates at the start. The short-term trend repeats the sign of the previous interval and anticipates only reversals already begun. On the pairs from 1995, 2000 and 2005, one census ahead, the out-of-sample forecast anticipates 23 of the 66 reversals (35%), the rates frozen at 1982–1995 anticipate 21, and the published fits, which include the tested intervals, anticipate 39. On the same pairs the short-term trend anticipates 29 (44%) and the long-term trend since 1982 none.

Rescaling the early matrices, those estimated from 1982–1995, by each species’ observed net growth rate does not improve the forecast, because the matrices already contain it. The ratio $\gamma_s$ of a species’ observed five-year growth factor in large trees over 1982–1995 to the factor its matrix implies has median 1.00 (range 0.96 to 1.10). Multiplying each early matrix by $\gamma_s^h$ at $h$ censuses changes the out-of-sample rank correlation by $-0.005$ (90% CI $-0.014$ to $+0.003$). Projected four intervals forward, to 2015, the early matrices predict 24,556 large trees and the long-term trend continued for four intervals predicts 23,967, against 21,859 censused. The net rates of 1982–1995 did not persist to 2015, which is why the stationary model holds every species at growth rate one and lets immigration set absolute abundances.

The census does not enumerate seedlings, so we complete the seedling entry of each stage vector from the species’ fecundity row and its censused counts. Deleting the seedling class instead (Section S6) changes the predicted large-tree total by 19.2 stems on average over the five starts, at most 0.12% of the observed total, and leaves the two-census rank correlation unchanged (0.86 without the class, 0.86 with it). Summed over species, the projections do not reproduce the change in the large-tree total (Table S8). For four of the five starts the observed and projected changes have opposite signs. Those changes are shared across species, and a projection with the environment at its mean cannot produce them. With the environmental noise at its estimated amplitude, the modeled total moves from census to census by amounts of the observed size (Section S7).

| starting census | large trees | observed $\Delta_5 N$ | predicted $\Delta_5 N$ | observed $\Delta_{10} N$ | predicted $\Delta_{10} N$ |
| --- | --- | --- | --- | --- | --- |
| 1982 | 20,952 | +288 | -166 | +1,152 | -211 |
| 1985 | 21,240 | +864 | -28 | +1,004 | +2 |
| 1990 | 22,104 | +140 | +107 | -183 | +255 |
| 1995 | 22,244 | -323 | +128 | -578 | +287 |
| 2000 | 21,921 | -255 | +126 | -194 | +306 |

Table S8: **With the environment held at its mean, the matrices do not predict the change in the large-tree total.** Predicted and observed change in the total of large trees of the 84 modeled species, from each of the first five censuses, one ($\Delta_5 N$) and two ($\Delta_{10} N$) censuses ahead. The prediction is the decadal change (Section S1) summed over species. For each species separately the same matrices largely reproduce the changes (Table S3).

We test the forecast against the null hypothesis of neutral drift among equivalent species, using Hubbell’s model with the parameters fitted to the plot. The community is of fixed size, every tree has the same per capita rates of death and recruitment, and each vacancy goes with probability $m = 0.026$ to an immigrant from a fixed pool with the plot’s starting composition. Under this null nothing measured on a species at a census, its size structure included, can rank its later changes. Let $K_i$ be the large trees that died and were replaced in interval $i$ and $p_i$ a species’ share of them. Under drift the species’ number changes with variance $2K_i p_i(1 - p_i)$ per interval, so we express its observed change $\Delta N_s$ from the start to 2015 in units of the drift standard deviation $\sigma_s$,

$$\sigma_s = \left[ \sum_i 2K_i p_i(1 - p_i) \right]^{1/2}, \quad z_s = \Delta N_s / \sigma_s. \quad (\text{S1})$$

The predictor is demographic momentum (Caswell, 2001), the factor by which a species’ large trees would eventually change under its 1982–1990 rates from its 1985 size structure, with recruitment replacing deaths. It is defined for the 75 species with a matrix estimated from those years and a large tree at the start. The outcome is the change from 1990 to 2015, so no interval enters both predictor and outcome. We compare the Spearman correlation $S$ between demographic momentum and $z_s$ with three null distributions of 200,000 draws each (Table S9): drift at the measured rate of replacement, drift at the fitted rate, which is 4.51 times faster, and the demographic momenta reassigned among species at random. The observed value is $S = +0.67$. None of the 600,000 draws reaches it (largest +0.52), so $P < 10^{-5}$ under each null, and drift is rejected with more than 99.9% confidence. The model’s forecast from 1990, in the same units, gives $S = +0.73$, also beyond every draw. A neutral model whose species share one set of size-dependent rates, the pooled matrix of the 84 species applied to each species’ 1990 structure, is rejected as well. Its expected changes rank the observed ones at 0.22, and over 20,000 simulated forests $S$ never reaches +0.15 ($P < 5 \times 10^{-5}$).

Table S9: **Demographic momentum and the model’s forecast rank species’ later changes beyond every draw of three neutral nulls.** Each predictor is correlated across the 75 species with the change in large trees from 1990 to 2015, in units of each species’ neutral drift (Eq. (S1)). $S$ is the Spearman correlation. A, B and P give how many of 200,000 draws reached $S$ under drift at the measured rate of replacement, under drift at the fitted rate, 4.51 times faster, and under random reassignment of the predictor. The remaining columns give the reassignment null’s standard deviation, the largest 99.9th percentile and the largest draw over the three nulls, and $z_S = S\sqrt{n-1}$. The first row is the test described in the text.

| start | predictor | $n$ | $S$ | A | B | P | null s.d. | 99.9th | largest | $z_S$ |
| --- | --- | --- | --- | --- | --- | --- | --- | --- | --- | --- |
| 1990 | demographic momentum, 1985 structure | 75 | +0.667 | 0 | 0 | 0 | 0.116 | +0.36 | +0.52 | 5.7 |
| 1990 | model forecast, 1990 structure | 75 | +0.728 | 0 | 0 | 0 | 0.116 | +0.36 | +0.52 | 6.3 |

### S4 The stationary state and long-run composition

#### The closed form

Each species $s$ is a stage-structured population, a vector $n_s(t)$ of stem counts in a seedling class (class 1) below the census threshold and in the diameter classes above it, projected one census interval $\Delta t$ ahead by the species’ measured matrix,

$$n_s(t + \Delta t) = A_s n_s(t), \quad A_s = U_s + F_s, \quad (\text{S2})$$

where $U_s$ and $F_s$ are the survival-and-growth and fecundity matrices of Jops et al. (2025), except that the seedling column and the level of the fecundity row are ours (Section S2). The matrix $U_s$ holds survival and movement among classes. The matrix $F_s$ holds seed production and is nonzero only in its first row, whose level is fixed by the normalization of each species’ growth rate to one. The model adds to this projection a stream of immigrant seed and the lottery, under the capacity, for the places that deaths open. With the environmental noise off and every stem-by-stem draw replaced by its expectation, the stationary stage vector $\bar{n}_s$ of species $s$ is the mean-field fixed point of those dynamics, $\bar{n}_s = (U_{d,s} + F_s) \bar{n}_s + M w_s e_1$. Here $e_1$ is the unit vector on the seedling class and $M w_s$ is the expected number of immigrant seeds per five-year interval. The matrix $U_{d,s}$ is $U_s$ with its class-1 promotion entries multiplied by $1 - d$ and the removed probability returned to the class-1 diagonal, where $d$ is the waiting fraction of the main text, the fraction of candidate recruits not admitted to the censused stand in a step, the same for every species. The capacity fixes $d$, because the sum of $\bar{n}_s$ over species and over the censused classes 2 to 8 must equal the observed number of censused stems, $K$. Its value follows in closed form (Eq. (S4) below). At Barro Colorado $d$ is 0.158 under the regional pool and 0.025 under the trade-off scheme.

Because every matrix $U_s + F_s$ has a long-run growth rate of one, the inverse that gives $\bar{n}_s$ factorizes. Let $\phi_s$ be the stable stage vector of $U_s + F_s$, $\mu_s$ a seedling’s probability of dying in a step, and $\Omega_s = \sum_{k=5}^8 \phi_{s,k} / \phi_{s,1}$ the number of large trees (stems of at least 79.4 mm dbh, classes 5 to 8) per seedling in that stable structure. Then

$$\bar{n}_s = M w_s \left[ \frac{e_1}{\mu_s} + \frac{1-d}{d \mu_s} \frac{\phi_s}{\phi_{s,1}} \right], \quad (\text{S3})$$

as substitution into the fixed-point equation confirms, using $(U_s + F_s) \phi_s = \phi_s$ and the facts that $U_{d,s}$ differs from $U_s$ only in its first column and that $F_s$ has no entry in that column. Above the seedling class every species therefore has its stable stage structure $\phi_s$, whatever the abundances of the other species. The immigration weight, the arrival rate, the lottery (through $d$) and any rescaling of fecundity set only the constant of proportionality.

Summed over classes 5 to 8, Eq. (S3) gives $\bar{c}_s = M w_s G_s$ large trees, with $G_s = (1 - d) \Omega_s / (d \mu_s)$ the local gain of the main text. The large-tree composition, $w_s \Omega_s / \mu_s$ normalized over species, contains neither $d$ nor $M$. Under the regional pool ($w_s = p_s$, the regional relative abundance) it is $p_s \Omega_s / \mu_s$ normalized, and under the trade-off scheme ($w_s \propto p_s / G_s$) it is the regional composition $p_s$ itself. The arrival rate cancels from every censused abundance as well, because the capacity condition constrains only the second term of Eq. (S3) and so fixes the product $M(1 - d)/d$. In class 1 the two terms sum to $1/(d \mu_s)$ standing seedlings per arriving seed. The immigrant seeds themselves are the fraction $d$ of these, and the seed of the standing trees is the rest. With $M(1 - d)/d$ fixed, the seedling pool responds to $M$ only through the immigrant part, so that halving $M$ lowers the predicted seedling density by the fraction $d/2$ and leaves the censused abundances unchanged, and no value of $M$ lowers it by more than the fraction $d$. Section S6 explains how $M$ is chosen.

The waiting fraction itself has a closed form. Summed over the censused classes, Eq. (S3) gives $M w_s (1 - d) C_s / (d \mu_s)$ censused stems of species $s$, with $C_s = \sum_{k=2}^8 \phi_{s,k} / \phi_{s,1}$ the censused stems per seedling in its stable structure. The capacity condition, $\sum_s M w_s (1 - d) C_s / (d \mu_s) = K$, is linear in $(1 - d)/d$, so that

$$d = \frac{MS}{K + MS}, \quad S = \sum_s w_s \frac{C_s}{\mu_s}, \quad (\text{S4})$$

where $S$ is an average over the arriving seed of the censused stems per seedling divided by the seedling mortality, weighted by the immigration shares. The waiting fraction is thus a function of the matrices, the shares $w_s$, the arrival rate and the capacity, and the numerical solution of the capacity condition in our code agrees with Eq. (S4) to within a relative difference of $10^{-7}$. Under the trade-off scheme the gain $G_s = (1 - d) \Omega_s / (d \mu_s)$ carries the same factor $(1 - d)/d$ for every species, which the normalization of $w_s$ removes, so $w_s \propto p_s \mu_s / \Omega_s$ and Eq. (S4) is explicit under both schemes. The waiting fraction rises with the arrival rate and with the number of censused stems per seedling in the arriving species, and falls with their seedling mortality and with the capacity, since $(1 - d)/d = K/(MS)$. The product $MS$ is 37,726 stems under the regional pool and 5,227 under the trade-off scheme. If all species had the same life history, then $S = C/\mu$ and $d = 1/(1 + K\mu/(MC))$. Taken one at a time, the 84 life histories would give $d$ from 0.0011 to 0.66 (median 0.061) at the arrival rate and capacity used throughout. The trade-off scheme weights arrivals toward species of small $\Omega_s/\mu_s$, which have small $C_s/\mu_s$ as well, so its $S$ and its $d$ are the smaller.

At Barro Colorado the inputs are the 84 matrices $U_s + F_s$, each normalized to growth rate one, the regional shares $p_s$ (Section S2), the arrival rate $M$ of 26,499 seeds per interval (Section S6) and the capacity $K$ of 200,721 censused stems, the census mean. The trade-off scheme is the stationary reference of the main text, and the regional pool is shown beside it for comparison.

The nine modeled species absent from the regional plots have no measured regional share. We give them the smallest share that any sampled species receives (Section S2), in all closed-form values of the main text and of this supplement. The simulations used below to test the closed form and to measure the settling time gave these species zero weight instead, and the closed form compared with those simulations uses the same weights. To see how much this choice matters, we also solved Eq. (S3) with zero weight for these species. The rank correlation with the census is then unchanged (0.672 under the regional pool, 0.727 under the trade-off scheme). The number of species predicted below one large tree rises from 17 to 18 (regional pool) and from 5 to 11 (trade-off scheme), and the identity-line $R^2$ on log abundance falls from $-0.715$ to $-0.756$ and from $+0.494$ to $+0.396$, because a species with no arrivals is predicted at zero where the census has a few stems.

### Simulations and demographic noise

The simulations follow every stem through the dynamics of Eq. (S2) in five-year steps with full demographic noise. Each stem’s fate is a multinomial draw from its species’ matrix, seed production and immigration are Poisson draws, and the places that deaths open among censused stems are filled by a lottery, blind to species, among the seedlings drawn for promotion. With the environmental noise on, every species’ sapling promotions are multiplied by that step’s environmental-noise factor $g_t$ (Eq. (S5), Section S5) before the draws. The mean-field fixed point above is an approximation to the expectation of this stochastic model. Every rate except the recruitment lottery is linear in the numbers of stems, and for linear rates demographic noise leaves the expected abundances exactly on the deterministic solution, as in birth, death and immigration treatments of neutral models (McKane et al., 2000; Volkov et al., 2003). The lottery is nonlinear in the numbers of candidates and vacancies, so the mean-field fixed point is not the exact expectation of the stochastic model. Solving for the exact expectation remains an open question, which we do not pursue because the simulations with full demographic noise depart from the mean field by little.

With the environmental noise off, Eq. (S3) reproduces the large-tree total of every simulated stationary state to within 0.3% and its species abundances at a rank correlation of 0.998 or better. Under the regional pool the median species’ simulated mean differs from the closed form by 0.8%. This difference is the only way in which demographic variance (Jops et al., 2025) could affect abundance other than through the mean field, and it shows no trend with a species’ demographic variance. Regressed on log demographic variance, the log ratio of simulated to closed-form abundance changes by at most 0.04 ($\log_{10}$ units) across the species under the regional pool. Species with low demographic variance therefore appear to be favored only through their larger local gain. Across the 79 species whose matrices reach the large-tree classes, the gain falls with demographic variance at a log-log slope of -0.88 (rank correlation -0.72).

The stationary abundances of main text Fig. 4 are those of the simulations with the environmental noise on, run from two unrelated sets of initial conditions until the stationarity test described below is met and averaged over the last five windows of both sets (Section S7). The closed form of Eq. (S3) is their mean-field reference, and the values in this supplement are those of the closed form unless we say otherwise (Table S10 gives both). In the simulations the rank correlation with the census is within 0.013 of the closed form’s under both schemes, and under the trade-off scheme the simulated abundances have a rank correlation of 0.994 with the closed form. Under the regional pool the $R^2$, the large-tree total and the seedling pool also differ little from the closed form’s. Under the trade-off scheme the environmental noise lowers the large-tree total from 1.23 to 0.84 times the observed and the $R^2$ from +0.494 to +0.362 (Section S7).

### Long-run composition

The capacity fixes only the total number of censused stems and leaves their distribution among size classes free. Under the trade-off scheme 13.31% of the censused stems stand in the large-tree classes, against 10.81% in the census and 26.09% under the regional pool (Table S11, upper table; Fig. S2a). Within species the share is nearly right, because each species keeps its stable stage structure at the stationary state (Eq. (S3)) and that structure places a share of the species’ censused stems in the large-tree classes close to the observed share (Table S11, last rows). At the observed species composition these structures would put 12.90% of the stand in those classes. The excess under the regional pool therefore comes from species composition, which that scheme weights toward species whose stable structures hold many large trees. The trade-off weighting shifts the composition away from them. As a species abundance distribution (Fig. S2b), the trade-off scheme’s stationary stand lies at Kolmogorov distance 0.127 from the census over the 79 species that reach the large-tree classes, closer than the regional pool (0.287), a log-series (0.37) and Hubbell’s dispersal-limited model fitted to the same distribution (0.18). In the simulations with environmental noise (main text Fig. 4c) the distances are 0.215 under the trade-off scheme and 0.278 under the regional pool, so the trade-off scheme remains closer than the regional pool and is about as close as the neutral fit.

Figure S3 shows the stems summed over the 84 species in each class, the seedling class and the seven censused classes, for the census, for each scheme’s closed form and for each scheme simulated with the environmental noise on. In the closed forms the trade-off scheme’s censused classes run from 0.89 (class 3) to 1.60 (class 8) times the census, and the regional pool’s from 0.70 (class 2) to 3.57 (class 6). With environmental noise the regional pool changes little (large-tree share 26.3%). Under the trade-off scheme every censused class falls by a similar factor, so the proportions among the classes are nearly unchanged (large-tree share 13.5%). In the simulations of Section S7 the saplings (classes 2 to 4) fall to 0.65 of the census and the large trees to 0.84 times the observed. The seedling class, which has no cap in the model, is 1.87 times the census in the closed form and 1.7 in those simulations under the trade-off scheme, and 0.43 and 0.44 under the regional pool. The simulated values in the figure come from a separate set of runs, from one of the two sets of initial conditions. Under the trade-off scheme their seedling, large-tree and censused totals lie 6%, 5% and 4% below those of Section S7, with the seedling class at 1.63 times the census. This is more than the two sets of runs of Section S7 differ from each other, though within 2.1 bootstrap standard deviations of one set. We therefore show the figure for the proportions among the classes, and the totals quoted in the main text are those of Section S7.

On all censused stems (at least 10 mm dbh) rather than on large trees, the rank correlation of stationary with census abundance falls to 0.339 under the trade-off scheme and 0.282 under the regional pool, with the identity-line $R^2$ strongly negative under both (Table S11, lower table). These stems are mainly saplings, 89.19% of the censused stems, belonging mostly to understory species that rarely or never reach 10 cm dbh. Within classes 2, 3 and 4 the trade-off scheme’s stationary totals are close to the census (1.02, 0.89 and 1.02 times the census) but its rank correlations are low (0.20, 0.35 and 0.45). The error enters with the recruits and not in the matrices’ transfer between classes. Started from the observed seedling composition, our matrices order the census saplings at a rank correlation of 0.79 and the large trees at 0.72, although at that composition our seedling promotion rates would have to be scaled up 1.3-fold to supply the observed sapling stand.

The regional pool’s one prediction beyond the regional shares is that abundance relative to regional share rises with the local gain. By Eq. (S3) the gain that would reproduce a species’ observed large-tree abundance under the regional pool is proportional to that abundance divided by its regional share. Across the 79 species whose matrices reach the large-tree classes this required gain falls with the gain computed from the matrix. The log-log slope is $-0.18$ (90% CI $-0.31$ to $-0.05$), where a slope of one is expected if the computed gains acted as modeled. A large local gain under shared recruitment thus does not make a species common at Barro Colorado. The slope lies 16 standard errors below one. With seedling survival and promotion set equal across species it is $-0.11$ (90% CI $-0.27$ to $+0.04$), still 13 standard errors below one, so the result holds without the seedling rates, the least certain entries of the matrices. The gain does mark the direction of recent change, since between 1985 and 2015 the large-tree shares of 53 of the 79 species moved toward the regional pool’s stationary composition $p_s G_s$ (two-sided binomial $P = 0.003$). The movement has a rank correlation of $+0.58$ (90% CI $+0.44$ to $+0.69$) with $\log G_s$. At the observed rate the gap would close with an $e$-folding time of some three centuries. We detect no movement toward the trade-off composition. The census therefore cannot separate arrivals that offset the gain from a stand still centuries from the regional pool’s stationary state. Two further measurements could separate these explanations, since the composition of arriving seed relative to regional share should fall with the local gain under the trade-off scheme, and the stationary composition of a second forest with published matrices should rise with it under the regional pool. Removing the regional pool's predicted dominant does not repair its absolute abundances. Without that species the large-tree total is 1.07 times the observed, but another long-lived species takes its place.

The seedling census (Section S2) tests the model's stationary class 1 (Table S13, closed form). Under the trade-off scheme the seedling class holds 2.09 million stems, 1.9 times the census mean (1.5 to 2.3 times the single censuses), and under the regional pool it holds 0.48 million, 0.43 times the census mean. In the simulations with environmental noise of Section S7, which the main text quotes, the ratios are 1.7 and 0.44. Both schemes lie within the factor of three required in Section S7. Each arriving seed leaves $1/(d\mu_s)$ seedlings standing. Because the trade-off scheme weights arrivals toward species whose seeds yield few large trees, the censused stand stays at capacity only if a larger share of candidate recruits is admitted each step ($d = 0.025$ in place of 0.158), and this smaller $d$ makes the seedling class larger. As shown above, a lower arrival rate cannot remove the excess, since it removes at most the seedlings of the immigrant seeds, the fraction $d$ of the class. *Gustavia superba* alone holds 38% of the class (53% with the noise on), and without it the class is 1.2 times the census mean. Its promotion rate is estimated from 12 promotions among 3,335 seedling-intervals at risk (Section S2). The model's predictions for censused stems do not involve the size of the seedling class. Neither scheme reproduces the seedling abundances of individual species. The trade-off scheme places 36 of the 84 species within a factor of three of their observed mean and the regional pool 29, with rank correlations of 0.26 and 0.00. The observed seedling composition resembles neither the regional composition (rank correlation $-0.02$, whereas the regional composition has a rank correlation of 0.72 with observed large-tree abundance) nor the plot's large-tree composition ($+0.23$). The model's seedling class has no density dependence, and the observed composition may instead reflect conspecific density dependence in seedling survival (Comita et al., 2010).

### Dependence on the normalization and on the sampled data

Unlike the rank order, absolute abundances and the identity of the leading species depend on the normalization of every matrix to growth rate one. Under that normalization each species' growth rate, once the stand is full, is 0.0003 to 0.03 below replacement per interval, and its stationary abundance is its immigrant supply divided by that gap. Raising one species' fecundity until its growth rate is 1.005 increases its abundance under the regional pool by a median factor of 4.1 and in seven of the 84 cases makes it the predicted dominant, whereas a half-percent error in its survival with fecundity rescaled to growth rate one changes its abundance by a median factor of only 1.12. Across these perturbations the rank correlation stays between 0.64 and 0.69. The census data cannot establish that the growth rates are equal at that precision. We redrew every transition probability of classes 2 to 8 from a multinomial at its census sample size and solved the closed form again, 500 times under each scheme, without renormalizing. Species' long-run growth rates then scatter with a median standard deviation of 0.0065, comparable to the gaps below replacement, and about 43 of the 84 species are self-sustaining in a typical draw. The predicted dominant differs from the one at the estimated rates in 99% of draws under the trade-off scheme and 88% under the regional pool. The identity-line $R^2$ is negative in every draw (median $-3.0$ under the trade-off scheme, against $+0.494$ at the estimated rates), and the large-tree total ranges several-fold either way. The rank correlation with the census changes little, with a median of 0.70 (5th to 95th percentile 0.65–0.74) under the trade-off scheme and 0.65 (0.60–0.68) under the regional pool, against 0.727 and 0.672 at the estimated rates. The decadal forecasts are unaffected, since the normalization enters them only through fecundity, which does not reach the large-tree classes in two steps (Section S3). The rank order and the forecasts are therefore the model’s tested predictions. The total, the $R^2$, the dominant and the abundance distribution are consequences that hold at exactly equal long-run fitness.

Table S12 shows how far the long-run values depend on the particular trees and the particular regional plots that were sampled, one source of sampling error at a time. In one set of draws we resampled with replacement every tree and seedling behind the species’ matrices, all records of an individual together, and rebuilt the matrices, each again normalized to growth rate one, and the arrival rate. In the other we resampled the 28 regional plots with replacement and recomputed the regional shares. There are 1,000 draws of each kind for the closed form and 24 of each for the simulations with the environmental noise on. The table gives, for the 84 species under both schemes, the value from the original data and the 90% range of each set of draws for the rank correlation with the census, the identity-line $R^2$, the large-tree total and the seedling pool. The order of species is stable under both kinds of draw. All the 90% ranges of the trade-off scheme’s rank correlation lie between 0.647 and 0.747 and those of the regional pool’s between 0.631 and 0.707, and over these ranges the trade-off scheme’s $R^2$ is positive and the regional pool’s negative. In the simulations, which give the values of the main text, the trade-off scheme’s $R^2$ and large-tree total vary about as much when the trees are redrawn as when the plots are. Its seedling pool varies more with the trees than with the plots, in the simulations and in the closed form. In the closed form its $R^2$ and rank correlation vary mainly with the plots, because its large-tree composition is then the regional shares themselves.

In Table S12 tied abundances are given equal rank, and with ties so treated the trade-off scheme’s closed-form $\rho$ is 0.728, against 0.727 in Table S11. In the simulations each draw is the mean of replicate runs and its range includes their scatter, which for the trade-off scheme’s $\rho$ under redrawn trees is as large as the spread across draws. With 24 draws the ends of a simulated range are themselves uncertain, most of all the upper end of the trade-off scheme’s seedling range under redrawn trees (sampling error 4.7). In the closed-form draws a species left with no promoted seedling either has its seedlings drawn again or takes its promotion rate from a posterior distribution, and the range spans both treatments, of which the simulations use the first. A redrawn list holds fewer distinct plots than the real one and usually leaves more species without a regional share, so the plot ranges of $\rho$ and $R^2$ under the trade-off scheme extend further below the value than above it.

### Time to reach the stationary state

Every simulation starts far from the stationary state, with the censused stand at 1.7 to 2.3 times its capacity. The main runs use two sets of initial conditions that differ in species composition, one even across species and the other spread at random over orders of magnitude, with each species at the stable stage structure of its matrix. We judge stationarity from the simulated abundances alone, without reference to the closed form or to the census. Each run of 12,000 steps is divided into 30 windows of 400 steps, and in each window the number of large trees of every species is averaged over the window and over the 36 replicates of each set. A window meets the drift test for a set when, across species, the change in that mean from the previous window, in $\log_{10}$ of the count plus one, has a median below 0.03 and a ninetieth percentile below 0.10. It meets the agreement test when the two sets’ means, species by species, differ by a median of less than 0.06 in the same units, have a rank correlation of 0.98 or above, and differ by more than twice the bootstrap standard error over replicates for at most 15% of the species. We call the state stationary from the first window at which both tests are met, for both sets, in that window and in the one before it. The settling time is the second window of that pair, so it is resolved to one window of 400 steps. All schemes reached stationarity before window 25, the latest at which we would have accepted a run as converged.

The stationary abundances are the means over the last five windows, with both sets pooled. Under the regional pool the settling time in the main runs is window 6 (2,400 steps, 12,000 model years), the latest among the schemes run in this way under the stem-count capacity and the basal-area capacity of Section S6. As a check of the lottery at a higher arrival rate we also simulated a scheme that weights arrivals by each species’ demographic variance (Jops et al., 2025) at 52,927 seeds per interval, which we call the demographic-variance weighting. The trade-off scheme was simulated in the same way in 4 runs of 36 replicates, one from each of the four kinds of start described below.

Linearized at the mean-field fixed point, the 672-dimensional Jacobian of the deterministic dynamics is block-diagonal in the species, one block $U_{d,s} + F_s$ per species, plus a rank-one coupling through the lottery. Its eigenvalues therefore interlace the blocks’ leading eigenvalues $\rho_s$, and the slowest mode is that of the slowest block, pulled below its $\rho_s$ by less than $5 \times 10^{-6}$. To first order in $d$, $1 - \rho_s = d q_s / T_s$, with $T_s$ the species’ generation time (29 to 478 years across species) and $q_s$ the probability that a seedling dies before its promotion draw, so the slowest e-folding time is $\max_s T_s / (d q_s)$. It is set by the life history of the species held least far below replacement, the one with the largest $T_s / q_s$ (487 years, with $q_s = 0.98$), which is the same species under every scheme. It is also set by $d$, hence by $M$, $w_s$ and the capacity through Eq. (S4), although the composition depends on neither $M$ nor the capacity. The first-order form is within 0.024 ($\log_{10}$ units) of the exact eigenvalue, and the ten slowest modes follow $1 - \rho_s(d)$ to within 0.026 under the regional pool and 0.005 under the trade-off scheme. The e-folding time is 2,900 model years under the regional pool ($d = 0.158$), 7,400 under the demographic-variance weighting ($d = 0.065$) and 19,000 under the trade-off scheme ($d = 0.025$). With zero weight for the species absent from the regional plots, as in the simulations, the last is 20,200. Along a series of arrival rates $M \times \{0.5, 1, 2, 4\}$ the e-folding time scales as $d^{-1.01}$ to $d^{-1.13}$ (5,500, 2,900, 1,600, 1,000 years under the regional pool).

Under the regional pool the simulations reach the stationary state in 10,000 to 12,000 years from each of the four kinds of start we tried (Table S14). Under the trade-off scheme they reach it within 16,000 years from a stand above its capacity and in 38,000 to 42,000 years from an empty plot, because the species of the slowest modes then have to fill from zero. The settling time under the trade-off scheme therefore depends on the start by a factor of about three, with and without demographic noise, whereas under the regional pool it is the same to within one window. With demographic noise, we measured this dependence in 8 further sets of 36 replicates under the regional pool and the demographic-variance weighting, and 4 under the trade-off scheme, run from four kinds of start: the two of the main runs with new random seeds, a flat random composition at twice the capacity, and an empty plot filled by immigration alone. The stationarity test was applied to each pair of starts, 12 pairs under the first two schemes and 6 under the trade-off scheme. Under the regional pool every pair settles at most one window before the main runs. Under the demographic-variance weighting the pairs settle in 10,000 to 22,000 years, and those that include the empty plot are the last. Without demographic noise, we iterated the deterministic dynamics for 12,000 steps from 260 initial compositions per scheme of eight kinds (the two of the main runs, flat random compositions at one, two and four times the capacity, one species at ninety percent of the stems, the empty plot, and the seven censuses themselves with their mean). The same test was applied to pairs of these, with the bootstrap condition, which has no meaning without sampling noise, replaced by a limit of $0.06 \log_{10}$ units on each species’ difference.

The settling time is not a fixed multiple of the slowest e-folding time, because the slowest mode is concentrated in one species and has little weight in the censused classes. The species held least far below replacement holds 0.74 (regional pool) and 0.41 (trade-off scheme) of the weight of the mode’s right eigenvector, and only 0.061 and 0.029 of that weight lies in the censused classes. A mode of e-folding time $\tau$ moves the fraction $1 - e^{-400/\tau}$ of its displacement per window, so the drift test detects it only while its displacement exceeds $0.03 / (1 - e^{-400/\tau})$ in $\log_{10}$ units, which is

0.06 under the regional pool and 0.32 under the trade-off scheme. Under the regional pool the runs start $0.73 \log_{10}$ units from the fixed point in the median species, and the ten slowest modes have e-folding times of 1,400 to 2,900 years. The drift test should then be met once the displacement has decayed to the detectable level, after $\tau_1 \ln(\Delta_0/\epsilon)$ and one window more, which is 5.0 windows from the even start and 4.0 to 5.0 from the random set. The simulations meet it there, at windows 5 to 6, after 4.1 e-folding times, and the deterministic dynamics iterated from the same starts meet it at window 5.

Under the trade-off scheme the slowest mode is 6.9 times slower (10.1 windows against 1.5), and the starts above the capacity displace it little. In the even start the slowest species lies $0.08 \log_{10}$ units from its fixed point, below the detectable level, so the drift test is decided by the faster modes of the other species. The median species' change per window falls by a factor 0.73 per window, an effective e-folding time of 6,500 years, and the deterministic dynamics iterated from the same starts meet the drift test at window 7 (14,000 years). With demographic noise the drift test is met at windows 7 to 8, after 0.7 to 0.8 of an e-folding time of the slowest mode, and for these starts the drift test alone sets the settling time. At window twelve the slowest species is still $0.04 \log_{10}$ units above its fixed point and still decaying with the e-folding time of the slowest mode. In runs from the empty plot the species of the slowest modes fill from zero and approach their stationary abundances from below, whereas from a start above the capacity they approach from above. The difference between the two kinds of start in those species therefore decays at the rate of the slowest modes (e-folding times of 10,000 to 20,200 years). From the empty plot the drift test is met at window 6, but the agreement test is not met until at most fifteen percent of the species differ by more than twice the replicate standard error, and the species of the slowest modes are the last to agree. With demographic noise this happens at windows 19 to 21, after 1.9 to 2.1 e-folding times. The deterministic difference between the empty plot and the even start reaches the same condition at window 17, when twice the replicate standard error is set to $0.035 \log_{10}$ units, its median in the simulations under the demographic-variance weighting at their settling time.

The deterministic test gives a similar range of settling times (Table S14), and under the trade-off scheme its drift condition alone is met between windows 6 and 8. Its fixed limit is stricter than the bootstrap condition for rare species of the slowest modes, whose slow drift the bootstrap condition tolerates. From starts above the capacity the deterministic pairs therefore settle later under the trade-off scheme (28,000 to 38,000 years for the two starts of the main runs, and window 14 for the flat random start at twice the capacity paired with the even start). Under the demographic-variance weighting they settle at a median window of 9, against window 5 at the earliest with demographic noise. The main text gives the settling times with demographic noise.

Whether the stand at Barro Colorado has stood under a constant immigration stream for as long as these settling times is not known. Late in the last glacial period lowland Panama was cooler and drier, with montane trees well below their present range, while lowland forest taxa persisted in communities with no modern analog, and the Holocene opened with an abrupt warming (Bush and Colinvaux, 1990; Bush et al., 1992); the regional pool itself assembled over the same interval, and the plot's composition at any earlier time is unrecorded.

|  | trade-off scheme |  | regional pool |  |
| --- | --- | --- | --- | --- |
|  | closed form | simulated | closed form | simulated |
| rank correlation with the census, $\rho$ | 0.727 | 0.714 | 0.672 | 0.675 |
| identity-line $R^2$ of log abundance | +0.494 | +0.362 | -0.715 | -0.726 |
| large-tree total over the census | 1.23 | 0.84 | 2.41 | 2.30 |
| Kolmogorov distance from the census | 0.127 | 0.215 | 0.287 | 0.278 |
| seedling pool over the seedling census | 1.9 | 1.7 | 0.43 | 0.44 |

Table S10: **Long-run values from the closed form and from the simulations with environmental noise.** Stationary state of the 84 species under the two immigration schemes. The closed form is Eq. (S3), with the environmental noise off. The simulated values have the environmental noise at its estimated amplitude and are means over the last five windows of 36 replicates from each of two sets of initial conditions, run for 60 windows of 400 steps (Section S7). The stationarity test was first met at window 9 under the trade-off scheme and 6 under the regional pool. The rank correlation and $R^2$ are those of stationary with census large-tree abundance across species, the Kolmogorov distance is that between the two species abundance distributions, and the totals are the model’s over the census mean and the seedling-census mean.

| *Stems by censused class relative to the census mean, and the share standing in classes 5 to 8* |  |  |  |  |  |  |  |  |
| --- | --- | --- | --- | --- | --- | --- | --- | --- |
|  | class 2 | class 3 | class 4 | class 5 | class 6 | class 7 | class 8 | % in 5 to 8 |
| census mean (stems) | 80,071 | 66,597 | 32,360 | 13,727 | 5,509 | 2,080 | 377 | 10.81 |
| trade-off scheme / census | 1.02 | 0.89 | 1.02 | 1.12 | 1.57 | 1.02 | 1.60 | 13.31 |
| regional pool / census | 0.70 | 0.79 | 1.22 | 2.20 | 3.57 | 1.00 | 1.13 | 26.09 |

| *The species comparison made on all censused stems, classes 2 to 8* |  |  |  |  |  |  |  |
| --- | --- | --- | --- | --- | --- | --- | --- |
| weighting | $\rho$, classes 2 to 8 | | | $R^2$ of log | sd ratio, | | $\rho$, classes |
|  | full | rare | common | abundance | pred./obs. |  | 5 to 8 |
| trade-off scheme, $w_s \propto p_s/G_s$ | 0.339 | +0.11 | +0.23 | -2.28 | 1.74 | | 0.727 |
| regional pool, $w_s \propto p_s$ | 0.282 | +0.16 | -0.15 | -7.78 | 2.35 | | 0.672 |

| *Share of each species’ censused stems in classes 5 to 8, predicted against observed (79 species)* |  |
| --- | --- |
| median predicted share | 0.140 |
| median observed share | 0.139 |
| predicted over observed, quartiles across species | 0.60 to 1.48 |
| rank correlation, predicted with observed | 0.77 |

Table S11: **Size structure of the stationary stand under the two immigration schemes.** Closed form of Eq. (S3) for the 84 species at Barro Colorado, at 26,499 immigrant seeds per interval. The upper table gives the census-mean number of stems in each censused diameter class (class 2 from 10 mm, class 5 from 79.4 mm dbh) and each scheme’s stationary number as a ratio to it. The capacity fixes the total of classes 2 to 8 at the observed value, so an excess among large trees is offset by a deficit among saplings. The lower table first gives the species comparison on all censused stems: the rank correlation over all species and over the rarer and commoner halves, the identity-line $R^2$ on log abundance and the ratio of predicted to observed standard deviation of log abundance across species, with each scheme’s large-tree rank correlation beside them. Its last rows compare each species’ predicted share of its censused stems in classes 5 to 8, a property of its matrix alone, with the observed share, over the species whose matrices reach those classes.

|  |  | trees redrawn |  | regional plots redrawn |  |
| --- | --- | --- | --- | --- | --- |
|  | value | 90% range | draws | 90% range | draws |
| *Simulated with the environmental noise on* |  |  |  |  |  |
| trade-off scheme, $w_s \propto p_s/G_s$ | | | | | |
| rank correlation with the census, $\rho$ | 0.714 | 0.705 to 0.723 | 24 | 0.647 to 0.735 | 24 |
| identity-line $R^2$ of log abundance | +0.362 | +0.100 to +0.399 | 24 | +0.074 to +0.387 | 24 |
| large-tree total over the census | 0.84 | 0.65 to 0.93 | 24 | 0.76 to 1.01 | 24 |
| seedling pool over the seedling census | 1.7 | 1.4 to 11 | 24 | 1.4 to 2.5 | 24 |
| regional pool, $w_s \propto p_s$ | | | | | |
| rank correlation with the census, $\rho$ | 0.675 | 0.639 to 0.682 | 24 | 0.631 to 0.707 | 24 |
| identity-line $R^2$ of log abundance | −0.726 | −0.863 to −0.689 | 24 | −1.104 to −0.684 | 24 |
| large-tree total over the census | 2.30 | 2.02 to 2.46 | 24 | 2.05 to 2.63 | 24 |
| seedling pool over the seedling census | 0.44 | 0.40 to 0.49 | 24 | 0.38 to 0.58 | 24 |
| *Closed form, environmental noise off* |  |  |  |  |  |
| trade-off scheme, $w_s \propto p_s/G_s$ | | | | | |
| rank correlation with the census, $\rho$ | 0.728 | 0.727 to 0.729 | 1,000 | 0.668 to 0.747 | 1,000 |
| identity-line $R^2$ of log abundance | +0.494 | +0.471 to +0.495 | 1,000 | +0.185 to +0.509 | 1,000 |
| large-tree total over the census | 1.23 | 1.15 to 1.29 | 1,000 | 1.09 to 1.40 | 1,000 |
| seedling pool over the seedling census | 1.9 | 1.6 to 4.6 | 1,000 | 1.4 to 2.5 | 1,000 |
| regional pool, $w_s \propto p_s$ | | | | | |
| rank correlation with the census, $\rho$ | 0.672 | 0.645 to 0.684 | 1,000 | 0.635 to 0.695 | 1,000 |
| identity-line $R^2$ of log abundance | −0.715 | −0.912 to −0.678 | 1,000 | −1.096 to −0.587 | 1,000 |
| large-tree total over the census | 2.41 | 2.13 to 2.81 | 1,000 | 2.18 to 2.74 | 1,000 |
| seedling pool over the seedling census | 0.43 | 0.39 to 0.48 | 1,000 | 0.35 to 0.54 | 1,000 |

Table S12: **Long-run values when the trees or the regional plots behind the model are resampled.** 84 species and 28 regional plots. The value is the estimate from the original data, which is the mean of the simulations of Section S7 in the upper block and the closed form of Eq. (S3) in the lower. When the trees are redrawn, every tree and seedling behind the species' matrices is resampled with replacement, all records of an individual together, and the matrices, their normalization to growth rate one and the arrival rate are rebuilt, with the regional shares as observed. When the regional plots are redrawn, the plots are resampled with replacement and the regional shares recomputed, with the matrices as observed. Ranges run from the 5th to the 95th percentile of the draws, each for one source of sampling error alone. The rank correlation $\rho$ is that of stationary with censused large-tree abundance across species, with tied abundances given equal rank. $R^2$ is taken about the identity line on log abundance. The large-tree total and the seedling pool are the model's totals over the census mean and the seedling-census mean. With 24 draws the simulated ranges are rough (see text).

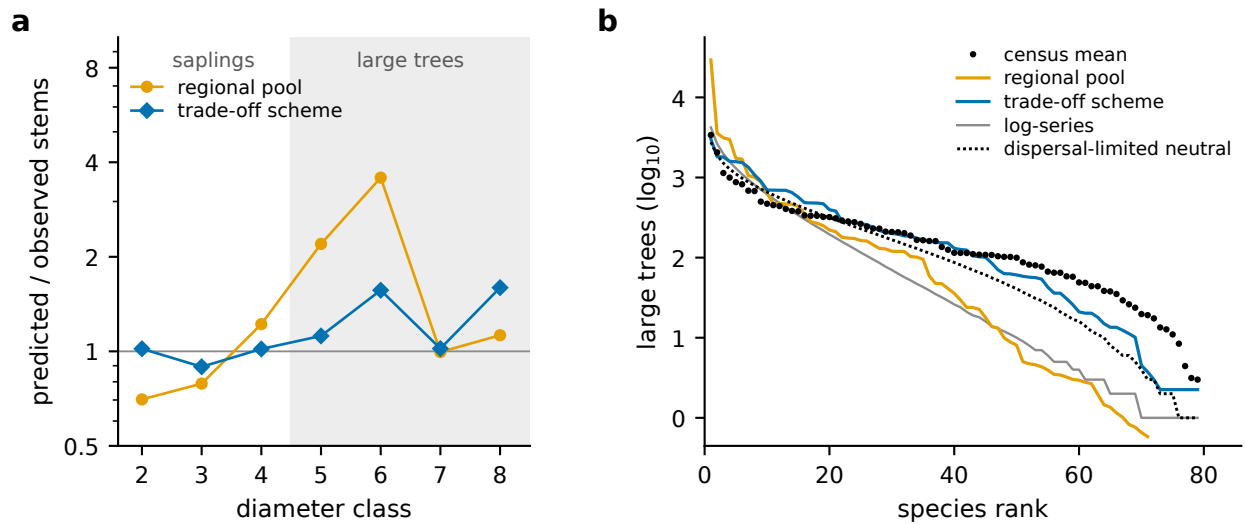

Figure S2: **Size classes and species abundance distribution of the stationary stand under the two immigration schemes.** Closed form at 26,499 immigrant seeds per interval, with the nine species absent from the regional plots at the smallest sampled share. (a) Stationary stems in each censused diameter class, classes 2 to 8, relative to the observed number. The large-tree classes are shaded, and the censused total is fixed at the observed value (Table S11). (b) Ranked log<sub>10</sub> abundance of the 79 species that reach the large-tree classes in the census (points), under the trade-off scheme and the regional pool, with a log-series and Hubbell’s dispersal-limited neutral model fitted to the census abundance distribution of the same species. The Kolmogorov distances from the census on log abundance are 0.127 (trade-off scheme), 0.287 (regional pool), 0.37 (log-series) and 0.18 (neutral model).

|  | seedling census<br>2001–2018, mean<br>(range over 14<br>censuses) | trade-off scheme,<br>stationary class 1 (ratio<br>to the census mean) | regional pool,<br>stationary class 1<br>(ratio to the census<br>mean) |
| --- | --- | --- | --- |
| 84 species, millions on the plot | 1.12 (0.92–1.37) | 2.09 (1.9; by census<br>1.5–2.3) | 0.48 (0.43; by census<br>0.34–0.52) |
| 84 species, per m<sup>2</sup> | 1.8–2.7 | 4.2 | 1.0 |
| 84 species, share of all live<br>seedlings (413 species found), % | 67–75 |  |  |
| fraction of candidate recruits<br>not admitted, $d$ | | 0.025 | 0.158 |
| species within a factor of three<br>of their census mean |  | 36 of 84 | 29 of 84 |
| species above, below that band |  | 25, 23 | 16, 39 |
| rank correlation with observed<br>seedling abundance, across<br>species |  | 0.26 | 0.00 |

Table S13: **The stationary seedling class compared with the seedling census.** The model’s stationary class 1 is the closed form of Eq. (S3) under the two immigration schemes, environmental noise off. The observed values count live free-standing woody stems at least 20 cm tall and under 1 cm dbh of the 84 modeled species in the 19,313 plots of 1 m<sup>2</sup> at each of 14 censuses, scaled to 50 ha. Ratios are model over the observed 14-census mean, and “by census” gives the range of that ratio over the individual censuses.

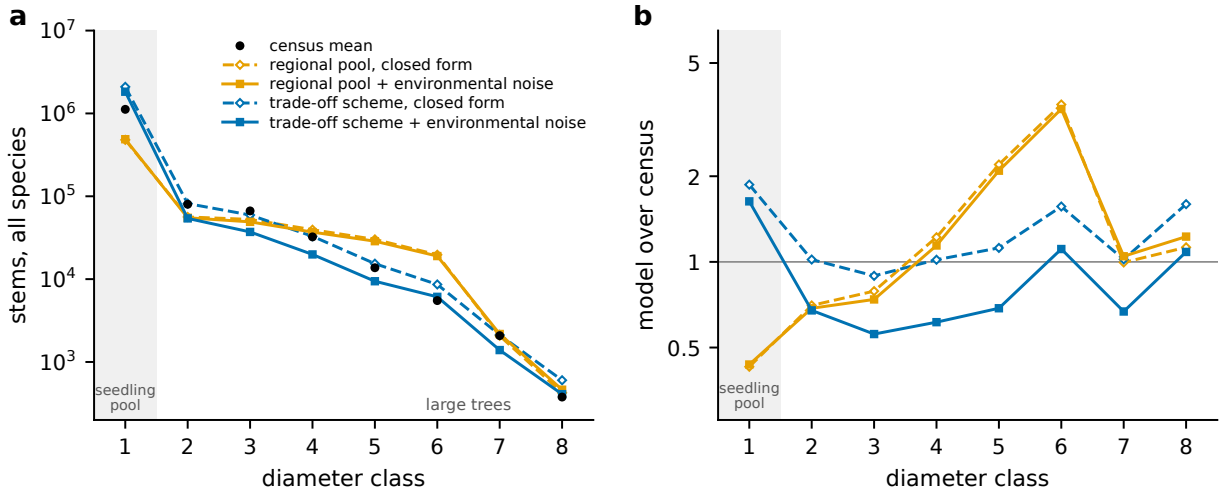

Figure S3: **Stems per size class at the stationary state, census and model.** Stems summed over the 84 modeled species in each diameter class, for the census and for the two immigration schemes with and without the environmental noise. Class 1 is the seedling class (shaded), which has no cap in the model and is measured by the seedling census of Section S2. Classes 2 to 8 are the censused stand, whose total cannot exceed the capacity, and classes 5 to 8 are the large trees. The census values are the seven-census mean for classes 2 to 8 and the seedling-census mean for class 1. Dashed lines and open symbols show each scheme’s closed form (Eq. (S3), noise off, at 26,499 immigrant seeds per interval). Solid lines and filled symbols show the same scheme simulated with the environmental noise at its estimated amplitude, as in Section S7 with every class recorded: 36 replicates from the even set of initial conditions, 60 windows of 400 steps, each class averaged over the last 5 windows. The simulated totals lie a few percent below those of Section S7 (see text). (a) Stems per class, logarithmic axis. (b) Model over census in each class, logarithmic axis about one.

| scheme | with demographic noise |  | deterministic |  |
| --- | --- | --- | --- | --- |
|  | window | years | window | years |
| regional pool | 5 or 6 | 10,000 to 12,000 | 4 to 6 (5) | 8,000 to 12,000 |
| demographic-variance weighting | 5 to 11 | 10,000 to 22,000 | 5 to 22 (9) |  |
| trade-off scheme |  |  | 8 to 21 | 16,000 to 42,000 (30,000) |
| starts above the capacity | up to 8 | up to 16,000 |  |  |
| pairs with the empty plot | 19 to 21 | 38,000 to 42,000 |  |  |

Table S14: **Time to reach the stationary state from different initial conditions.** Window at which pairs of starts first meet the stationarity test, and the corresponding model years (windows of 400 five-year steps). The columns with demographic noise are from simulations of 36 replicates from four kinds of start. The deterministic columns are from the mean-field dynamics iterated from 260 initial compositions per scheme, with the bootstrap condition of the test replaced by a fixed limit. Medians are in parentheses, and under the regional pool the median window is the same for every kind of start. Rows without a qualifier cover all pairs of starts, and an empty cell means that the value was not separated in that way.

### S5 Environmental noise estimated from the transition data

Environmental variation enters the model as one random factor $g_t$ per census interval, the environmental noise, which multiplies a set $T$ of transitions, the same set in every species,

$$[A_s]_{ij} \rightarrow g_t [A_s]_{ij}, \quad (i, j) \in T, \\ g_t = \max(0, 1 + x_t), \quad x_t \text{ Gaussian}, \quad \mathbb{E}[x_t] = 0, \quad \text{sd}(x_t) = \sigma_g, \quad \text{corr}(x_t, x_{t+\Delta t}) = e^{-\Delta t/\tau_g}, \quad (\text{S5})$$

with unpromoted stems staying in their class. Here $A_s$ is the species' projection matrix (Eq. (S2)), $\sigma_g$ the amplitude of the factor, the standard deviation of $x_t$ and so of $g_t$ between intervals, and $\tau_g$ its memory, the correlation time of $x_t$. In this form the factor is shared by all species. The species-specific alternatives in the model comparison of Table S17 (M4F, M4R) give each species its own effect on the same transitions. At Barro Colorado $T$ is the three sapling promotions, out of classes 2, 3 and 4. We estimate the amplitude from the censuses and assume the memory.

The data are the plot's censuses of 1982–2015, seven intervals, the first of 3.4 years and the rest of five. For each of the 84 species and each interval they give nine rates: three sapling promotions (out of classes 2, 3 and 4, from 10 mm dbh to the large-tree threshold of 79.4 mm), three tree promotions (out of classes 5, 6 and 7), mortality in classes 2 to 4 and in classes 5 to 8, and recruitment across 10 mm per standing stem. A species' deviation in an interval is the complementary log-log of its promoted or dead fraction, adjusted for the interval's length and centered on its seven-interval mean (for recruitment, the centered log rate per stem per year). It is a log hazard ratio, close to $\ln g_t$ of Eq. (S5).

The component common to all species in a rate is the set of centered interval effects of a binomial regression (Poisson for recruitment) with species and interval terms (Table S16). In each rate this component exceeds its sampling floor, the 95th percentile expected with no environmental noise, and in the sapling promotions it carries most of the between-interval variance. The species-level statistics behind main text Fig. 2 are in Table S15. Sapling growth varies in common across species, with 99% of the 175 species-by-promotion series correlated positively with the component common to the other species and 69% above the seven-point null. The median species follows the common component at close to full amplitude. Mortality in classes 2 to 4 varies within a species as much as sapling growth does, but not in common (median $r + 0.44$, 27% above the null).

Table S17 compares structures of the environmental noise fitted to the species-by-interval counts of all nine rates, 4,228 observations (4.0 million stem-intervals). We fit them on the seven intervals and on the six from 1985, because the 1982 census measured small stems by a different method. The Bayesian information criterion retains a term of some form on each group of rates and rejects only two elaborations, free interval effects for the tree promotions in place of one loading and free species-by-interval effects in place of a variance component. The composite model M6 has the lowest BIC under either sample size and on both seven and six intervals. The data therefore support more structure than one shared effect on sapling growth. The three sapling promotions, for instance, differ in amplitude on seven intervals (M1b), though not on 1985–2015 ($P = 0.46$).

In amplitude, however, the shared effect on sapling growth dominates. It has standard deviation 0.20 over the seven intervals. The tree promotions load $\lambda_{\text{late}} = 0.86$ on it (90% profile CI 0.75–0.97) and, given interval effects of their own, reproduce it (correlation 0.95), so growth follows the same interval effects at every size. The systematic between-interval variance of the sapling promotions divides into 71% common to all species, 10% species-by-interval deviation common to a species' sapling stages ($\sigma_b = 0.075$) and 20% count-level overdispersion. Mortality loads $-0.06$ (classes 2 to 4) and $+0.39$ (classes 5 to 8) on the shared growth effect, under half the growth amplitude, and its separate interval effects (standard deviations 0.09 and 0.16) are only weakly correlated with those of growth. We therefore give the model one factor $g_t$ per interval, common to species, on the three sapling promotions with one amplitude, no factor on mortality and no species-specific response.

Species-specific responses (model M4F) add 498 parameters and raise the log-likelihood by 909, which is 3.6 units of deviance per parameter against 932 for the six common effects. They are favored by Akaike’s criterion but not by the Bayesian criterion (8.35 per parameter), and a single number for their spread, $\sigma_b$, is favored by both. They amount to 10% of the systematic variance, and we use the shared effect alone. We did not make the corresponding comparison for mortality, and test species-level mortality variation by simulation instead (Section S7).

The mortality term tested in Section S7 multiplies each species’ mortality hazard at every step by a lognormal factor of mean one, independent across species and steps. Its log-scale standard deviation is the median species value of Table S15 (0.19 in classes 2 to 4, 0.24 in classes 5 to 8). The term has no persistent direction and, apart from a slight rise in mean survival (the five-year death probability falls by 0.4% of its value), leaves each species’ mean matrix, forecast and stationary abundance unchanged. It adds only to the size of census-to-census changes. Section S7 also tests a mortality factor shared by all species, with one multiplier on the hazard at each step for classes 2 to 4 and one for classes 5 to 8, correlated at 0.58. Converted to amplitudes by the method used below for $\sigma_g$, the common mortality components of Table S16 give log-scale standard deviations of 0.146 and 0.254 for the two groups if the factor has the memory assumed for growth, and 0.095 and 0.167 if it has none. The seven intervals cannot tell these two cases apart.

We estimate the amplitude $\sigma_g$ from the pooled sapling path, the mean of the three sapling promotions’ centered interval effects (dashed line in main text Fig. 2). The three promotions are consistent with one path (the first principal component of their effects carries 93% of the variance, with loadings 0.72, 1.24 and 1.03). The standard deviation of the path over the seven intervals, with the sampling variance removed, is $S = 0.222$. The estimate is the amplitude at which simulated environmental noise, sampled at the census dates with the census numbers at risk, reproduces this spread, and the median of 20,000 simulated statistics equals $S$ at $\sigma_g = 0.318$. The 90% confidence interval is 0.17–0.84, wide at the top because seven serially correlated samples of a slowly varying factor often show much less than its full spread. Without the first interval, whose pooled effect is the largest, $\sigma_g = 0.27$ (0.14–0.71). The results for growth above also hold on 1985–2015, where the tree-promotion loading is 0.75 (0.62–0.88) and the common and species-specific shares of the variance are 80 and 13%.

The memory $\tau_g$ cannot be estimated from seven intervals. The pooled path’s lag-one autocorrelation, +0.32, lies inside the range (-0.34 to +0.58) that a factor with $\tau_g = 18$ yr gives on seven points, and inside the range for all correlation times from 5 to 40 yr. The relative performance of species does persist from one interval to the next. Across 255 species the net rate of gaining trees of at least 100 mm dbh has a rank correlation of 0.33 between consecutive intervals, which decays with a time constant of 17.9 yr (90% CI 10.8–33.2 yr). This persistence does not constrain $\tau_g$, because a factor common to all species moves them together. Records simulated from the model at the census sample sizes show no such persistence (median rank correlation -0.08 to +0.02 at all lags, whatever the imposed correlation time), so the persistence reflects species-specific departures from the mean rates. We set $\tau_g = 18$ yr ($\phi = 0.76$ per five-year step) by assumption and consider 5 to 40 yr, over which the amplitude estimate ranges from 0.23 to 0.43. The climate series of Table S20 all lose their memory within 4.4 yr.

Recruitment across the 100-mm line varies between censuses with a standard deviation of 18% of its mean rate, nearly all of it common to the species. This value, 0.18, is the measured recruitment variability used in the rest of this section and in Section S7. We placed a factor shared by all species on each transition in turn and asked whether it could produce the measured variability of recruitment into the large-tree classes without changing the size of the stand or the rank order of its species (Table S18, Fig. S4). Under the regional pool, for which Table S18 is computed, the variability of recruitment into class 5 reaches the measured value at $\sigma_g = 1.07$ on promotion out of class 3, at 0.30 on promotion into class 5 and at 0.27 on the three sapling promotions together. Only the last two leave the composition and the size of the stand as they are without the noise. On seed production, establishment and promotion out of class 2 the variability levels off below the measured value at all amplitudes up to 1.5, and on survival it stays below while the stand changes in size or rank order. A factor below class 3 fails because its variation is absorbed as stems pass slowly through the sapling classes above. Each of these classes holds a stem for 13 to 25 years, comparable to or longer than the assumed 18-year memory (Fig. S4b).

We repeated the test under the trade-off scheme at the same amplitudes. The three sapling promotions together, the placement used in the model, still reach the measured variability (0.206) and keep the rank order (the rank correlation with the census changes by $-0.016$ relative to the simulation without the noise). They lower the stand to 0.71 of its size without the noise, however, and change the identity-line $R^2$ of log abundances by $-0.124$. This exceeds the limit of five hundredths that we set for a change in composition, so this placement does not preserve the composition under the trade-off scheme. Promotion into class 5 alone reaches the measured variability under that scheme (0.180) with the stand at 0.88 of its size without the noise, the rank correlation changed by $-0.008$ and the $R^2$ by $-0.010$, and so meets the conditions on variability, stand size and composition (Section S7).

The common components of Table S16 distinguish between these two placements. A factor on promotion into class 5 alone, with $\sigma_g = 0.30$ and $\tau_g = 18$ yr and sampled at the census dates and numbers at risk, implies a standard deviation of the common component of that promotion of 0.099 to 0.473 over seven intervals (central 90% of 2,000 draws), 0.088 to 0.448 over six and 0.099 to 0.468 per stem at risk. These ranges contain the census values, but such a factor implies no common component above the sampling floors at the two lower promotions. The three sapling promotions together at $\sigma_g = 0.27$ give 0.10–0.45 out of class 2 and 0.10–0.40 out of class 3, ranges that contain the census values.

Noise on seed production at larger amplitudes (a mean-one lognormal factor, trade-off scheme) only reduces the stand, to 0.14 of its size without the noise, because the seedling pool supplies just 3% more candidates than there are places. The same shortage is why the sapling-growth factor of the model lowers the censused stand under that scheme, to 0.669 of the capacity (Section S7). A shared factor on seed production also cannot supply the variability that the stationary model lacks at intermediate abundance, because its contribution to a species’ mean squared change scales as abundance squared, whereas demographic variance scales as abundance.

The comparisons with traits and climate are exploratory, because each interval-level statistic rests on seven points. Each species’ loading on the common sapling-growth variation (76 species) is compared in Table S19 with nineteen traits, by rank correlation and by partial correlation given log abundance, since the abundant species dominate the common series. The loadings are correlated neither with demographic variance (Jops and O’Dwyer, 2023) nor with stature. The largest correlations, all negative, are with the light-response and fast-growth indices of Rüger et al. (2022) and Wright et al. (2010) and with seedling mortality. The bootstrap intervals of nine of the nineteen rank correlations exclude zero (1.9 expected by chance), and the largest $|\rho|$, 0.49, exceeds the family-wise 95% level of 0.34. Demographic variance and the six quantities computed from our matrices come from the same matrices and form one family. Given abundance, only sapling growth potential, growth in shade and seedling mortality keep negative partial correlations. If synchronized canopy opening drove the common variation, the light-demanding half of the species should follow it more strongly than the shade-tolerant half. With the species split at the median growth response to light, the two halves’ common series are correlated at 0.96 and the light-demanding half’s amplitude is 0.88 (90% CI 0.75, 1.01) times the other’s, which lends no support to a light-mediated driver. The light indices, however, describe contrasts sustained over whole census intervals, whereas a driver would act through the light of particular years.

| rate | $n$ | size: median s.d.<br>[quartiles] | direction: median $r$<br>[quartiles] | $r > 0$<br>(%) | above<br>null (%) | in phase<br>(%) | memory: median $\rho_1$<br>[quartiles] |
| --- | --- | --- | --- | --- | --- | --- | --- |
| promotion 2 → 3 | 76 | 0.19 [0.15, 0.24] | +0.77 [+0.60, +0.85] | 100 | 67 | 88 | +0.02 [-0.13, +0.23] |
| promotion 3 → 4 | 62 | 0.27 [0.22, 0.32] | +0.83 [+0.64, +0.93] | 98 | 73 | 76 | +0.27 [+0.16, +0.36] |
| promotion 4 → 5 | 37 | 0.24 [0.13, 0.32] | +0.77 [+0.64, +0.90] | 100 | 68 | 73 | +0.16 [+0.01, +0.29] |
| promotion 5 → 6 | 13 | 0.12 [0.07, 0.18] | +0.69 [+0.60, +0.83] | 100 | 62 | 85 | +0.04 [-0.07, +0.14] |
| promotion 6 → 7 | 3 | 0.33 [0.28, 0.34] | +0.78 [+0.71, +0.87] | 100 | 67 | 67 | +0.07 [-0.07, +0.13] |
| mortality, classes 2–4 | 75 | 0.19 [0.12, 0.25] | +0.44 [+0.30, +0.68] | 95 | 27 | 75 | +0.20 [-0.11, +0.39] |
| mortality, classes 5–8 | 28 | 0.24 [0.16, 0.31] | +0.60 [+0.49, +0.84] | 93 | 43 | 68 | +0.14 [-0.05, +0.29] |
| recruitment at 10 mm | 82 | 0.41 [0.33, 0.53] | +0.84 [+0.69, +0.92] | 95 | 77 | 66 | +0.33 [+0.27, +0.40] |

Table S15: **Growth and recruitment vary in step across species.** Census-to-census variation of each transition rate within species at Barro Colorado, 1982–2015 (seven intervals; census data). Classes are the eight stages of the published matrices (Jops et al., 2025), class 2 from 10 mm dbh and class 5, the first large-tree class, from 79.4 mm. Statistics are over the $n$ species with at least twenty expected events per interval. Size is the median standard deviation of a species’ seven interval deviations on the log-hazard scale, with sampling variance removed. Direction is the median correlation of a species’ series with the component common to the other species. The next two columns give the percentage of species correlated positively with that component and the percentage above the 95th percentile for an unrelated seven-point series (0.67). In phase is the percentage best aligned with the common component at lag zero rather than one interval ahead or behind (16–29% for an unrelated series). Memory is the median lag-one autocorrelation (–0.62 to +0.36 for seven points with no memory, –0.36 to +0.60 with the memory of the environmental noise). No species qualifies for the promotion out of class 7 (340 events in all).

Seven intervals can neither establish nor exclude a climate driver. None of the nine climate covariates of Table S20 is correlated with the pooled sapling path beyond what so many comparisons on seven points give by chance. The largest $|\rho|$, 0.86 with January-to-April radiation, has family-wise $P = 0.175$ (0.61 and $P = 0.70$ with windows one year later), and its sign (–0.86) is opposite to that expected under light limitation. The path’s rank correlation with the interval midpoint is –0.29 ($P = 0.56$), whereas a steadily changing driver such as rising CO<sub>2</sub> or warming would give a value near plus or minus one.

### Fluctuations with and without the omitted noise terms

The model keeps one term of the environmental noise, the factor shared by all species on the three sapling promotions. It omits the others measured above, among them each species’ year-to-year variation in mortality and the deviations of each species’ sapling growth from the shared factor. In the model the noise perturbs each species’ stage vector away from its stationary vector, and the species’ matrix converts the perturbation into changes in its large trees over the following censuses. We therefore compare the size of the simulated fluctuations with the census; earlier work added environmental noise to neutral models to enlarge these fluctuations (Kalyuzhny et al., 2015; Fung et al., 2016). Section S7 and Table S26 give the full comparison.

Without environmental noise the large-tree total changes over five years by a median 0.0012 under the regional pool, 11 times less than the census’s 0.0127. With the shared sapling-growth factor alone, at its estimated amplitude, the total changes by 0.017 under the regional pool and 0.020 under the trade-off scheme, and recruitment into class 5 varies over seven censuses with a coefficient of variation of 0.21, against 0.18 measured. The mortality term, simulated alone at its measured size, gives a total change of 0.0057 and 0.0068 under the two schemes and a coefficient of variation of recruitment of only 0.02 and 0.03. These are about half the census’s change in the total and a tenth to a sixth of the measured variability of recruitment. Added to the shared factor, the mortality term leaves the total’s change at 0.0176 and 0.0172, so the shared factor sets the size of the community’s fluctuations. A shared factor placed instead on seed production, establishment or the first sapling promotion cannot produce the measured variability of recruitment, because the sapling classes absorb it (Fig. S4).

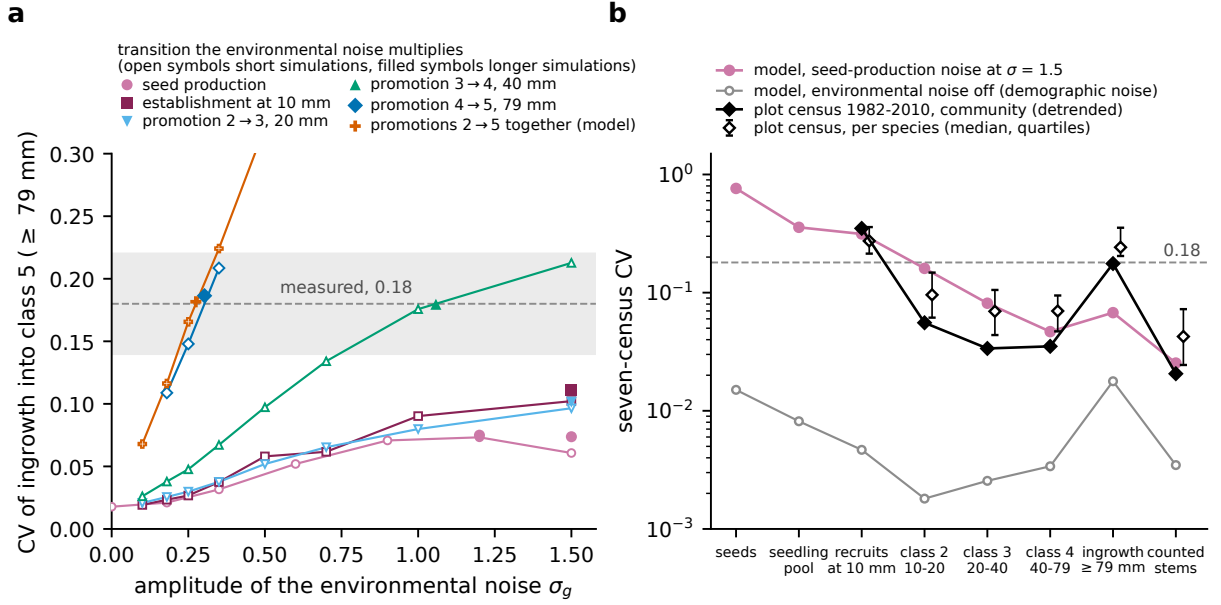

Figure S4: **Recruitment variability against noise amplitude for each transition the shared factor multiplies.** (a) Model: the between-census coefficient of variation of recruitment into class 5, the first large-tree class, over seven censuses, against the amplitude $\sigma_g$ of the environmental noise. Open symbols are short simulations and filled symbols the longer simulations that locate each crossing. The dashed line and band mark the value measured at Barro Colorado (0.18) and the range accepted around it. Promotion into class 5 and the three sapling promotions together cross the measured value at $\sigma_g = 0.30$ and $0.27$, and promotion out of class 3 at $1.07$. Seed production, establishment at 10 mm and promotion out of class 2 level off below it up to $1.5$. (b) The seven-census coefficient of variation of each quantity from seed production up the size classes to the large-tree total, in the model with the noise on seed production at $\sigma_g = 1.5$ (filled circles) and with no noise (open circles), and in the censuses of 1982 to 2010 for the whole community (filled diamonds, detrended) and the median species (open diamonds, quartiles). In model and census the variation present at 10 mm has largely disappeared in the sapling classes, yet census ingrowth at the large-tree threshold varies at the measured 0.18.

| rate | species | sd, all | sd, from 1985 | floor | common (%) | sd<sub>common</sub> | sd<sub>specific</sub> |
| --- | --- | --- | --- | --- | --- | --- | --- |
| promotion 4 → 5, per survivor | 77 | 0.232 | 0.174 | 0.030 | 78 | 0.231 | 0.122 |
| promotion 4 → 5, per stem at risk | 77 | 0.245 | 0.178 | 0.031 | 77 | 0.244 | 0.135 |
| promotion 2 → 3, per survivor | 84 | 0.182 | 0.186 | 0.012 | 86 | 0.182 | 0.073 |
| promotion 3 → 4, per survivor | 84 | 0.284 | 0.176 | 0.018 | 81 | 0.283 | 0.137 |
| classes 2–4 into class 5 | 77 | 0.178 | 0.162 | 0.029 | 36 | 0.177 | 0.237 |
| mortality, classes 2–4 | 84 | 0.090 | 0.094 | 0.011 | 10 | 0.089 | 0.271 |
| mortality, classes 5–8 | 77 | 0.158 | 0.152 | 0.032 | 100 | 0.156 | 0.000 |
| recruitment at 10 mm, per stem | 84 | 0.364 | 0.226 | 0.010 | 47 | 0.364 | 0.388 |

Table S16: **The sapling promotions vary mostly in common across species.** The component of each transition rate common to the species at Barro Colorado, seven intervals from 1982 to 2015 (census data; the bold lines of main text Fig. 2). Species are those with at least five events in at least five intervals. The standard deviations (sd) are those of the centered interval effects of a regression with species and interval terms (binomial, complementary log-log link; Poisson for recruitment), over all intervals and over the six from 1985. Floor is the 95th percentile of the same statistic with no environmental noise (1,000 simulated tables). Common is the share of the interval-to-interval variance of the species’ rates common to them, and the last two columns are the common and species-specific standard deviations behind that share.

For individual species the mortality term does matter. With it, the census’s mean squared five-year change in the two most abundant bands is 1.26 and 1.24 times the model’s under the regional pool, against 1.61 and 1.77 without it. Neither term closes the stationary model’s shortfall at intermediate abundance, where species of 16 to 255 large trees still change 2.6 to 2.9 times as much in the census as in the model with both terms. Started from the 1982 forest, the model with the shared factor alone has the census inside its 95% range in every band in which the census records a change (Fig. S9), though above its mean from 16 trees upward. We did not simulate the species-specific deviations of sapling growth.

| | environmental-noise terms | df | log $L$ | ΔBIC | ΔBIC<br>(stems) | ΔBIC<br>1985–2015 |
| --- | --- | --- | --- | --- | --- | --- |
| M0 | none: the mean rates only | 0 | −31,944 | 25,397 | 25,136 | 11,002 |
| M1 | one effect $a_t$, common to species, on the sapling promotions $2 \rightarrow 3$,<br>$3 \rightarrow 4$, $4 \rightarrow 5$ | 6 | −29,146 | 19,852 | 19,633 | 7,229 |
| M1b | M1 with an amplitude per promotion ($\lambda_{23} = 0.75$, $\lambda_{34} = 1.21$;<br>1.06, 1.02 on 1985–2015) | 8 | −29,022 | 19,619 | 19,414 | 7,244 |
| M1c | interval effects free per sapling promotion | 18 | −28,668 | 18,996 | 18,859 | 7,103 |
| M2 | M1 + the same $a_t$ on the tree promotions $5 \rightarrow 6$, $6 \rightarrow 7$, $7 \rightarrow 8$ | 6 | −29,053 | 19,664 | 19,445 | 7,140 |
| M2b | M1 + $\lambda_{\text{late}} a_t$ on the tree promotions ($\lambda_{\text{late}} = 0.86$) | 7 | −29,050 | 19,668 | 19,455 | 7,137 |
| M2c | M1 + interval effects of their own on the tree promotions | 12 | −29,041 | 19,692 | 19,514 | 7,158 |
| M3 | M2 + the same $a_t$ on mortality in classes 2–4 and 5–8 | 6 | −30,172 | 21,904 | 21,685 | 8,976 |
| M3b | M2b + $\lambda a_t$ on each mortality (−0.06, +0.39) | 9 | −29,005 | 19,594 | 19,395 | 7,026 |
| M3c | M2c + interval effects of their own on each mortality | 24 | −28,400 | 18,510 | 18,414 | 6,084 |
| M1+Mb | M1 + $\lambda a_t$ on each mortality, no tree-growth term | 8 | −29,101 | 19,777 | 19,572 | 7,120 |
| M1+Mc | M1 + own interval effects on each mortality, no tree-growth term | 18 | −28,505 | 18,670 | 18,533 | 6,155 |
| M5s | M1 + the same $a_t$ on recruitment at 10 mm | 6 | −21,804 | 5,167 | 4,948 | 1,850 |
| M5 | M1 + $\lambda_{\text{rec}} a_t$ on recruitment ($\lambda_{\text{rec}} = 1.83$) | 7 | −20,902 | 3,372 | 3,160 | 1,722 |
| M5c | M1 + own interval effects on recruitment | 12 | −20,637 | 2,883 | 2,704 | 1,657 |
| M4F | M1 + species × interval effects on the sapling promotions, free | 504 | −28,237 | 22,192 | 25,386 | 9,320 |
| M4R | M1 + species × interval effects, random ($\sigma_b = 0.099$) | 7 | −28,803 | 19,173 | 18,961 | 6,791 |
| M4Rk | M1 + count-level random effects ($\sigma_c = 0.128$) | 7 | −28,468 | 18,503 | 18,291 | 6,790 |
| M4R2 | M1 + both ($\sigma_b = 0.075$, $\sigma_c = 0.107$) | 8 | −28,459 | 18,495 | 18,289 | 6,752 |
| M6– | the supported shared terms together: $a_t$ free per sapling<br>promotion, own effects on recruitment and on each mortality,<br>$\lambda_{\text{late}} a_{45,t}$ on the tree promotions ($\lambda_{\text{late}} = 0.79$) | 37 | −19,414 | 646 | 640 | 356 |
| M6 | M6– + the species × interval random term ($\lambda_{\text{late}} = 0.76$,<br>$\sigma_b = 0.099$) | 38 | −19,087 | 0 | 0 | 0 |
| Mown | every rate its own interval effects (reference) | 54 | −19,379 | 718 | 828 | 417 |
| Msat | Mown + the free species × interval effects (reference) | 540 | −18,948 | 3,915 | 7,355 | 2,635 |

Table S17: **The Bayesian information criterion favors the composite model M6.** Structures of the environmental noise fitted to the Barro Colorado transition data. Each model is fitted by maximum likelihood to the same 4,228 species-by-interval-by-rate counts (binomial with complementary log-log link for promotion and death, Poisson for recruitment), with one intercept per species and rate (604); df counts the parameters beyond those intercepts. ΔBIC is the criterion less its lowest value in the column, with the number of counts as the sample size and, beside it, the number of stem-intervals at risk. The last column refits the models on the six intervals of 1985–2015 (3,624 counts). M4R2 holds the intercepts and $a_t$ at M4R’s values, so its likelihood is a lower bound.

| transition multiplied<br>by the environmental noise | $\sigma_g$ reaching 0.18 | CV at $\sigma_g=1.5$<br>if never | stand size<br>(no noise =1) | composition<br>unchanged |
| --- | --- | --- | --- | --- |
| seed production | none | 0.039 | 0.87 | no |
| seedling survival (class 1) | none | 0.093 | 0.85 | no |
| establishment into class 2 (crossing<br>10 mm) | none | 0.074 | 0.78 | no |
| promotion out of class 2 | none | 0.090 | 0.88 | no |
| promotion out of class 3 | 1.07 |  | 0.92 | no |
| promotion into class 5 (first<br>large-tree class) | 0.30 |  | 1.01 | yes |
| the three sapling promotions<br>together (used in the model) | 0.27 |  | 0.97 | yes |
| sapling survival, classes 2–4 | none | 0.103 | 1.14 | no |
| adult survival, classes 5–8 | none | 0.033 | 0.55 | no |

Table S18: **Recruitment variability and stand size with a shared factor on each transition in turn.** Model, regional pool. The second column is the amplitude $\sigma_g$ at which the modeled between-census coefficient of variation (CV) of recruitment into class 5 over seven censuses reaches the measured 0.18. Where no amplitude up to 1.5 reaches it, the third column gives the CV attained there. Stand size is the stationary number of censused stems at that amplitude as a fraction of its value without the noise. The last column says whether the long-run composition remains that of the model without the noise, judged by the change in rank correlation allowed in Section S7. On the survival rows the factor multiplies the mortality hazard. The model uses $\sigma_g = 0.32$, which is estimated from the transition rates and not taken from this table.

| trait | source | $n$ | $\rho$ [90%] | given abundance [90%] |
| --- | --- | --- | --- | --- |
| demographic variance | our matrices; Jops and O'Dwyer (2023) | 76 | -0.08 [-0.29, +0.12] | +0.04 [-0.17, +0.23] |
| stature (mean height of the six largest stems, else maximum height), m | Wright et al. (2010); Condit et al. (2020) | 73 | -0.10 [-0.29, +0.11] | -0.12 [-0.30, +0.07] |
| maximum height, m | Rüger et al. (2022) | 76 | -0.07 [-0.26, +0.12] | -0.05 [-0.24, +0.13] |
| wood density, $\text{g cm}^{-3}$ | Wright et al. (2010) | 72 | +0.28 [+0.10, +0.45] | +0.28 [+0.09, +0.44] |
| seed dry mass, g | Wright et al. (2010) | 65 | +0.13 [-0.09, +0.34] | +0.05 [-0.18, +0.27] |
| dry-season moisture association | Condit et al. (2013) | 66 | +0.02 [-0.20, +0.24] | -0.02 [-0.23, +0.19] |
| sapling growth potential RGR95SAP | Wright et al. (2010) | 72 | -0.49 [-0.64, -0.32] | -0.34 [-0.52, -0.13] |
| mortality of the slowest-growing quarter of saplings MRT25SAP | Wright et al. (2010) | 72 | -0.30 [-0.47, -0.10] | -0.08 [-0.27, +0.11] |
| growth response to light (growth in high light over growth in low light) | Rüger et al. (2022) | 75 | -0.39 [-0.56, -0.21] | -0.19 [-0.39, +0.03] |
| mortality response to light (high over low) | Rüger et al. (2022) | 75 | +0.21 [+0.01, +0.39] | +0.06 [-0.14, +0.26] |
| recruitment response to light (high over low) | Rüger et al. (2022) | 75 | -0.39 [-0.56, -0.19] | -0.23 [-0.45, -0.01] |
| growth in shade ($\text{mm yr}^{-1}$ of a 10-mm stem in low light) | Rüger et al. (2022) | 75 | -0.45 [-0.60, -0.28] | -0.32 [-0.49, -0.12] |
| mortality in shade ($\text{yr}^{-1}$ of a 10-mm stem in low light) | Rüger et al. (2022) | 75 | -0.28 [-0.44, -0.09] | -0.06 [-0.27, +0.15] |
| annual survival in the large-tree classes | our matrices | 71 | +0.18 [-0.04, +0.39] | +0.12 [-0.08, +0.31] |
| probability that a 10-mm recruit reaches the large-tree classes | our matrices | 71 | -0.08 [-0.27, +0.12] | -0.14 [-0.31, +0.06] |
| first-class recruits per large tree per year | our matrices | 71 | -0.05 [-0.27, +0.17] | +0.07 [-0.14, +0.26] |
| local gain $G_s$ (large trees standing per arriving seed) | our matrices; Section S4 | 71 | +0.01 [-0.20, +0.21] | -0.08 [-0.27, +0.12] |
| large trees per seedling in the stable structure, $\Omega_s$ | our matrices | 76 | -0.13 [-0.33, +0.07] | -0.14 [-0.33, +0.06] |
| seedling mortality per five-year step, $\mu_s$ | our matrices | 76 | -0.37 [-0.51, -0.20] | -0.27 [-0.42, -0.10] |

Table S19: **Species' loadings on the common sapling-growth variation compared with their traits.** Barro Colorado, 1982–2015 (seven intervals; exploratory); 76 species and nineteen traits. The loading is the inverse-variance mean, over the sapling promotions for which the species has at least twenty expected events per interval, of the slope of its interval deviations on the leave-one-out common series (Table S15). The median loading is 0.89 and its standard deviation across species 0.30, or 0.19 after subtracting sampling variance. $n$ is the number of species with the trait, $\rho$ the Spearman rank correlation with its bootstrap 90% interval (10,000 resamples of species), and the last column the partial rank correlation given the logarithm of mean stems at risk. High and low light are the two reference conditions of Rüger et al. (2022).

| covariate, mean over the interval's calendar years | $\rho$ | $P$ | $\rho$, windows | $P$ |
| --- | --- | --- | --- | --- |
|  |  |  | one year later |  |
| Oceanic Niño Index, mean | −0.32 | 0.498 | −0.50 | 0.267 |
| fraction of months in El Niño conditions | +0.13 | 0.800 | −0.36 | 0.429 |
| fraction of months in La Niña conditions | +0.13 | 0.798 | +0.29 | 0.556 |
| dry-season rainfall | −0.57 | 0.200 | −0.61 | 0.167 |
| annual rainfall | −0.57 | 0.200 | −0.18 | 0.713 |
| solar radiation, January to April | −0.86 | 0.024 | −0.61 | 0.167 |
| solar radiation, annual | −0.79 | 0.048 | −0.50 | 0.267 |
| air temperature | −0.21 | 0.662 | −0.18 | 0.713 |
| dry-season length | +0.39 | 0.396 | +0.29 | 0.556 |

Table S20: **No climate series tracks the common sapling-growth variation beyond chance.** Rank correlations of the common sapling-growth variation with published climate series over the seven census intervals (descriptive only). The series is the pooled sapling path (+0.35, +0.22, −0.26, −0.20, −0.10, −0.06, +0.05 for 1982–85 to 2010–15). $\rho$ is the Spearman rank correlation with each covariate averaged over the interval's nominal calendar years (1982–84, 1985–89, ..., 2010–14) and over windows one year later, since each census took about a year. $P$ is two-sided, over all 5,040 orderings of seven values. The Oceanic Niño Index and its El Niño and La Niña classifications are the NOAA Climate Prediction Center series (14, 17, 20, 14, 13, 13, 6 El Niño months in the seven windows). Rainfall, radiation, temperature and season dates are the Barro Colorado clearing-station summaries of Paton (2024).

### S6 Model variants and sensitivity of results

Each variant in this section is the model of the main text with one element changed. That model has the trade-off scheme, 26,499 immigrant seeds per interval, the published eight stages and a capacity on censused stems, and the stationary results of this section are all for the trade-off scheme. We score each variant on the two main results. The first is the out-of-sample forecast with the rates inferred from the censuses of 1982–1995, scored by the rank correlation of predicted with observed change in large trees one and two censuses ahead over the 252 species-start pairs of Section S3. The second is the stationary large-tree abundance of each species, scored by its rank correlation and identity-line $R^2$ against the census means (Section S4). Table S21 collects the scores, Fig. S5 draws the variants, and Figs. S6 and S7 repeat Figs. 3 and 4 of the main text for a representative set.

The arrival rate is $M = \hat{m}B$. The immigrant fraction of recruits, $\hat{m} = 0.026$, is estimated from the plot itself, as the value at which Hubbell’s neutral model best reproduces the plot’s census-to-census turnover of trees of at least 10 cm dbh. The seed production, $B = 1.01$ million seeds per interval, is the order of magnitude that the fecundity rows give for a stand of this size, since seed rain was not measured. The censuses bound the immigrant fraction from below: 30 of 16,982 recruits belonged to species not previously recorded on the plot (0.0018). Because $\hat{m}$ comes from the fit of a different model and $B$ is nominal, $M$ is known only to its order of magnitude. This is enough for our purposes, because by the closed form (Section S4) the censused composition does not depend on $M$, and only the seedling pool does, through $d$. A better value of $M$ would have to be measured directly.

The eight alternative stage structures rebuild the diameter classes from the census measurements. They are four classes, thirteen log-spaced classes at half the published width, ten log-spaced classes up to the largest stem, equal-width classes with and without an edge at 79.4 mm, finer sapling classes, and two integral projection models on a continuous diameter axis, with parametric or non-parametric vital rates. All keep the seedling class, with its rates inferred as described in Section S2. The fecundity schedule is rebuilt at growth rate one, a sparse class’s rate is pooled over species, and the weights, arrival rate and capacity are held fixed.

The forecast’s rank correlations, 0.712 and 0.683 one and two censuses ahead for the model, change little with the stage structure (Table S21). Only the parametric continuum lowers them by more than 0.05 at both horizons, to 0.628 and 0.622. The two structures without an edge at 79.4 mm lower the rank correlation by slightly more than 0.05 one census ahead only. Under every structure the forecast ranks the changes better than an extrapolation of each species’ long-term trend (Section S3).

The expected stationary composition is the regional composition $p_s$ whatever the structure (Section S4), so the rank correlation stays at 0.73. The identity-line $R^2$, +0.494 for the model, lies between +0.48 and +0.54 for every structure except the parametric continuum. There the stand collapses to 289 large trees, because two species whose gain $G_s$ (Section S4) is near zero under those rates receive 87% of the arrivals, which the trade-off scheme weights by $1/G_s$. The gain is near zero because the parametric model fits mean growth as a straight line in log diameter. For these two understory species, whose stems are mostly small, the line predicts that the average stem stops growing, and then shrinks, at a diameter well below 79.4 mm, although both species have large trees in the census. With nonparametric rates on the same axis the composition fits as well as with the published classes ($R^2 = +0.498$). The ordering of the gain across species is nearly the same under every structure (rank correlation at least 0.96 with the ordering under the classes of the main text).

We also compared the eight size classes with finer descriptions of size on twelve species: the five most abundant, the six with the fewest stems, and *Gustavia superba*, which holds 53% of the model’s seedlings in the simulations of the main text. We fitted each description to the censuses of 1982–1995 and scored it on the later censuses, from the starts of 1995, 2000 and 2005. With eight classes the rank correlation of predicted with observed change in large trees is 0.92 one census ahead and 0.97 two ahead. With sixteen classes and with two descriptions of growth and loss on a continuous diameter axis it is 0.83 to 0.94 one census ahead and 0.87 to 0.98 two ahead.

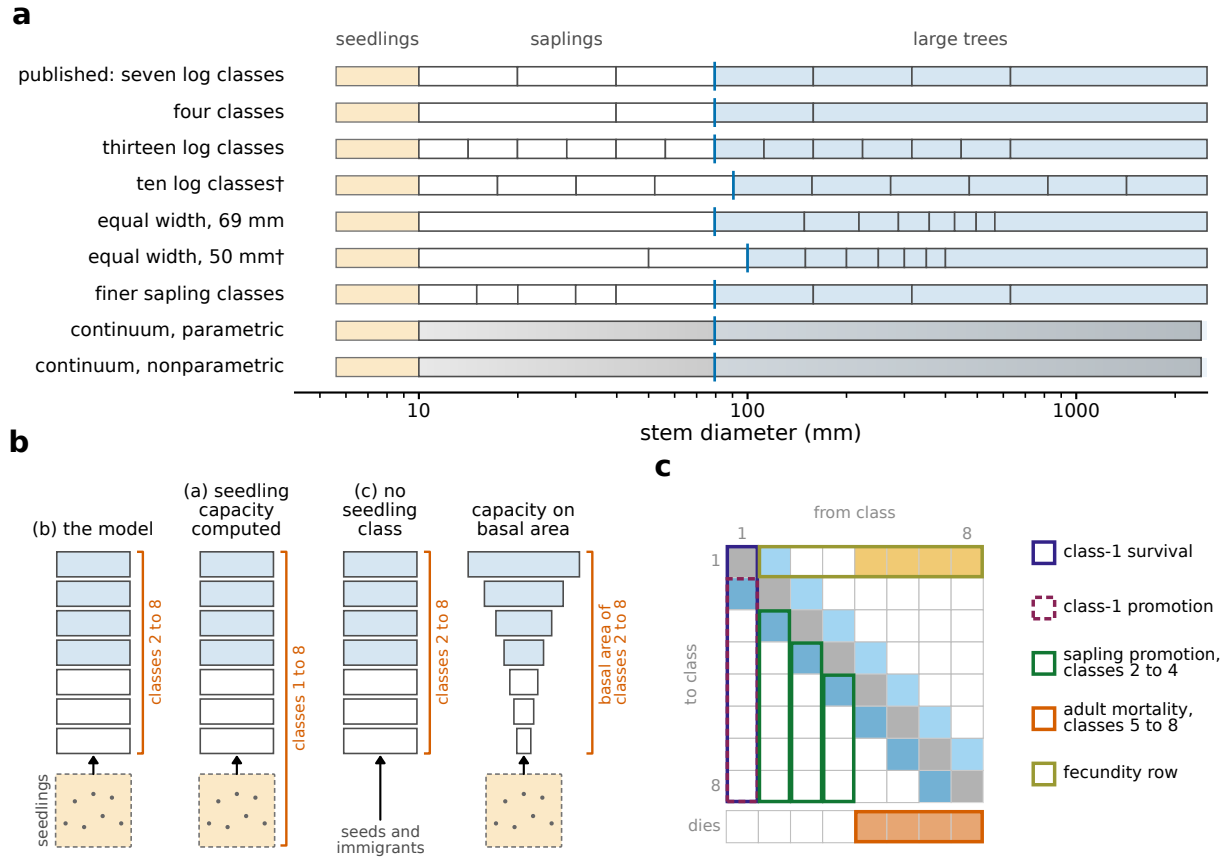

Figure S5: **The variants of the model.** Each variant changes one element. (a) The diameter classes of the published matrices and of the eight alternative stage structures, on a logarithmic diameter axis. Ticks are class edges, the seedling pool below 10 mm is at the left, and the large-tree classes are shaded from 79.4 mm, or from the nearest edge where no edge lies there. The continuum models have a continuous axis. (b) The classes that the capacity holds: classes 2 to 8 with the seedling pool free (the model), all classes with a computed seedling component, classes 2 to 8 with no seedling class, and basal area, with the classes drawn in proportion to their weights. (c) The five groups of rates changed by 20% in the sensitivity test, shown as blocks of the matrix.

The census enumerates only stems of at least 10 mm dbh, which leaves open how the seedling class should be limited. We compared three treatments of the seedling class, lettered as in Table S21. In the model, treatment (b), the capacity applies to the censused stand only and births enter class 1 freely, so the size of the seedling class, fed by the standing trees and limited by seedling mortality, is a prediction. Treatment (a) places one capacity on all eight classes, with a seedling component of 1,786,436 stems. This component is the number of seedlings the standing trees would produce, obtained by applying the fecundity rows of our matrices to the first-census counts in classes 2 to 8. We did not take this component from the seedling census, which enters the model only through the seedling column of the matrices (Section S2) and whose counts of standing seedlings serve only as a test (Section S4). Treatment (c) omits the seedling class, and recruits enter class 2 directly.

We solved the three treatments in closed form, with the gains recomputed for each. Relative to the model, treatment (a) lowers the large-tree total from 1.23 to 1.09 times the observed and treatment (c) lowers it to 1.22. The rank correlation stays at 0.73, and $R^2$ goes from +0.494 to +0.478 and +0.493. The seedling pool holds 1.81 million stems under (a) and 2.09 million under (b), and the flow into class 2 is 28,261 and 31,873 stems per interval, compared with 14,900 to 35,300 in the censuses. Under (a) the assumed seedling capacity sets the large-tree total and little else. As the capacity goes from a quarter to four times the computed value, the total runs from 0.36 to 4.04 times the observed, scaling with the capacity at log-log slope 0.880, while the rank correlation is unchanged (range 0.00). Omitting the seedling class changes the forecast large-tree total two censuses ahead by at most 0.12% of the observed total (Section S3).

In the model the capacity is a number of censused stems, whatever their size. We also placed the capacity on basal area. Each class $k$ is then weighted by the square of its midpoint diameter, $10^{0.6(k-2)}$ times the class-2 value and 3,981 in the top class (closed at 1,259 mm). The capacity is the sum of the observed class totals so weighted, and recruits fill the unused basal area, each counted at the basal area of a class-2 stem. With the stem capacity the stationary stand carries 1.27 times the observed basal area. With the capacity at the observed basal area and the trade-off weights recomputed, the stand holds 0.79 of the observed censused stems and 0.97 times the observed large trees (1.23 in the model). The composition is unchanged (rank correlation 0.73, $R^2 = 0.456$ compared with +0.494). The seedling pool falls from 2.09 to 1.66 million, which is still 1.2 to 1.8 times the seedling census (Section S4).

The gains can also be equalized through each species’ fecundity, instead of through the weights $1/G_s$ on arrivals. The expected composition is then the same (main text, Long-run composition), but the fecundities required are not feasible. The nearest feasible adjustment gives a rank correlation of 0.725 and $R^2 = +0.469$, compared with 0.727 and $R^2 = +0.494$ for the model.

To see how an error in the rates affects the results, we changed the rates by 20% up and down and recomputed the forecast and the closed form. The rates were changed in five groups (the row groups of Table S21 and the fecundity row), one group at a time and in every species at once. Each affected fecundity row was then rescaled to growth rate one, which cancels a uniform change of that row. A 20% error in sapling promotion or adult mortality moves the forecast’s rank correlation by at most 0.06 and leaves its margin over the long-term trend of each species at least +0.11. With the gains recomputed, the stationary composition is the regional one in every scenario, and the rank correlation and $R^2$ move by at most 0.003 and 0.07. The same error in one species alone moves the forecast by at most 0.001. If arrivals are instead weighted with the unperturbed gains, an error in one species moves the rank correlation and $R^2$ by at most 0.005 and 0.08, and an error common to all species lowers $R^2$ to +0.15 only when every species' seedling survival lies above its estimate at once. Without the rescaling a species whose growth rate rises above one takes over the stand (15 of 30 such scenarios). With every growth rate held at one, the uncertain seedling and fecundity entries have little effect on the forecast or on the stationary abundances.

| variant | forecast $\rho$ | | stationary, trade-off scheme | | |
| --- | --- | --- | --- | --- | --- |
| | 1 ahead | 2 ahead | $\rho$ | $R^2$ | large trees<br>(× obs.) |
| *stage structure* |  |  |  |  |  |
| published, seven log classes (the model) | 0.712 | 0.683 | 0.73 | +0.494 | 1.23 |
| four classes | 0.711 | 0.674 | 0.73 | +0.499 | 1.31 |
| thirteen log classes | 0.708 | 0.684 | 0.73 | +0.485 | 1.15 |
| ten log classes<sup>†</sup> | 0.656 | 0.654 | 0.73 | +0.539 | 0.97 |
| equal width, 69 mm | 0.674 | 0.652 | 0.73 | +0.497 | 1.28 |
| equal width, 50 mm<sup>†</sup> | 0.660 | 0.648 | 0.73 | +0.510 | 0.71 |
| finer sapling classes | 0.713 | 0.683 | 0.73 | +0.484 | 1.14 |
| continuum, parametric rates | 0.628 | 0.622 | 0.73 | -3.698 | 0.01 |
| continuum, nonparametric rates | 0.725 | 0.684 | 0.73 | +0.498 | 1.29 |
| *seedling class and capacity* |  |  |  |  |  |
| (a) seedling capacity computed | — | — | 0.73 | +0.478 | 1.09 |
| (b) seedling pool uncapped, the model | — | — | 0.73 | +0.494 | 1.23 |
| (c) no seedling class | — | 0.86<sup>‡</sup> | 0.73 | +0.493 | 1.22 |
| capacity on basal area | — | — | 0.73 | 0.456 | 0.97 |
| *rates changed by 20% for every species at once, fecundity rescaled* |  |  |  |  |  |
| class-1 survival, down | — | — | 0.73 | +0.494 | 1.23 |
| class-1 survival, up | — | — | 0.73 | +0.494 | 1.23 |
| class-1 promotion, down | — | — | 0.73 | +0.494 | 1.23 |
| class-1 promotion, up | — | — | 0.73 | +0.494 | 1.23 |
| sapling promotion, classes 2–4, down | 0.725 | 0.687 | 0.73 | +0.425 | 0.86 |
| sapling promotion, classes 2–4, up | 0.662 | 0.642 | 0.73 | +0.500 | 1.61 |
| adult mortality, classes 5–8, down | 0.657 | 0.640 | 0.73 | +0.502 | 1.44 |
| adult mortality, classes 5–8, up | 0.724 | 0.685 | 0.73 | +0.476 | 1.08 |

Table S21: **Forecast and stationary scores of the model variants.** Each row is the model of the main text with one element changed: the stage structure rebuilt from the census stem measurements, the placement of the capacity, or one group of the rates changed by 20% in every species at once, with each species’ fecundity rescaled to keep its growth rate at one. The first row and row (b) are the model. The forecast columns give the Spearman rank correlation of predicted with observed change in large-tree abundance one and two censuses ahead, over the 252 species-start pairs of the out-of-sample forecast with the rates inferred from the censuses of 1982–1995. The forecast uses the rate matrices alone, which the capacity does not enter, so rows (a), (b) and the basal-area capacity share the forecast of the first row, and the class-1 rate rows do not reach it. The stationary columns give the Spearman rank correlation and the identity-line $R^2$ on log abundance of the stationary large trees (stems of at least 79.4 mm dbh) against the census mean over the 84 species, and the stationary large-tree total relative to the observed 21,693. They are from the closed form for the trade-off scheme at 26,499 immigrant seeds per interval, with the nine species absent from the regional plots entered at the smallest regional share. The expected composition is the regional composition for every variant, so the rank correlation is the same in every row to two decimals and only the total and $R^2$ change. Its third decimal depends on the order given to species with identical predicted abundance. The text explains the low values on the parametric continuum. <sup>†</sup>No class edge at 79.4 mm, so the large-tree threshold moves to the nearest edge and the large-tree population differs from that of the other rows. <sup>‡</sup>In sample (every start census), two censuses ahead, where the seedling class enters the forecast through the stems it promotes. One census ahead it does not enter. The in-sample value of the model is 0.86.

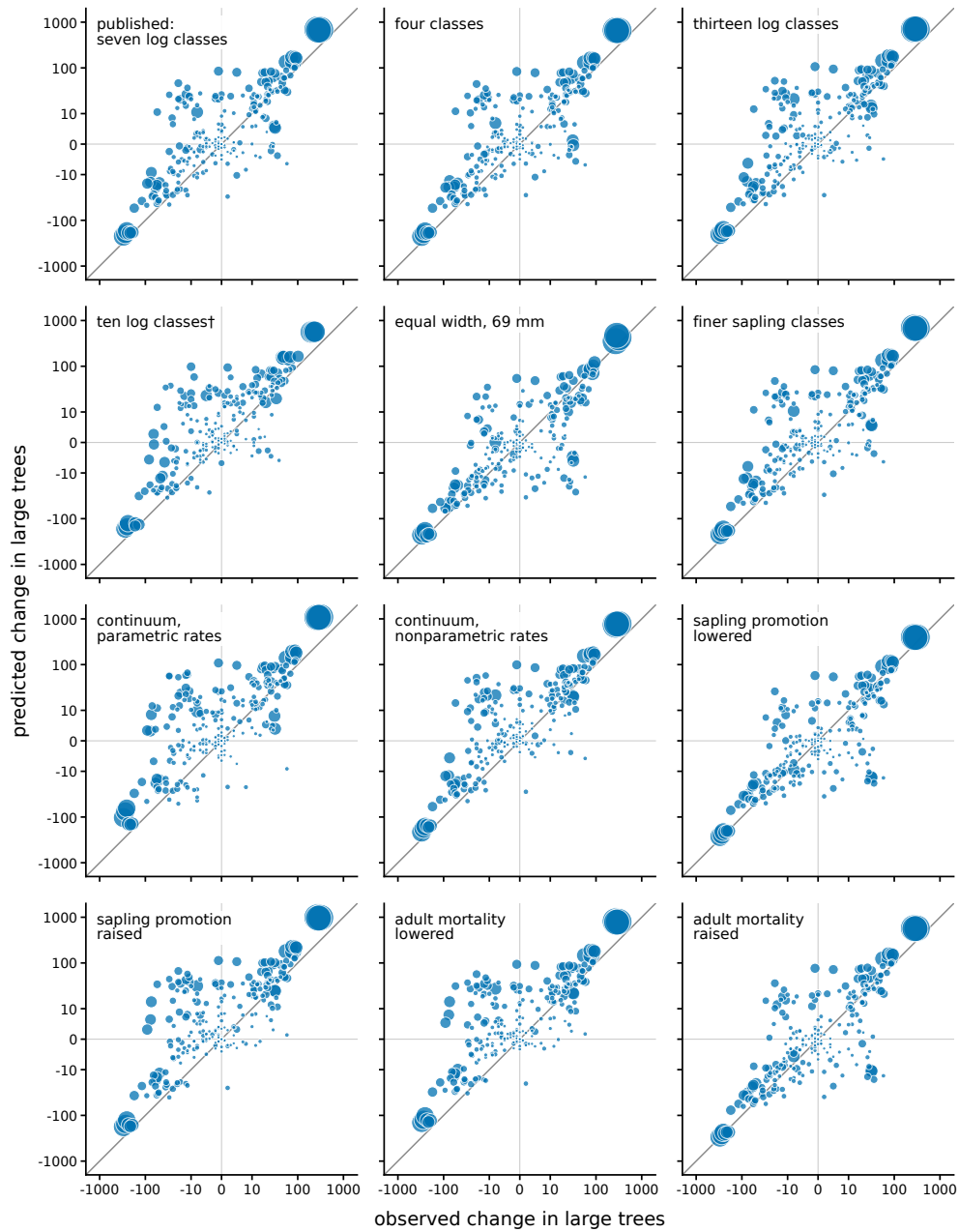

Figure S6: **Forecast of change in large trees two censuses ahead under the model variants.** Predicted against observed change in each species' large trees two censuses ahead, out of sample (rates from 1982–1995 applied at 1995, 2000 and 2005, 252 pairs per panel). The panels are the published stage structure, seven alternative structures, and sapling promotion and adult mortality changed by 20% in every species. Symbol area scales with the species' large trees at the start, the line is the identity, and the axes are logarithmic away from zero. The rank correlation of each panel is its variant's forecast entry in Table S21. It is close to that of the model in every panel and lowest on the parametric continuum. <sup>†</sup>No class edge at 79.4 mm, so large trees are counted from 90.6 mm.

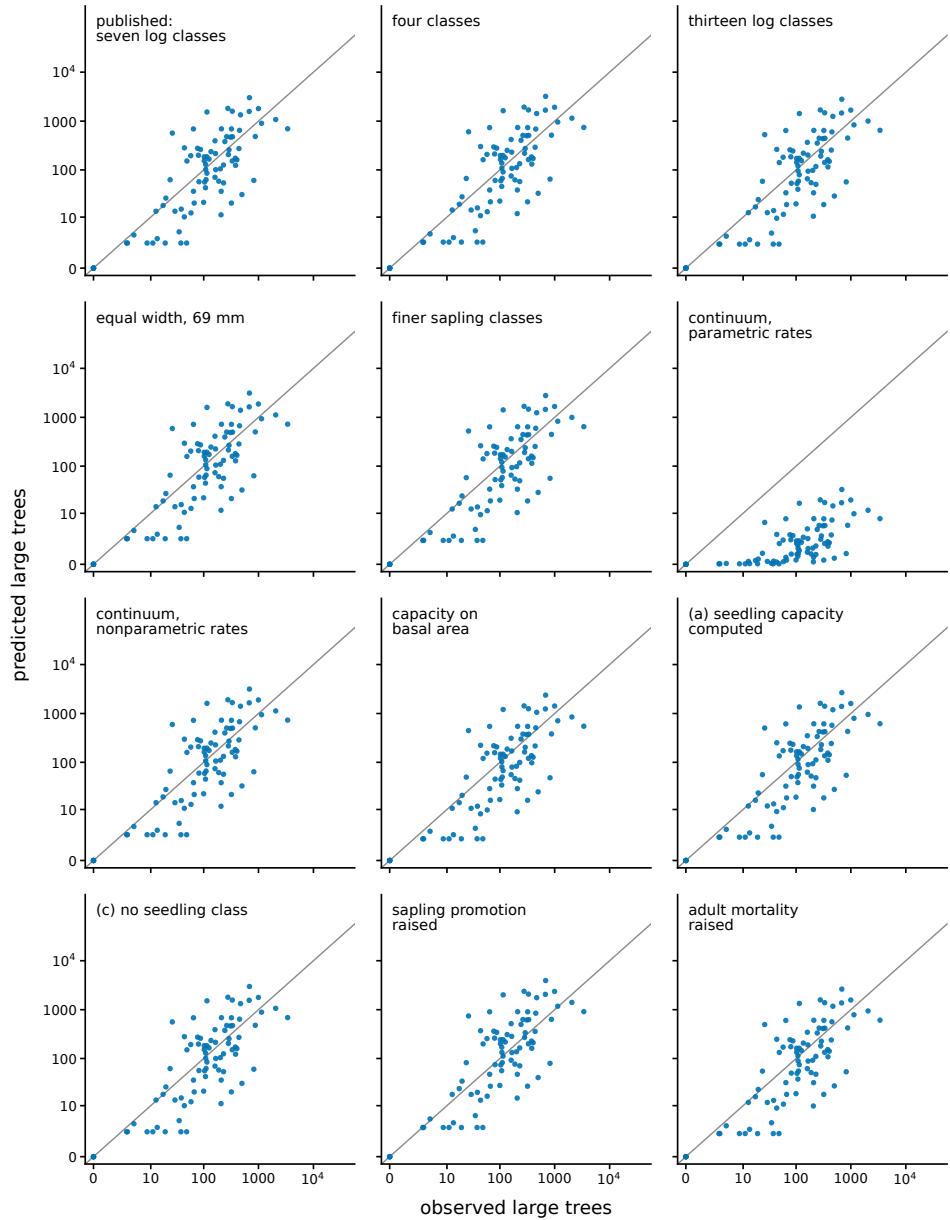

Figure S7: **Stationary large-tree abundances under the model variants.** Stationary large-tree abundance of each species under the trade-off scheme against its census mean (log axes, identity line), for the published structure, six alternative structures, the capacity on basal area, the two other treatments of the seedling class, and sapling promotion and adult mortality raised by 20%. The rank correlation and $R^2$ of each panel are its variant's entries in Table S21. The abundances fit the census about as well under every variant as under the model, except on the parametric continuum. There two understory species receive 87% of the arrivals and the stand holds 289 large trees in all (see text). With nonparametric rates on the same continuous axis the panel matches that of the published structure.

### Adding further species to the model

Jops et al. (2025) published matrices for the 90 species of the plot that have at least 200 stems in every census and enough seedling records, a threshold they set so that the rates could be estimated with confidence. The tree censuses of 1982–2015 record 328 species codes with a live stem of at least 10 mm dbh at some census. They are the 90 published species, seven codes not identified to species and 231 further species, which we call the candidates. For each candidate we built a matrix by the procedure of Section S2. Columns 2 to 8 are estimated from the tree censuses by the method of Jops et al. (2025). The seedling column is inferred from the seedling census by the rules described in Section S2, and the fecundity row is scaled to growth rate one. Applied to the 90 published species with their published reproductive classes, the same procedure returns the matrices used in this paper. The procedure gives a matrix for 76 of the candidates. Most of the other 155 lack a promoted seedling: 77 have seedlings in the seedling census of which none was promoted, and 67 have no seedling in that census. The remaining species have a gap in the size classes between the seedlings and the first reproductive class (four species), no tree with a recorded position (three), no occupied reproductive class (one), a growth rate of one or more from survival and growth alone (two), or a seedling column that sums to one or more (one). The species without a matrix held 3,647 of the plot's 28,348 large trees at the census of 2015. Palms account for 1,964 of these trees, and their eleven species have no seedling in the seedling census.

We obtained error bars by resampling the individuals of each species. For each of the 84 modeled species and of the 76 candidates with a matrix we drew its trees and its seedlings with replacement, each individual with all of its records, and rebuilt the matrix, 1,000 times. We measure the uncertainty of a quantity by its width, the ratio of its 95th to its 5th percentile over the replicates. The stationary abundances depend on the matrix through the gain $G_s$ (Section S4), the number of large trees that a species holds per arriving seed. For the modeled species the width of the gain has a median of 3.2 and runs from 1.3 to 20 over the 76 species for which it is finite. It is infinite for five species, for which at least one replicate in twenty has no large tree, and three species have no large tree in at least 95% of the replicates. For the candidates the median width is 10, with a range of 2.5 to 61 over 54 species. Another four widths are infinite, and 18 species have no large tree in at least 95% of the replicates. The rates of the candidates are thus less certain than those of the modeled species, as Jops et al. (2025) expected when they set their threshold. The gain depends on the promotion rate out of the seedling class, and promotions are few. The median candidate has five promoted seedlings and the median modeled species has 33. For the 36 candidates with fewer than five (and for six of the modeled species) the width understates the uncertainty, because a replicate with no promoted seedling gives no matrix and is drawn again.

We take the error bars of a candidate to lie within those of the modeled species when four quantities are each no larger than the 90th percentile of the same quantity among the 84 modeled species. The four are the width of its gain, the widths of its seedlings per arriving seed and of its fecundity, and the share of its replicates that give no matrix. The condition is met by sixteen candidates. Among them are six shrubs and treelets whose matrices stop below the large-tree classes, like the six published species that the model leaves out (Section S2), and one species that meets the condition only if the five infinite widths are counted in the percentile. The other nine are the species we add to the model: *Cecropia insignis*, *Jacaranda copaia*, *Zanthoxylum ekmanii*, *Cordia alliodora*, *Spondias radlkoferi*, *Inga thibaudiana*, *Guarea bullata*, *Ocotea oblonga* and *Hampea appendiculata* (Table S22). The same nine species meet the condition under a second set of bootstrap replicates, although four of the other candidates then cross the 90th percentile. The widths of their gains run from 2.5 to 7.8. They held 1,277 large trees in 2015, and adding them would raise the share of the plot’s large trees that the model covers from 77.1% to 81.6%. A weaker condition, every width no larger than the widest among the modeled species, is met by 64 candidates, the sixteen among them, and would bring the share to 86.4%. It sets no bound on the gain, because the widest gain among the modeled species is infinite.

| species | records behind the matrix | | | | width<br>of $G_s$<br>(ratio) | regional<br>plots<br>(of 28) | large trees | | |
| --- | --- | --- | --- | --- | --- | --- | --- | --- | --- |
|  | trees | seedlings | seedlings<br>promoted | handed<br>over |  |  | census,<br>2015 | census<br>mean | model |
| *Cecropia insignis* | 3,138 | 421 | 21 | 21 | 2.5 | 17 | 523 | 328 | 249 |
| *Jacaranda copaia* | 547 | 61 | 5 | 5 | 5.4 | 20 | 229 | 238 | 1,416 |
| *Zanthoxylum ekmanii* | 740 | 79 | 4 | 4 | 7.8 | 7 | 145 | 141 | 77 |
| *Cordia alliodora* | 409 | 260 | 7 | 5 | 6.2 | 10 | 80 | 64 | 41 |
| *Spondias radlkoferi* | 701 | 194 | 16 | 7 | 4.1 | 17 | 74 | 72 | 256 |
| *Inga thibaudiana* | 442 | 169 | 20 | 11 | 4.4 | 7 | 68 | 33 | 32 |
| *Guarea bullata* | 2,192 | 88 | 20 | 12 | 4.6 | 4 | 59 | 106 | 39 |
| *Ocotea oblonga* | 748 | 316 | 30 | 20 | 3.4 | 6 | 57 | 42 | 40 |
| *Hampea appendiculata* | 406 | 123 | 18 | 8 | 3.7 | 3 | 42 | 26 | 21 |
| all nine |  |  |  |  |  |  | 1,277 | 1,049 | 2,171 |

Table S22: **The nine species added to the model.** These are the species beyond the 90 published ones whose error bars lie within those of the 84 modeled species. The first four columns count the trees and the seedlings of the species that were resampled, the seedlings promoted out of class 1, and those among the promoted that the seedling census recorded as handed over to the tree census (Section S2). The width of $G_s$ is the ratio of the 95th to the 5th percentile of the gain over 1,000 resamplings of the individuals, and its median among the modeled species is 3.2. The regional plots counted are those of the 28 plots of Section S2 on which the species has a large tree (a stem of at least 79.4 mm dbh). The census columns give the count of large trees at the census of 2015 and the mean over the censuses of 1982–2010, with which the model is compared. The model column is the stationary number under the trade-off scheme, from the closed form of Section S4 with the environmental noise off and the nine species added to the modeled species. The model places seven of the nine within a factor of three of their census mean and predicts *Jacaranda copaia*, which is far more common in the region than on the plot, at 6.0 times its census mean. Rows are in order of the 2015 count.

For *Cecropia insignis*, *Jacaranda copaia* and *Zanthoxylum ekmanii* every promoted seedling is one that the seedling census recorded as handed over to the tree census (Section S2), and none was promoted on a measured dbh. The promotion rates of these three species therefore depend entirely on counting a seedling that was handed over as promoted. For the first two, no seedling stayed in the seedling class from one seedling census to the next. The fecundity row of a candidate starts at the first reproductive class of the species, but the table of minimum reproductive diameters published with the matrices (Jops et al., 2025) covers the 90 species only. For a candidate we therefore set the first reproductive class to the highest occupied class minus two, and no lower than class 2. On the 90 published species this rule returns the published class for 49, a lower class for 39 and a higher class for two. The choice changes the seed production and with it the arrival rate, on which the censused composition does not depend.

We solved the closed form of Section S4 for the modeled species alone, for the modeled species with the nine species added and for the modeled species with all 64 added. All model values in this subsection and its two tables come from this closed form, for the trade-off scheme with the environmental noise off. We did not run the stochastic simulations with added species, and the seed production of the larger sets is the mean-field value with the correction found for the 84 modeled species. The capacity and the arrival rate of each set are derived as for the modeled species. With the nine species added the capacity is 203,808 censused stems instead of 200,721, and the arrival rate is 27,175 seeds per interval instead of 26,499. The regional shares are normalized over the species of the set, which multiplies the shares of the modeled species by 0.923 with the nine species added and by 0.771 with all 64 added. All nine added species occur on the regional plots. Among the 64 species, 18 are absent and enter at the smallest share of any sampled species (Section S2).

Table S23 gives the scores with their 90% ranges from two sources of uncertainty, the rates (the replicates described above) and the regional shares (the 28 regional plots resampled 1,000 times). Each range varies one source with the other held fixed. Adding the nine species lowers the large-tree total of the modeled species slightly, from 1.232 to 1.206 times the observed. The range of this total for the modeled species alone is 1.149 to 1.287 from the rates and 1.089 to 1.399 from the regional plots. $R^2$ goes from +0.494 to +0.492, and the rank correlation stays at 0.73. Every score of the modeled species lies inside its range for those species alone, from both sources. This holds also with the capacity and the arrival rate held at their values for the modeled species, where the large-tree total is 1.188 times the observed.

The fall in the large-tree total is small but systematic. At the estimated rates and regional shares the total falls by 0.025. Replicate by replicate it falls by 0.019 to 0.033 over the rates and by 0.008 to 0.044 over the regional plots. The fall arises because the modeled species give up regional share to the added species, and the larger capacity offsets part of the loss. With all 64 species added the total of the modeled species falls to 1.082 times the observed, below both ranges.

As a group the nine added species are predicted less well than the modeled species, and *Jacaranda copaia* accounts for most of the difference. Together they hold 2.07 times their observed large trees, compared with 1.206 for the modeled species in the same solution, and their $R^2$ is $-0.069$ compared with $+0.492$. *Jacaranda copaia* is modeled at 1,416 large trees, and its census mean is 238. Under the trade-off scheme the expected large trees of a species are proportional to its regional share (Section S4), and *Jacaranda copaia* is far more common in the region than on the plot. It has 321 large trees on 20 of the 28 regional plots, 5.0% of the regional total of the 93 species. Its census mean lies below the model's range from the rates (1,322 to 1,478) and from the regional plots (344 to 2,573), so sampling error in the rates or in the regional plots cannot explain the difference. The other eight species hold 0.93 times their observed large trees, with $R^2 = +0.325$.

| species in<br>the model | species<br>scored | | seedlings<br>(× census) | large trees<br>(× obs.) | $R^2$ of log<br>abundance | rank<br>correlation $\rho$ |
| --- | --- | --- | --- | --- | --- | --- |
| the 84 | the 84 | value | 1.87 | 1.232 | +0.494 | 0.73 |
|  |  | range, rates | 1.56 to 4.26 | 1.149 to 1.287 | +0.471 to +0.495 | 0.72 to 0.73 |
|  |  | range, regional plots | 1.41 to 2.47 | 1.089 to 1.399 | +0.185 to +0.509 | 0.66 to 0.75 |
| the 84 and the nine | the 84 | value | 1.83 | 1.206 | +0.492 | 0.73 |
|  |  | range, rates | 1.52 to 4.17 | 1.126 to 1.260 | +0.469 to +0.493 | 0.72 to 0.73 |
|  |  | range, regional plots | 1.38 to 2.43 | 1.062 to 1.374 | +0.180 to +0.508 | 0.66 to 0.75 |
|  | all 93 | value | 1.86 | 1.246 | +0.479 | 0.73 |
|  |  | range, rates | 1.55 to 4.19 | 1.163 to 1.301 | +0.456 to +0.480 | 0.72 to 0.74 |
|  |  | range, regional plots | 1.41 to 2.45 | 1.132 to 1.378 | +0.146 to +0.498 | 0.66 to 0.75 |
| the 84 and the 64 | the 84 | value | 1.65 | 1.082 | +0.477 | 0.73 |
|  |  | range, rates | 1.35 to 3.71 | 1.003 to 1.114 | +0.451 to +0.478 | 0.72 to 0.73 |
|  |  | range, regional plots | 1.24 to 2.20 | 0.955 to 1.244 | +0.142 to +0.490 | 0.66 to 0.75 |
|  | all 148<sup>†</sup> | value | 1.91 | 1.260 | +0.608 | 0.73 |
|  |  | range, rates | 1.70 to 4.04 | 1.168 to 1.297 | +0.597 to +0.610 | 0.73 to 0.74 |
|  |  | range, regional plots | 1.49 to 2.48 | 1.162 to 1.388 | +0.427 to +0.625 | 0.68 to 0.75 |

Table S23: **Scores of the model with species added.** All values are for the trade-off scheme, from the closed form of Section S4 with the environmental noise off, and the capacity and the arrival rate are derived for each set of species as for the 84 modeled species. Seedlings and large trees are the stationary totals of the species scored, relative to their seedling-census mean and to their census mean of large trees. $R^2$ is the identity-line $R^2$ on log abundance of large trees, and the rank correlation is the Spearman correlation of stationary with census-mean large-tree abundance. Each range runs from the 5th to the 95th percentile over 1,000 resamplings of one source of uncertainty with the other held fixed, and its ends are rounded outward. The sources are the rates, with the individuals of every species resampled, and the regional shares, with the 28 regional plots resampled. With the nine species of Table S22 added, every score of the modeled species stays inside its range for the modeled species alone. With 64 species added, the large-tree total of the modeled species falls below that range. <sup>†</sup>Of the 64 added species, 17 have no large tree in the model or in the census, and their agreement at zero raises $R^2$. On the other 131 species $R^2$ is +0.297.

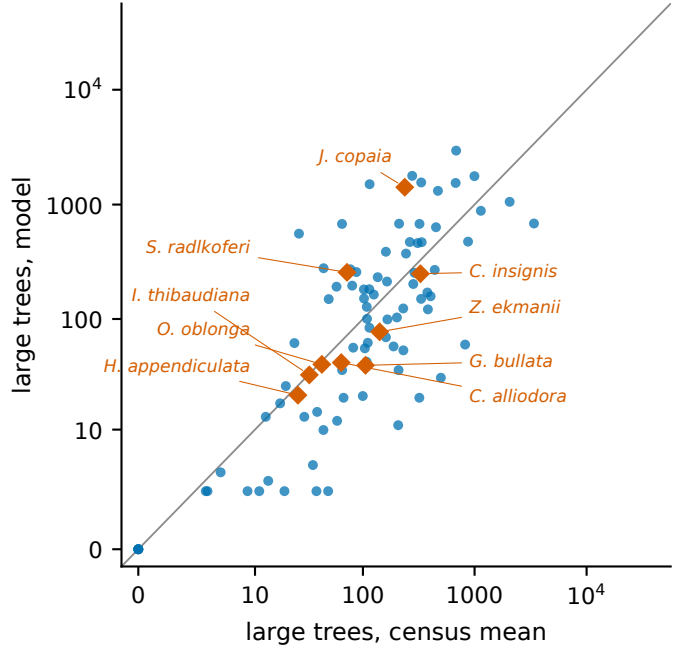

Figure S8: **Stationary large trees of the modeled and the added species against the census.** Stationary large trees of each species under the trade-off scheme against its census mean (log axes away from zero, identity line), from the closed form with the environmental noise off, for the 84 modeled species (circles) and the nine added species (diamonds, named). Over the 93 species the rank correlation is 0.73 and $R^2 = +0.479$, compared with 0.73 and $R^2 = +0.494$ for the modeled species alone. The largest departure is *Jacaranda copaia*, modeled at 1,416 large trees with a census mean of 238.

Species by species, the added species are no further from the census than the modeled species. With one added to every abundance, so that species with no large tree are counted, seven of the nine lie within a factor of three of the census mean, as do 54 of the 84 modeled species. The root-mean-square difference in $\log_{10}$ of these abundances is 0.37, compared with 0.54 for the modeled species. The $R^2$ of the added species is nevertheless low, because their census abundances span a narrow range. The census mean lies inside the model's range from the regional plots for six of the nine and inside its narrower range from the rates for one. The 64 species that meet the weaker condition are over-predicted, by a factor 2.8 in large trees and 7.5 in seedlings, and 26 of the 47 whose matrices reach the large-tree classes lie within a factor of three. Figure S8 shows the 93 species against the census.

Adding the nine species leaves the results for the modeled species unchanged within their ranges. The added matrices, however, depend on counting seedlings handed over to the tree census as promoted and on our rule for the first reproductive class. We therefore keep the 84 modeled species in the main text and in the rest of this supplement.

### Trees that fall below the 10 mm census threshold

A tree with a stem of at least 10 mm dbh at one census can be recorded at the next as alive with no dbh. We call such a record a fall-back record and the tree a fall-back tree, and we count tree-intervals, each one tree over one census interval. Over the seven census intervals of 1982–2015 the 84 modeled species have 46,204 fall-back records, from 44,933 trees, and 98.8% of the records carry a resprout code. The records are 3.3% of the 1,404,973 tree-intervals that start with a stem of at least 10 mm. In *Gustavia superba* they are 51.6% of the tree-intervals that start in class 2, 28.6% of those that start in class 3 and 2.4% of those that start in class 4 (Table S24). Jops et al. (2025) count these trees as lost, and so do our matrices (Section S2).

We followed the 35,144 fall-back records of 1990 to 2010 to the next census. By then 39.2% were dead, 43.5% were still alive with no stem of 10 mm and 17.3% had such a stem again. For comparison, seedlings in our matrices die at 37.0% per five years and class-2 trees at the census of the fall-back die at 11.2%, each rate being that of the record’s species averaged over the records followed. Fall-back trees thus die about as often as seedlings, and they return to 10 mm 5.9 times as fast as seedlings are promoted. Among the 100 records of *Gustavia superba*, 31 were dead, 62 were alive with no stem of 10 mm and 7 had such a stem again, and the seedlings of this species die at 48.2% per five years in our matrices. We left out the 5,087 records of 1985 because the census of 1990 recorded 2,294 of them as not found and none as dead, and the records of 2015 have no later census. With the records of 1985 included and those not found counted as dead, 40% of all records and 36% of the 138 records of *Gustavia superba* are dead at the next census.

To see whether the treatment of these trees matters, we built alternative matrices in which they enter the seedling class. In each column of classes 2 to 8 of $U_s$, the share of the trees of the class that fall back is added to row 1. The seedling column is that of our matrices, and the fecundity row is rescaled so that the growth rate is one. The constants that follow from the fecundity rows change with it. The seed production $B$ goes from 1.01 to 0.99 million seeds per interval, the arrival rate $M = \hat{m}B$ from 26,499 to 26,072 seeds per interval, and the seedling component of the capacity in treatment (a) of the seedling class, described earlier in Section S6, from 1,786,436 to 1,774,091 stems.

The predictions of the model are nearly the same with the two sets of matrices (Table S25, trade-off scheme). In the closed form of Section S4, without environmental noise, every abundance agrees at the precision of the table, and the e-folding time goes from 19,000 to 19,100 years. In the simulations with environmental noise (72 runs per set), the large-tree total stays at 0.84 times the observed and the seedling pool at 1.7 times the census mean, and the saplings go from 0.65 to 0.66 times the census mean. The simulated rank correlation and $R^2$ in the table are computed on the pooled runs, and measured against the spread of the pooled value over resampled runs their differences between the two sets are $-1.1$ and $-0.4$. The decade-ahead forecast with the rates inferred from the censuses before each start (Section S3) is unchanged.

The seedling census does not count tagged trees, so the comparison of seedlings can also be made with the fall-back trees added to the census. The tagged trees that are alive with no stem of 10 mm number 10,716 on average, 0.96% of the seedling-census mean. With these trees added to that mean, the seedlings of the alternative set are 1.85 times the census in the closed form and 1.7 in the simulations.

With the alternative matrices we also repeated the simulations with the noise off (36 runs), the simulations behind the five-year relative change of the large-tree total given in Section S7 (36 runs), the in-sample forecast, in which the matrices are applied at the first five censuses (Section S3), and the trait comparison (Table S19). In the noise-off simulations 6 of the 8 quantities we compared differ between the two sets in the digits we report. In the simulations behind the relative change 5 of 7 do, and the relative change of the large-tree total itself stays at 0.020. In the in-sample forecast only stem totals and the differences between the model with and without its seedling class change, and in the trait comparison the values that change do so in the second decimal. We did not recompute the other results with the alternative matrices.

|  | 84 modeled species | *Gustavia superba* |
| --- | --- | --- |
| *fall-back records, % of the tree-intervals that start in the class* |  |  |
| class 2 | 6.1 (33,993 of 560,438) | 51.6 (99 of 192) |
| class 3 | 2.2 (10,206 of 466,165) | 28.6 (36 of 126) |
| class 4 | 0.77 (1,740 of 226,485) | 2.4 (9 of 375) |
| classes 5 to 8 | 0.17 (265 of 151,885) | 0.17 (8 of 4,792) |
| all classes | 3.3 (46,204 of 1,404,973) | 2.8 (152 of 5,485) |
| *state at the next census, % of the fall-back records followed* |  |  |
| fall-back records followed | 35,144 | 100 |
| dead | 39.2 (13,759) | 31.0 (31) |
| alive with no stem of 10 mm | 43.5 (15,294) | 62.0 (62) |
| with a stem of at least 10 mm | 17.3 (6,089) | 7.0 (7) |
| *rates for comparison, % per five years* |  |  |
| class-2 trees, dead | 11.2 | 13.4 |
| class-2 trees, alive with no stem of 10 mm | 8.4 | 46.8 |
| seedling class of our matrices, dead | 37.0 | 48.2 |
| seedling class of our matrices, promoted | 3.0 | 0.36 |
| *ratios* |  |  |
| death, fall-back trees over class-2 trees | 3.5 | 2.3 |
| death, fall-back trees over the seedling class | 1.06 | 0.64 |
| return to 10 mm over promotion of seedlings | 5.9 | 19.5 |

Table S24: **Frequency and fate of trees that fall below 10 mm.** A fall-back record is a tree with a stem of at least 10 mm dbh at one census that is recorded as alive with no dbh at the next, and such a tree is a fall-back tree. A tree-interval is one tree over one census interval, and a tree can fall back more than once. First block: the seven intervals of 1982–2015, by the class of the tree at the start of the interval, with the counts in parentheses. Another 1,016 tree-intervals end with status AD, AM or AR, and both sets of matrices count them as lost. Second block: the fall-back records of 1990 to 2010 at the census after the fall-back, with the counts in parentheses. Among the 35,144 records, 2 were not found. The 5,087 fall-back records of 1985 and the 5,973 of 2015 are left out, for the reasons given in the text. With those of 1985 included, 40,231 records are followed. The records alive with no stem of 10 mm include 57 with status AD, AM or AR (1 for *Gustavia superba*). Third block: class-2 trees are the trees of class 2 at the census of the fall-back (377,567 tree-intervals, 123 of *Gustavia superba*), and the seedling class is the seedling column of our matrices (Section S2). In the column of the 84 species each rate is that of the record’s species, averaged over the fall-back records followed. The class-2 rate of trees alive with no stem of 10 mm counts status AD, AM and AR as the second block does. With status A alone it is 8.3% (46.2% for *Gustavia superba*).

|  | our<br>matrices | alternative<br>matrices | difference<br>(in s.e.) |
| --- | --- | --- | --- |
| *closed form, environmental noise off* |  |  |  |
| rank correlation $\rho$ | 0.728 | 0.728 | |
| $R^2$ of log abundance | +0.494 | +0.494 | |
| large trees (× obs.) | 1.23 | 1.23 |  |
| saplings (× census) | 0.97 | 0.97 |  |
| seedlings (× census) | 1.87 | 1.87 |  |
| seedlings without *Gustavia superba* (× census) | 1.17 | 1.17 |  |
| waiting fraction $d$ | 0.025 | 0.025 | |
| e-folding time (years) | 19,000 | 19,100 |  |
| *simulation, environmental noise on, 72 runs per set* |  |  |  |
| rank correlation $\rho$ | 0.714 | 0.710 | -0.7 |
| $R^2$ of log abundance | +0.362 | +0.358 | -0.1 |
| large trees (× obs.) | 0.84 | 0.84 | +1.6 |
| saplings (× census) | 0.65 | 0.66 | +7.2 |
| seedlings (× census) | 1.7 | 1.7 | -0.2 |
| seedlings without *Gustavia superba* (× census) | 0.82 | 0.82 | -0.1 |
| *decade forecast, 252 pairs (independent of the immigration scheme)* |  |  |  |
| rank correlation, five years ahead | 0.72 | 0.72 |  |
| rank correlation, ten years ahead | 0.70 | 0.70 |  |

Table S25: **The model with our matrices and with the alternative matrices.** In the alternative matrices a tree that is alive with no stem of 10 mm enters the seedling class. The closed form (Section S4) and the simulations (Section S7) are for the trade-off scheme. The rank correlation is the Spearman correlation of model with census-mean large-tree abundance over the 84 species, in the closed form with tied abundances given equal rank (Table S12), and $R^2$ is the identity-line $R^2$ on log abundance. Large trees, saplings and seedlings are the model's totals of classes 5 to 8, 2 to 4 and 1 over the census mean, for class 1 the seedling-census mean. The difference is the mean of alternative minus our matrices over runs paired by their random numbers, in units of its standard error. For $\rho$ and $R^2$ the tabulated values are computed on the runs together and the differences on single runs. The forecast rows give the rank correlation of predicted with observed change in large trees over the species-start pairs of the main text, with the rates and the share of trees that fall back inferred from the censuses before each start.

Neither the seedling class nor loss describes the fall-back trees exactly, since they die about as often as seedlings but return to 10 mm several times as fast. Because the predictions are nearly the same with both sets of matrices, we keep the convention of Jops et al. (2025), and the main text and the other sections of this supplement use our matrices. The alternative matrices would not reduce the model’s excess of *Gustavia superba* seedlings (Section S4), because the stable stage structure of this species holds 257 seedlings per large tree under both sets.

### S7 Fluctuation predictions

The form of the environmental noise and its amplitude $\sigma_g$ are estimated from the transition data, and its memory $\tau_g$ is assumed (Section S5). The fluctuations that the noise produces in the number of large trees (stems of at least 79.4 mm dbh) therefore involve no constant fitted to an abundance or to a count of stems. We compare them here with the censuses from which $\sigma_g$ was estimated. The large-tree numbers compared are those of the seven censuses from 1982 to 2010, at which stage vectors are built (Section S2), that is six of the seven intervals behind $\sigma_g$. Later censuses will test them out of sample.

We ran the model to its stationary state under the regional pool and under the trade-off scheme, with 36 replicates of each setting and $\tau_g = 18$ yr throughout. The amplitude was set to its estimate, $\sigma_g = 0.32$, to either end of its 90% confidence interval (0.17 and 0.84) and to zero, which we call the noise off. Each model statistic is a median, with its range, over consecutive non-overlapping windows of seven steps that span the last 2,000 steps of each replicate. We did not vary the memory separately, because the estimated amplitude stays inside that confidence interval over the assumed range of $\tau_g$, 5 to 40 yr.

Table S26 compares the simulated stand with the census. For each statistic we fixed, before the runs, the range of model values that we would accept, and these ranges are the last row of the table. We also required the stationary seedling total to lie within a factor of three of the seedling-census mean (Section S4). To compare individual species we group them in ten bands of initial large-tree abundance on a doubling scale and take, in each band, the mean squared five-year change of a species’ large-tree count. In the two most abundant bands (species of 256 or more large trees) the census value is compared directly with the model’s. In all bands we also ask whether the census value, divided by the model’s value with the noise off, lies inside the range over windows of the model’s ratio of noise on to noise off. For each band we compute a synchronous fraction as well, in the same way for model and census: the part of its species’ squared five-year changes that occurs in proportion to the change of the whole large-tree total.

Without the noise the model is too steady: under the regional pool the median absolute five-year relative change of the large-tree total is 0.0012, against 0.0127 in the census, a factor of 11 smaller (main text, Model section). At the estimated amplitude the regional pool is inside the accepted range on recruitment, on the large-tree total and on the composition. The trade-off scheme is outside it on two statistics. Its range ratio slightly exceeds the limit of 1.10. Its mean large-tree total is 0.66 of its value with the noise off (0.68 in the longer runs described below), where we had accepted a difference of 20%. The regional pool’s total agrees with the census only in its relative fluctuations, since with the noise off its mean is 2.42 times the census mean.

The total falls under the trade-off scheme because too few candidates arrive for the vacancies. In the longer runs with the noise on there are 0.272 candidates for each vacancy opened in a step. The censused classes (2 to 8) then hold 0.669 of the capacity (0.67 and 0.63 from the two sets of initial conditions in the runs of Table S26). With the noise off there are 1.03 candidates per vacancy and the classes hold 0.9998 of the capacity. Under the regional pool they hold 0.956 with the noise on. The capacity is a ceiling on the censused classes, not a fixed occupancy.

For individual species, the census’s mean squared change is within a factor of two of the model’s in both top bands under the regional pool, and in the lower of the two under the trade-off scheme. It lies inside the model’s range in 5 of the ten bands under the regional pool and in 6 under the trade-off scheme, where we had required seven. At stationarity the model falls short at intermediate abundance: for species of 16 to 255 large trees the census’s mean squared change is 2.9 to 3.5 times the model’s under the regional pool, and similar under the trade-off scheme (Fig. S9b; the model started from the 1982 forest is compared below). At the upper end of the amplitude’s confidence interval the census lies inside the model’s range in more bands. The recruitment CV is then three times the measured 0.18, however, and the species of both top bands fluctuate at least twice as much as observed. Under the trade-off scheme the rank correlation also changes by more than five hundredths.

In the top band the synchronous fraction is larger in the model than in the census, because most of the observed variation there is the long-term trend of each species (78% of those species’ summed squared five-year change), which the projection matrices carry (Section S3). After detrending, the abundant species in the census share 29% of their remaining squared change with the total (95% CI 12–57%), against 2% in simulations with the noise off.

Two terms that we measured in the transition data but left out of the model could supply the variability the stationary model lacks at intermediate abundance (Section S5). One is the deviation of each species’ sapling growth from the shared factor, and the other is the year-to-year variation of each species’ mortality. We added the second to the simulations at its measured size, alone and together with the shared growth factor at $\sigma_g = 0.32$ (Table S26, lower rows). It is a lognormal factor of mean one on each species’ five-year mortality hazard, drawn independently for every species at every step, with the standard deviations measured in the transition data for small and for large stems. Alone, the mortality term moves the large-tree total much less than the shared growth factor does: the growth factor moves it 3.4 times as much under the regional pool and 3.6 times as much under the trade-off scheme.

With both terms every species fluctuates more between censuses, and abundant species most. The census’s mean squared change is then within a factor of two of the model’s in both top bands under either scheme. The census lies inside the model’s range in 5 bands under the regional pool and in 6 under the trade-off scheme, still fewer than the seven required, and every other condition is met under both schemes. At intermediate abundance the census’s mean squared change is still 2.6 to 2.9 times the model’s under the regional pool. The species-specific growth deviations ($\sigma_b = 0.075$) are the term that remains to be tested.

Under the trade-off scheme the large-tree total is no longer reduced when the mortality term is added, but this is an effect of the term on mean survival, which a factor of mean one on the hazard raises. We repeated the runs with each species’ baseline hazard rescaled so that its expected five-year survival under the term equals the measured value. The total is then 0.66 of its value with the noise off, and the species-level results are unchanged, with the census inside the model’s range in 5 bands under the regional pool.

The comparisons above set a stationary model against a forest whose species were changing in abundance. We also started the model from the forest of 1982 (Fig. S9). Each of the 84 species starts at its 1982 numbers in classes 2 to 8, with seedlings set as in the forecast of Section S3, and the full model runs six steps, one for each census interval to 2010, in 1,000 realizations. The ratios that follow are ranges over the bands named and over both immigration schemes. With the shared growth factor the census’s mean squared change is 1.09 to 1.49 times the model’s mean in the six bands from 16 trees upward, and 0.33 to 1.02 times in the three bands below, which hold only 6 to 17 species-intervals each. It lies inside the model’s 95% range in all nine bands in which the census records a change, under both schemes. Without environmental noise the ratio from 16 trees upward is 1.49 to 1.92, and the census is above the range in each of those six bands. With mortality noise in place of the growth factor the census is outside the range in two bands, whether the noise is the term above or one factor shared by all species with the memory of the growth factor (Section S5), and in four bands if the shared factor has no memory. Most of the stationary shortfall is thus trend: at 16 to 255 trees the census is 3.0 to 7.7 times the model without noise at stationarity, and 1.49 to 1.92 times that model started from 1982. A species’ mean squared change is the square of its mean change per interval, the trend, plus the variance of its changes about that mean. For the 47 species of mean abundance 16 to 255 trees the trend is 47% of the census’s mean squared change, and the census’s trend term is 0.97 times the model’s under the regional pool and 1.00 times under the trade-off scheme.

This agreement of the trends is in sample, because the matrices are estimated from all seven intervals. The fluctuation about the trend of the same 47 species is still 1.43 and 1.45 times the model’s mean under the two schemes, inside its range. The first interval is 3.4 years and is run as one five-year step; over the other five intervals the ratios from 16 trees upward are 1.11 to 1.64.

The stationary curve of Fig. S9b does not average trajectories over starting states before the change is squared: it is computed within single realizations, with each species at each step in the band of its own count, so every sample of seven censuses starts from one state of the stationary model and keeps that state’s deviation from the stationary mean. The two panels differ because the census forest is not such a state. In 1982, 39 of the 84 species have a large-tree count outside the central 95% of their own stationary distribution under the trade-off scheme (64 under the regional pool). The count is 32 to 39 (60 to 64) at each of the seven censuses. Single states of the stationary model, each compared with the other 35 realizations, have 3.7 (2.7) such species on average, with a 97.5th percentile of 20 (16). This is a standing difference between the forest and the model’s stationary state, not an unusual fluctuation in 1982. The comparison cannot show whether the forest is away from equilibrium or the model’s equilibrium is wrong. Started from the census state, the model reproduces the directed change of the census: for the 47 species of mean abundance 16 to 255 trees, the mean path over realizations from 1982 accounts for 80% of the excess of that model’s mean squared change over that of stationary samples of equal abundance (79% under the regional pool), the rest being larger variability about the path, and the census’s trend term lies at the 98th percentile of the stationary samples (97th).

With the environmental noise on, the stationary rank order of species remains that of Eq. (S3). In the runs of Table S26 the rank correlation of the stationary abundances with the census changes by +0.004 under the regional pool and by −0.013 under the trade-off scheme, at most 0.02 in size. The leading species’ share of the large-tree stand changes by at most 2.0 percentage points. We have no analytical argument that the noise preserves the order in general, so this result rests on the simulations at the estimated amplitude.

The long-run values of the main text (Long-run composition and Fig. 4) come from longer simulations with the noise on, with the settings of the runs above. There are 36 replicates from each of two sets of initial conditions, run for 60 windows of 400 steps (120,000 model years). The stationarity test of Section S4 is met from window 9 (18,000 years) under the trade-off scheme and from window 6 under the regional pool. The abundances are means over the last five windows with the two sets pooled, and Table S10 gives their fit to the census beside that of the closed form. Sections S4 and S6 give the closed form’s values or those of the simulation with the noise off, except where they label a value as taken with the noise on. A shorter set of runs of the trade-off scheme (36 replicates, 30 windows) gave a rank correlation, an $R^2$, a large-tree total and a seedling pool within 1.2 standard errors of the values in Table S10 (1.6 under the regional pool). The trade-off scheme’s large-tree total at the end of the longer runs is 0.9% above its value in the same runs at the length of the shorter ones.

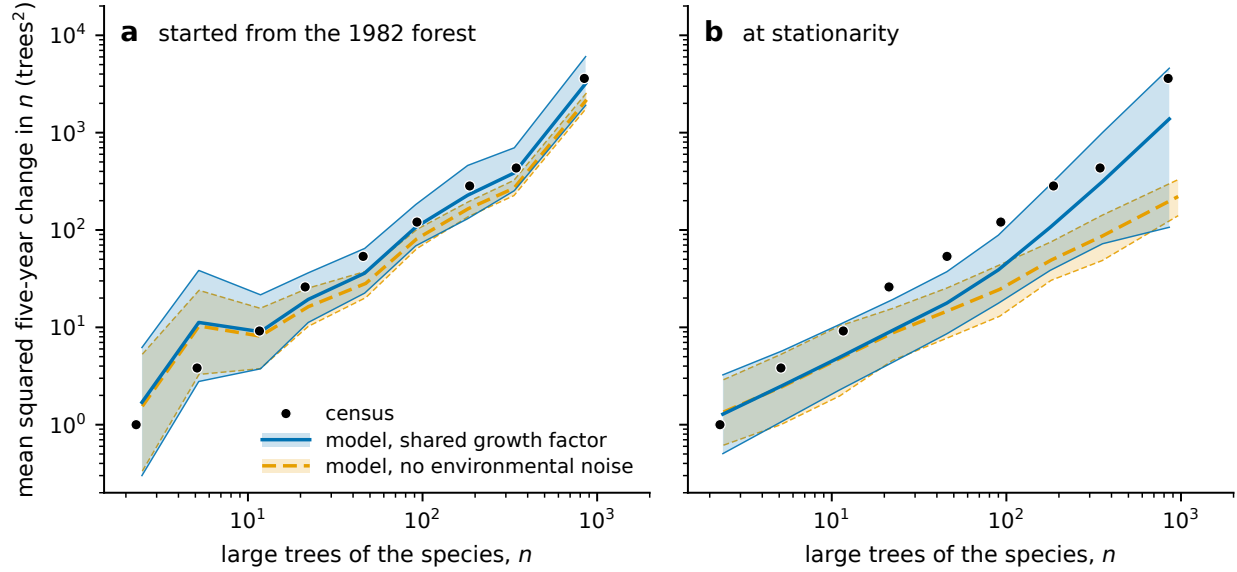

Figure S9: **Started from the 1982 forest, the model with the shared growth factor has the census's fluctuations inside its range in every abundance band.** Mean squared five-year change in a species' large trees (in trees squared) against its number of large trees at the start of the interval, in bands of doubling abundance, under the trade-off scheme. Points: the census, 1982 to 2010. Lines: the model's mean with the shared sapling-growth factor (solid) and without environmental noise (dashed). Shading with thin lines at its limits: the central 95% range of the model. (a) The model started from each species' 1982 stage vector and run over the same six intervals, with the range taken over 1,000 realizations. (b) The model at stationarity, as in Table S26, with the range taken over samples of seven censuses (six intervals) of single realizations; the census forest is not a typical state of the stationary model at any census (see text), so the samples in (b) lack most of the directed change present in (a). The lowest band, in which the census records no change, is not drawn, and the three bands below 16 trees hold 6 to 17 species-intervals each. The first census interval is 3.4 years. The text gives the ratios under both schemes.

The noise leaves the rank order unchanged to within 0.019 of the same simulations with the noise off. Under the regional pool it also leaves the size of the stand nearly unchanged, with the large-tree total at 0.95 of its value with the noise off. The rank correlation with the census changes by +0.002 and the identity-line $R^2$ on log abundance by +0.032. Under the trade-off scheme the noise lowers every censused class by a similar factor, about a third, so the distribution of stems over the classes keeps its shape (Fig. S3). The abundance with the noise on is 0.60 to 0.71 of the closed form’s in classes 2 to 8, against 0.93 to 1.09 under the regional pool. The large-tree total falls to 0.68 of its value with the noise off (18,204 against 26,858 large trees), which is the reduction described above. The rank correlation with the census changes by $-0.018$ and the $R^2$ by $-0.145$ (from +0.507 to +0.362), and the abundances have rank correlation 0.994 with those of the closed form. The statement of Section S5 that the shared factor acts without resizing or reordering the stand therefore holds for the order under both schemes, and for the size under the regional pool.

We tested whether the reduction of the stand under the trade-off scheme depends on the arrival rate $M$ or on the normalization of every matrix to growth rate one (Table S27). These runs have the noise on and the settings of the runs above, with 36 replicates of 12000 steps for each row of the table and means taken over the last five windows. They start from one set of initial conditions, so the test of Section S4 that compares two sets was not applied. Its test of drift from one window to the next is met in every row, although the large-tree total still drifts by up to 6.5% across the last five windows. Before the runs we fixed three outcomes that would count against the reduction. They were that the censused classes fill 0.9 of the capacity or more at four times $M$, that a net growth rate of 1% per interval holds them above 0.95, and that the rank correlation with the census moves by more than 0.05. Only the second occurred. At four times $M$ the classes fill 0.899 of the capacity, below 0.9.

A larger arrival rate brings more candidates for each vacancy, and the fraction of the capacity that is filled and the large-tree total rise with it. The identity-line $R^2$ is higher at four times $M$ than at half $M$, and the rank correlation with the census moves by at most 0.006. The reduction is thus a shortage of candidates, which a larger arrival rate makes up. A larger $M$, however, departs from its estimate from the immigrant fraction (Section S6). With the noise off the closed form’s censused abundances are independent of $M$, and only its seedling class moves with it (Section S4).

For a net growth rate we rescaled every matrix to growth rate $1 + \epsilon$ and left the trade-off weights unchanged. At each $\epsilon$ in the table the candidates exceed the vacancies and the censused classes are full, but the large-tree total is 3.6 to 3.8 times the observed, and the rank correlation with the census again moves by at most 0.017. Recomputing the trade-off weights at the rescaled gains leaves the total at 3.6 times the observed at the smallest rate. A net growth rate thus fills the stand only with more large trees than are observed, and we keep growth rate one in the model.

The environmental noise also widens the prediction intervals of the species-level forecasts. For the 75 species of the test of neutral drift (Section S3) we projected each species’ 1990 stage vector to 2015 with its matrix from the 1982–1990 tree tables, in 2,000 simulations with the noise at its estimated amplitude and common to all species. The central 90% of the simulated changes covers

Table S26: **Fluctuations of recruitment and of large-tree numbers in the model and in the census.** Stationary simulations under the two immigration schemes. Upper rows: the environmental noise at $\sigma_g = 0.32$ and at the two ends of its 90% confidence interval, all at $\tau_g = 18$ yr, 36 replicates per row. Lower rows: the mortality term of the text, alone and with the environmental noise at $\sigma_g = 0.32$ (36 replicates each). Statistics are taken over seven-step windows of the last 2,000 steps. CV, class 5: coefficient of variation of recruitment into class 5, the first large-tree class, over seven censuses (median over windows, with the fifth to ninety-fifth percentile at $\sigma_g = 0.32$). Inside: number of the ten abundance bands in which the census's mean squared five-year change lies inside the model's range over windows. Total: median absolute five-year relative change of the large-tree total. Range: thirty-year range ratio of that total. Stand: mean large-tree total over its mean in the same scheme with the noise and the mortality term off. Top bands: the census's mean squared five-year change over the model's in the two most abundant bands, the lower band first. Mid bands: the range of that ratio over the four bands of species with 16 to 255 large trees. Synchr.: synchronous fraction (%) of the most abundant band's squared five-year change, for the census before detrending. $\Delta\rho$: change in the rank correlation of the stationary abundances with the census means, from the same scheme with both terms off. In the upper rows the range ratio, the mid-band ratio and the synchronous fraction are given at the estimated amplitude only. The row marked census gives the observed values. Last row: the range accepted for each statistic. There $\times 2$ is a factor of two either side of the census value (of one, for the top bands), the stand may depart from one by the percentage shown, and the size of $\Delta\rho$ may not exceed the value shown.

| scheme | $\sigma_g$ | CV, class 5 | inside | total | range | stand | top bands | mid bands | synchr. | $\Delta\rho$ |
| --- | --- | --- | --- | --- | --- | --- | --- | --- | --- | --- |
| regional pool | 0.17 | 0.11 | 2 | 0.009 | – | 0.99 | 2.99, 4.36 | – | – | –0.001 |
|  | 0.32 | 0.21 (0.10–0.45) | 5 | 0.017 | 1.092 | 0.95 | 1.61, 1.77 | 2.9–3.5 | 33 | +0.004 |
|  | 0.84 | 0.55 | 9 | 0.041 | – | 0.76 | 0.40, 0.47 | – | – | –0.008 |
| trade-off | 0.17 | 0.11 | 3 | 0.011 | – | 0.84 | 2.77, 5.69 | – | – | –0.006 |
|  | 0.32 | 0.21 (0.10–0.45) | 6 | 0.020 | 1.109 | 0.66 | 1.37, 2.61 | 2.7–3.1 | 60 | –0.013 |
|  | 0.84 | 0.56 | 9 | 0.043 | – | 0.54 | 0.37, 0.47 | – | – | –0.065 |
| regional pool, mortality | off | 0.02 | 2 | 0.0057 | 1.023 | 0.98 | 2.60, 2.97 | – | 7 | –0.003 |
|  | 0.32 | 0.21 | 5 | 0.0176 | 1.098 | 0.94 | 1.26, 1.24 | 2.6–2.9 | 26 | +0.001 |
| trade-off, mortality | off | 0.03 | 4 | 0.0068 | 1.029 | 1.11 | 1.65, 1.84 | – | 7 | –0.037 |
|  | 0.32 | 0.21 | 6 | 0.0172 | 1.096 | 1.01 | 0.92, 1.16 | 1.9–2.7 | 20 | –0.031 |
| census |  | 0.18 |  | 0.0127 | 1.062 |  |  |  | 7.5 |  |
| accepted range | | 0.14–0.22 | $\geq 7$ | $\times 2$ | $< 1.10$ | 20% | $\times 2$ | | | 0.05 |

60% of the observed changes, against 27% with demographic noise alone and 96% at the upper end of the amplitude's confidence interval. The noise thus accounts for part of the spread that the forecasts lack, without bringing their coverage to the nominal 90%. These forecasts are also biased upward, mainly because their matrices come from the two intervals of strongest sapling growth. Sapling promotion rates over the forecast window were 0.58 to 0.79 of their 1982–1990 values. Rescaling the class 2 to 4 promotions by those ratios removes 92% of the mean bias and changes the rank correlation little. This rescaling uses conditions realized inside the forecast window.

Table S27: **The trade-off scheme with the noise on, at other arrival rates and with a net growth rate.** Upper rows: the arrival rate at half, one, two and four times its estimate $M$. Lower rows: every matrix rescaled to growth rate $1 + \epsilon$, with $\epsilon$ in percent per interval. Filled: the stems in the censused classes (2 to 8) as a fraction of the capacity. Candidates: candidates per vacancy. Large trees, seedlings and $R^2$ as in Table S10. For the arrival rate, $R^2$ is given at the two ends of the range.

| | filled | candidates | large trees | seedlings | $R^2$ |
| --- | --- | --- | --- | --- | --- |
| arrival rate $M/2$ | 0.53 | 0.17 | 0.69 | 1.7 | +0.19 |
| arrival rate $M$ | 0.65 | 0.25 | 0.81 | 1.6 | – |
| arrival rate $2M$ | 0.80 | 0.43 | 1.00 | 1.9 | – |
| arrival rate $4M$ | 0.90 | 0.67 | 1.12 | 2.1 | +0.48 |
| $\epsilon = 0.5\%$ | 1.00 | 1.6 | 3.6 | 2.1 | –1.6 |
| $\epsilon = 1\%$ | 1.00 | 2.3 | 3.7 | 2.9 | –2.6 |
| $\epsilon = 2\%$ | 1.00 | 4.3 | 3.8 | 5.5 | –3.8 |
